# Cross-modality representation and multi-sample integration of spatially resolved omics data

**DOI:** 10.1101/2024.06.10.598155

**Authors:** Zhen Li, Xuejian Cui, Xiaoyang Chen, Zijing Gao, Yuyao Liu, Yan Pan, Shengquan Chen, Hairong Lv, Lei Zhai, Rui Jiang

## Abstract

Spatially resolved sequencing technologies have revolutionized our understanding of biological regulatory processes within the microenvironment by accessing the states of genomic regions, genes and proteins as well as spatial coordinates of cells. However, discrepancies between different modalities and samples hinder the analysis of spatial omics data, necessitating the development of advanced computational methods. In this article, we propose PRESENT, an effective and scalable contrastive learning framework, for the cross-modality representation and multi-sample integration of spatial multi-omics, epigenomics and transcriptomics data. Through comprehensive experiments on spatial datasets, PRESENT demonstrates superior performance across various species, tissues, and technologies. Specifically, PRESENT effectively integrates spatial dependency and omics information simultaneously, facilitating the detection of spatially functional domains and the exploration of biological regulatory mechanisms. Furthermore, PRESENT can be extended for the integrative analysis of tissue samples across different dissected regions or developmental stages, promoting the identification of hierarchical structures from systematic and spatiotemporal perspectives.

## Introduction

Single-cell sequencing technologies have enabled the characterization of states and activities within individual cells, promoting molecular cell biology and disease research in areas such as cell atlas production, tumor immunology, and cancer genetics^1^. However, an indispensable step of single-cell sequencing techniques involves the dissociation of cells from the tissue of interest, resulting in the loss of spatial information^2^. Complex biological regulatory processes and cellular activities within the microenvironment are driven by intricate cell-cell interactions mediated by ligand-receptor partners, which heavily depend on the spatial distance between cells^3-5^. Recently, rapid advancements in spatially resolved sequencing technologies have enabled the simultaneous profiling of both spatial and omics data, such as the spatially resolved transcriptomics (SRT)^6, 7^, spatial epigenomics assay for transposase-accessible chromatin sequencing (ATAC-seq)^8, 9^ and spatial multi-omics such as spatial CITE-seq^10^, SPOTS^11^ and MISAR-seq^12^. These technologies have accumulated substantial spatially resolved omics data in repositories^9, 12^ and databases^13, 14^, thereby granting us a new perspective to decipher biological regulatory mechanisms in the context of microenvironment.

However, spatial coordinate data and omics data exhibit significant modality discrepancies, posing substantial challenges for the analysis of spatial omics data. Specifically, spatial coordinate data anchors the relative positions of cells within tissues, while omics data depicts the states of chromatin regions, genes and proteins involved in the gene regulatory processes within cells. Additionally, different layers of omics data further present distinct patterns and distributions^15-18^. For example, compared with RNA-seq, ATAC-seq datasets suffer from larger dimensionality, more severe sparsity and higher dropout rates, while proteomics datasets, sequenced by antibody-derived tags (ADT), demonstrate opposite characteristics, requiring targeted modeling for each omics layer. Cross-modality representation, a critical step in spatial omics data analysis, addresses these challenges by integrating multi-modal data from individual spatial datasets into a low-dimensional embedding matrix. This approach generates efficient and informative features for various downstream applications, including the detection of spatial domains, identification of domain-specific markers, and elucidation of domain-related functions, thereby facilitating the understanding of biological regulatory mechanisms within the microenvironment^19-21^.

Due to technical limitations, such as size restrictions of the captured area during data acquisition, individual spatial sequencing samples typically focus on a specific dissected region or developmental stage^22, 23^. Consequently, discrepancies in dissection and placement prevent the direct comparison of spatial coordinates between different samples^23^. Additionally, the intricate coupling between batch effects and biological variations hinders the applications of single-sample representation methods to the joint embedding of multiple samples, impeding the exploration of tissues under different conditions^23-25^. Therefore, the integrative analysis of multiple samples, known as multi-sample integration, is another critical focus in the research of spatially resolved omics data. While facing greater challenges compared to representation of individual samples, multi-sample integration offers a more comprehensive and systematic perspective for deciphering the hierarchical structures of cellular organizations^22, 23, 26, 27^. For example, literatures have demonstrated that integrated analysis of samples under different stages reveals developmental dynamics in the early mouse embryo data^27, 28^. Moreover, recent studies have confirmed that integrating disease and normal samples facilitates the identification of disease-specific domains and promotes the investigation of disease onset and progression mechanisms^27, 29^.

Many computational methods have been proposed to address the integrative analysis of SRT data^30^, including single-sample representation algorithms like SpaGCN^31^, STAGATE^32^ and PAST^19^, as well as multi-sample integration methods like STAligner^27^, Stitch3D^33^ and GraphST^22^. Despite the achievements, the development of algorithms for spatial ATAC-seq^8, 9^ and spatial multi-omics data^11, 12, 34^ remains in preliminary stage. Recently, SpatialGlue^35^ was introduced as the first attempt for the single-sample representation of spatial multi-omics data using an attentional feature fusion mechanism. Tian et al. proposed a dependency-aware generative variational autoencoder framework for the single-sample representation of SRT, spatial ATAC-seq and spatial CITE-seq data^36^. While these studies have addressed the cross-modality representation of individual spatial epigenomics and multi-omics samples, they have overlooked the integrated analysis of multiple samples across different conditions or time points, impeding the understanding of tissue structures from a spatiotemporal perspective. To address these challenges, there is an urgent need for advanced computational algorithms capable of integrating and analyzing cross-modality and multi-sample spatially resolved omics data.

To this end, we propose PRESENT, an advanced and scalable contrastive learning framework for the cross-modality representation and multi-sample integration of s**P**atially **RE**solved omics data, including spatial multi-omic**S, E**pigenomics, tra**N**scrip**T**omics (Fig. 1). Through comprehensive experiments on datasets generated by various spatial sequencing technologies, including spatial multi-omics^11, 12, 34^, epigenomics^8, 9^, and transcriptomics^6, 7^, we demonstrated that PRESENT can simultaneously capture spatially coherent patterns and complementary multi-omics information, obtaining interpretable representations for various downstream analyses, including spatial visualization and the detection of spatial domains. Based on the Bayesian neural networks, PRESENT also offers the potential to incorporate reference data generated by single-cell or spatial technologies, addressing issues including low sequencing depth and signal-to-noise ratio associated with spatial data. Furthermore, PRESENT can be extended for the integrative analysis of multiple samples across different dissection regions and developmental stages, effectively eliminating batch effects while maintaining shared and sample-specific biological variations, paving the way to explore the hierarchical tissue structures and sophisticated cellular activities from systematic and spatiotemporal perspectives.

**Fig. 1.**
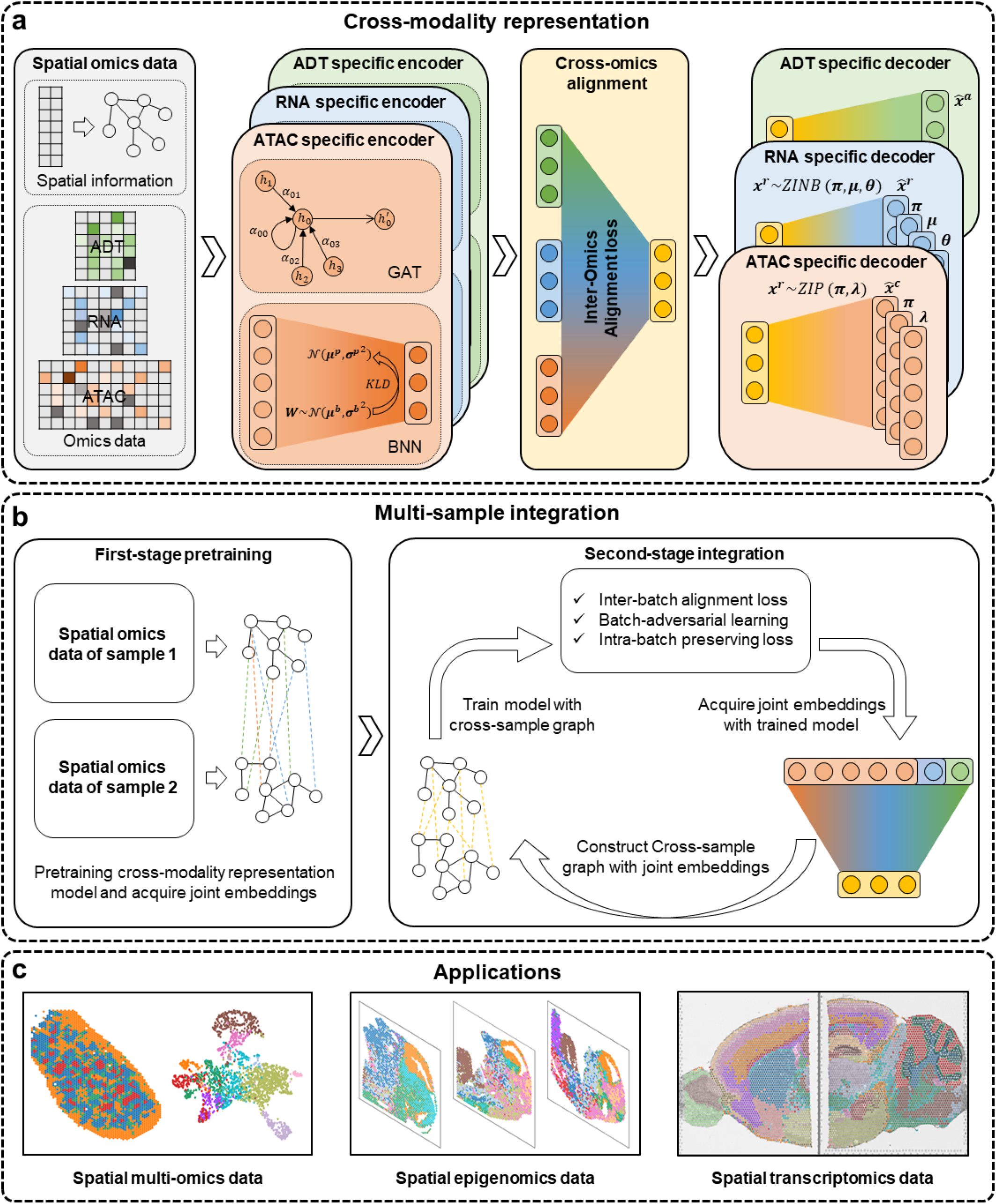
The framework of PRESENT. **a**, Cross-modality representation of spatial multi-omics data. PRESENT is capable to integrate spatial coordinate data and multi-omics data, including proteomic data sequenced by antibody-derived tags (ADT), transcriptomic data by RNA sequencing (RNA), and epigenomic data generated by the assay for transposase-accessible chromatin with sequencing (ATAC). Firstly, an omics-specific encoder is used to extract specific biological variations contained in each omics layer. Each omics-specific encoder is composed of a graph attention neural network (GAT) and a Bayesian neural network (BNN) module. All the omics-specific feature matrices extracted by encoders are fused by a cross-omics alignment module, which integrates multi-omics information via the contrastive learning inter-omics alignment loss function. The output of cross-omics alignment module refers to the final representation matrix of PRESENT. The distinct patterns of different omics layers are addressed via omics-specific decoders, which take the output of cross-omics alignment module as input and model the count matrices of RNA and ATAC data using the zero-inflation negative binomial (ZINB) and the zero-inflation Poisson (ZIP) distribution, respectively. **b**, Multi-sample integration of spatial omics data. PRESENT adopts a two-stage workflow for the integration of multiple spatial samples. In the first pretraining stage, PRESENT is pretrained using omics data and the cross-sample graph obtained from all the samples. The intra-sample edges of the cross-sample graph are built using spatial coordinates, while inter-sample connections are established based on the similarity of omics features across samples. In the second integration stage, PRESENT implements a cyclic iterative strategy to refine the cross-sample graph using the joint embeddings obtained by the trained model. PRESENT further introduces a contrastive learning-based inter-batch alignment loss, a batch-adversarial learning strategy and an intra-batch preserving loss in the second stage to eliminate batch effects while preserving biological signals. **c**, Applications of PRESENT in spatial omics data. PRESENT can be easily adapted from spatial multi-omics to spatial single-omics data by simplifying the multi-view autoencoder into a single-view autoencoder. The joint embeddings obtained by PRESENT can be applied to various downstream analyses, including spatial visualization, domain identification and the integrative analysis of samples across conditions and developmental stages.

## Results

### Overview of PRESENT

Using a spatial multi-omics dataset that includes spatial coordinates, epigenomic profiles, transcriptomic expressions, and proteomic profiles, we illustrate the cross-modality representation architecture of PRESENT in Fig. 1a. Key challenges in the cross-modality representation of spatial multi-omics data include recognizing patterns within individual modalities and integrating multi-omics biological signals. Built on a multi-view autoencoder, PRESENT recognizes spatially biological variations contained in each omics layer via an omics-specific encoder consisting of graph attention neural networks^37^ (GAT) and a Bayesian neural network^38^ (BNN). The GAT module aims to incorporates spatial dependencies and omics information, while the BNN module, known for its effectiveness in non-linear regression problems^38^, brings the potential to leverage reference data from existing sequencing data^19^. To further capture the distinct patterns of various omics layers, PRESENT employs omics-specific decoders which model the counts of transcriptomic and epigenomic data via zero-inflation negative binomial (ZINB) distribution^39^ and zero-inflation Poisson (ZIP) distribution^40^ (Methods), respectively. Moreover, to facilitate the integration of multi-omics biological signals, PRESENT fuses omics-specific embeddings from the encoders using a cross-omics alignment module which implements a multi-modal contrastive learning loss function, namely the inter-omics alignment (IOA) loss^41^ (Methods).

Based on the above cross-modality representation framework, PRESENT can be further extended to the multi-sample integration of spatial multi-omics data, as illustrated in Fig. 1b. Critical challenges in multi-sample integration include addressing the spatial coordinate discrepancies between samples to capture cross-sample relationships^23^ and distinguishing complexly coupled biological variations from technical batch effects^22, 26, 27, 33^. PRESENT adopts a two-stage training workflow to address these issues. In the first stage, PRESENT is pretrained based on the cross-sample graph and omics data acquired from all the input samples. Specifically, the intra-sample edges of the cross-sample graph are established using the spatial coordinates, while the inter-sample connections are built using omics information (Methods). In the second stage, to align spots from multiple samples and eliminate technical batch effects from the coupled biological variations, we introduce a contrastive learning strategy^42^ and a batch-adversarial learning strategy^43^. Additionally, we incorporate an intra-batch preserving loss during the training process to maintain biological signals within each sample while integrating multiple samples. Moreover, the second stage employs a cyclic iterative optimization process to refine the cross-sample graph using the joint embeddings generated by the newly trained model at each epoch, enabling a progressively improved characterization of cross-sample relationships (Methods).

On the other hand, PRESENT can be seamlessly adapted from spatial multi-omics data to spatial single-omics data, such as SRT and spatial epigenomics data, by simplifying the multi-view autoencoder framework into a single-view autoencoder (Methods). Through experiments on massive spatial datasets generated by spatial multi-omics, epigenomics and transcriptomics technologies, we have demonstrated that the joint embeddings obtained by PRESENT facilitate various downstream applications, including spatial visualization, domain detection, the integrative analysis of horizontal slices from different dissection regions, and vertical integration of samples from different developmental stages (Fig. 1c), paving the way to explore the cellular activities from spatiotemporal perspectives.

### PRESENT facilitates accurate spatial domain identification in spatial RNA-ADT data

Spatial domain identification, which aims to decipher tissue regions with distinct anatomical structures, cell-type compositions, and cell-cell interactions^44, 45^, is a fundamental step in revealing gene regulatory processes and tissue functions in spatial omics data^31, 32^. We first collected a spatial RNA-ADT human lymph node dataset generated by 10x Genomics Visium technology^35^. This dataset simultaneously accessed the spatial locations, transcriptomic and proteomic expression of spots and was manually annotated by the clinician group according to the histological image^35^ (Fig. 2a). We benchmarked PRESENT with SpatialGlue^35^ and spaMultiVAE^36^, two representation learning methods recently designed for spatial multi-omics data, and several methods developed for single-cell RNA-ADT paired sequencing data, including totalVI^46^, CiteFuse^47^ and Seurat-v4^48^. Based on the latent embeddings generated by these methods, we implemented the Leiden algorithm^49^ to acquire the same number of clusters as the ground truth domains (Methods). Four commonly used metrics^50, 51^, including adjusted rand index (ARI), adjusted mutual information (AMI), normalized mutual information (NMI) and homogeneity (Homo), were utilized to quantitatively evaluate the performance of spatial domain detection. As shown in Fig. 2b, PRESENT achieved the best performance on all the four metrics for spatial domain identification. As a qualitative evaluation and intuitive illustration of detected domains, we also demonstrated the uniform manifold approximation and projection (UMAP) manifold structure^52^ and spatial visualization results (Fig. 2d, e, f). Specifically, CiteFuse and Seurat-v4 obtained unsatisfactory visualization results, where domains such as medulla cords and sinuses scattered in the UMAP plot. spaMultiVAE detected stripe-like spatial domains, resulting in biased spatial pattern identification in the lymph node tissue. Although SpatialGlue and totalVI achieved relatively better results, they failed to recognize capsule from pericapsular adipose tissue. In contrast, PRESENT successfully distinguished the pericapsular adipose tissue (cluster 8), capsule (cluster 1), follicle (cluster 3) and cortex (cluster2 and cluster 10). Additionally, we applied PRESENT to only a single omics layer of the human lymph node dataset, thus leveraging expression information from only RNA data or ADT data, denoted as RNA-only and ADT-only, respectively. As shown in Fig. 2c, PRESENT utilizing multiple omics layers (RNA & ADT) achieved improved performance compared to the model that only leveraged RNA or ADT data, highlighting the capabilities of PRESENT in integrating cross-modality spatial multi-omics information.

**Fig. 2.**
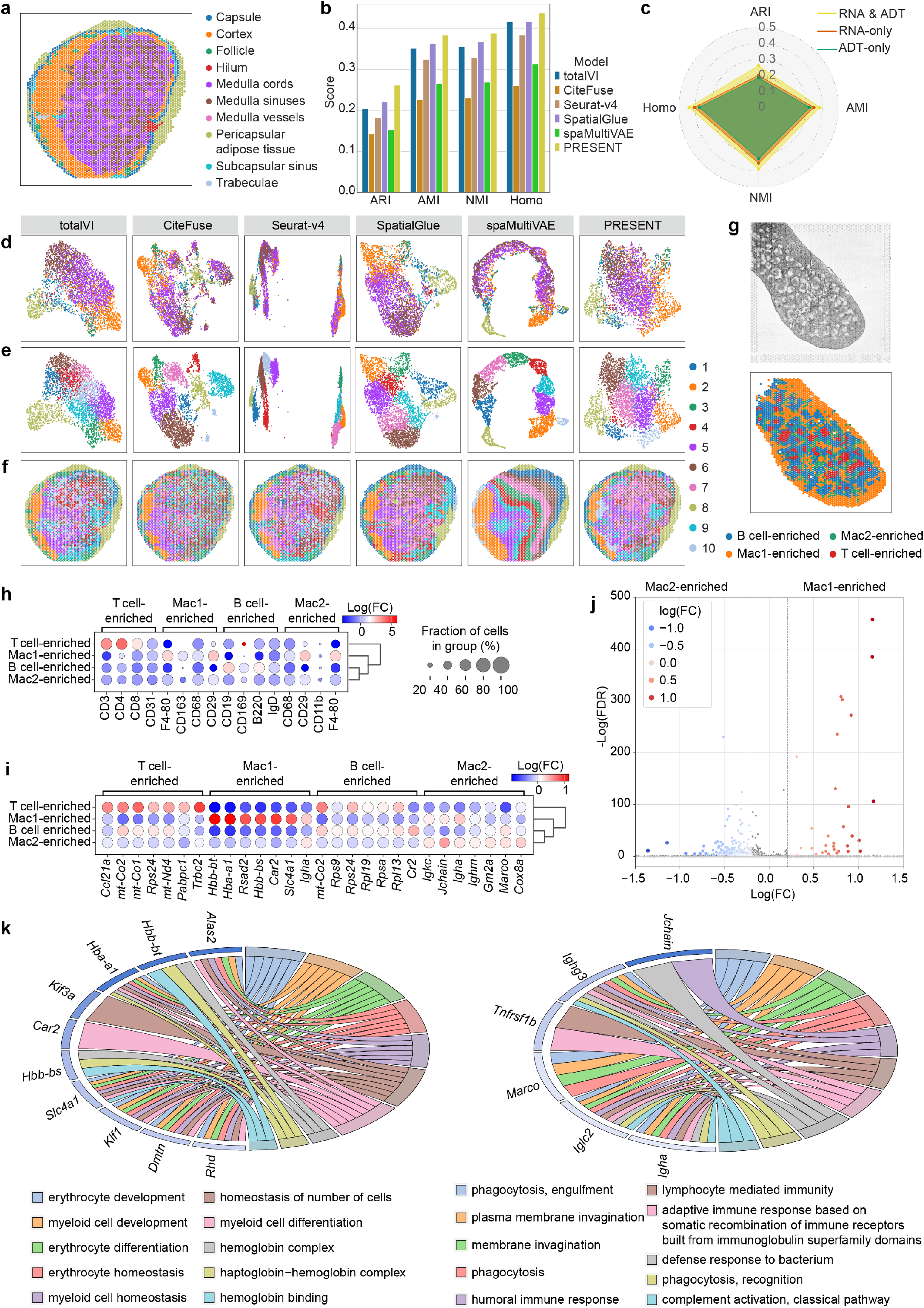
PRESENT facilitates accurate spatial domain identification in spatial RNA-ADT data. **a**, The spatial visualization of the 10x Genomics Visium RNA-Protein human lymph node sample colored by ground truth domain labels. **b**, The quantitative comparison of spatial domain identification performance between PRESENT and other baseline methods based on bar plot and, **c**, the comparison between PRESENT utilizing both RNA and ADT data (RNA & ADT) and PRESENT using only RNA (RNA-only) or ADT data (ADT-only) based on radar plot on the human lymph node dataset. **d**, The UMAP visualization of latent embeddings obtained by different methods colored by ground truth domain labels and **e**, cluster labels identified based on different methods on the human lymph node dataset. **f**, The spatial visualization of clusters identified based on different methods on the human lymph node dataset. **g**, The histology image and spatial visualization of spatial clusters identified based on PRESENT on the SPOTS mouse spleen dataset. **h**, The dot plot demonstrating the difference of ADT expression levels and **i**, RNA expression levels between each domain and the rest domains. **j**, The volcano plot illustrating the difference of RNA expression levels between Mac1 and Mac2-enriched domains. The vertical dashed line represents the threshold for log(F**C**) = ±0.2, while the horizontal dashed line denotes the threshold for −log(F**D**R = 0.05). **k**, The chordal plots of the Mac1-enriched domain (left) and the Mac2-enriched domain (right) demonstrating the linkage of differential expressed genes (DEGs) in each domain and the corresponding enriched biological processes. In each chordal plot, the left semicircle represents DEGs while the right semicircle denotes the enriched biological processes.

As the largest secondary lymphoid organ in the body, the spleen plays a significant role in hematopoiesis and the clearance of erythrocytes, contributing significantly to immune functions^53^. We further applied PRESENT to a SPOTS mouse spleen dataset^11^, which measured spatial transcriptomic and proteomic profiles simultaneously, to decipher the functionality of mouse spleen in the context of cellular organizations. Through cross-modality representation and spatial clustering, PRESENT identified four main spatial domains (Fig. 2g), denoted as the B cell-enriched domain, the T cell-enriched domain and two macrophage-enriched subdomains, i.e., Mac1-enriched and Mac2-enriched domains, according to our differentially expressed analysis (Fig. 2h, i). The differentially expressed analysis compared the protein expression level of each domain with those of the rest domains based on the Wilcoxon rank-sum test^54^ (Fig. 2h), identifying protein markers distinguishing the B cell-enriched domain (CD3, CD4, CD8), the T cell-enriched domain (CD19, B220, lgD) and the macrophage-enriched domain (F4-80, CD163, CD68), consistent with the original study^11^. Specifically, the Mac1-enriched and Mac2-enriched domains had nearly identical protein markers (Fig. 2h) as the macrophage markers identified in the original study^11^, making it hard to distinguish the two subdomains purely by ADT data. Therefore, we further conducted differentially expressed analyses on the transcriptomic data between domains identified based on PRESENT (Fig. 2i). Results demonstrated that Mac1-enriched domain could be marked by genes including *Hbb-bt, Hbb-a1, Rsad2, Hbb-bs, Car2* and *Slc4a1*, while Mac-2 enriched domain could be recognized with *Igkc, Jchain, Igha, Ighm, Gm2a* and *Cox8a*. The RNA expression levels between the Mac1 and Mac2-enriched domains were also directly compared and numerous significantly differentially expressed genes (DEGs) were found (Fig. 2j). These results showed that, despite the similarity of the two macrophage-enriched subdomains in ADT expression levels, the distinction of transcriptomic patterns between them could still be identified by PRESENT, emphasizing the ability of PRESENT in leveraging complementary proteomic and transcriptomic information for effective domain identification.

To further decipher the functional distinction between the two macrophage-enriched subdomains, we conducted Gene Ontology (GO) enrichment analysis based on the identified DEGs (Methods). Specifically, for the Mac1-enriched domain, we enriched biological processes including erythrocyte development, erythrocyte differentiation, erythrocyte homeostasis, myeloid cell development, myeloid cell differentiation and myeloid cell homeostasis with false discovery rates (FDRs) of 1.29e-8, 5.48e-6, 6.70e-6, 3.59e-7, 8.71e-5 and 2.06e-5, respectively (Fig. 2k and Supplementary Table 1). Macrophages are myeloid lineage cells which derive from bone marrow^55^, reasonably explaining the enrichment of development, differentiation and homeostasis of myeloid cell in the Mac1-enriched domain. In addition, previous studies have stressed the importance of macrophages in the entire life of erythrocytes by providing necessary factors for the survival, proliferation and development of erythrocytes^56, 57^, agreeing with the development, differentiation and homeostasis of erythrocyte found in the Mac1-enriched domain. In comparison, the Mac2-enriched domain found biological processes including phagocytosis & engulfment, plasma membrane invagination, membrane invagination, phagocytosis, humoral immune response and lymphocyte mediated immunity with FDRs of 7.11e-5, 7.11e-5, 7.11e-5, 4.34e-4, 5.58e-4 and 5.94e-4, respectively (Fig. 2k and Supplementary Table 2). Phagocytosis and other related processes, called the process of ‘cell eating’, are commonly considered as classical functions of macrophages^58, 59^, validating the enrichment of macrophages in the Mac2-enriched domain. On the other hand, compared with the Mac1-enriched domain, the Mac2-enriched domain was spatially close to the B cell-enriched and T cell-enriched domains (Fig. 2g), thereby explaining the enrichment of lymphocyte mediated immunity process^60^. In conclusion, PRESENT effectively integrates multi-omics information and spatially coherent patterns in spatial RNA-ADT data, thereby facilitating accurate spatial domain identification and systematic exploration of biological implications in the identified domains.

### PRESENT integrates spatially aware multi-omics biological signals in spatial RNA-ATAC data

Next, we collected MISAR-seq mouse brain datasets^12^, which simultaneously measured the epigenomics and transcriptomics, i.e., paired spatial RNA-ATAC profiles, of four mouse brain samples from four developmental stages, involving E11.0, E13.5, E15.5 and E18.5. Compared with spatial-RNA-ADT co-profiling data, spatial RNA-ATAC datasets suffer from higher dimension and sparsity. We firstly compared the performance of PRESENT in spatial domain identification with the spatial multi-omics method SpatialGlue^35^ and six methods for the representation of single-cell RNA-ATAC data, including MOFA+^61^, scAI^62^, scMDC^63^, scMVP^21^, Seurat-v4^48^ and SnapATAC2^64^. Here spaMultiVAE^36^, totalVI^46^ and CiteFuse^47^ were not used as they are specially designed for single-cell or spatial RNA-ADT data. As shown in Fig. 3a, PRESENT achieved overall the best spatial clustering performance, illustrating the its superiority in extracting spatial multi-omics information into effective cross-modality embeddings for accurate spatial domain identification. We also used PRESENT to obtain latent representations based on only ATAC data or RNA data, denoted as ATAC-only and RNA-only respectively, which led to decreased performance, highlighting the importance of integrating multi-omics information in deciphering biological variations in the context of cellular organizations (Fig. 3b).

**Fig. 3.**
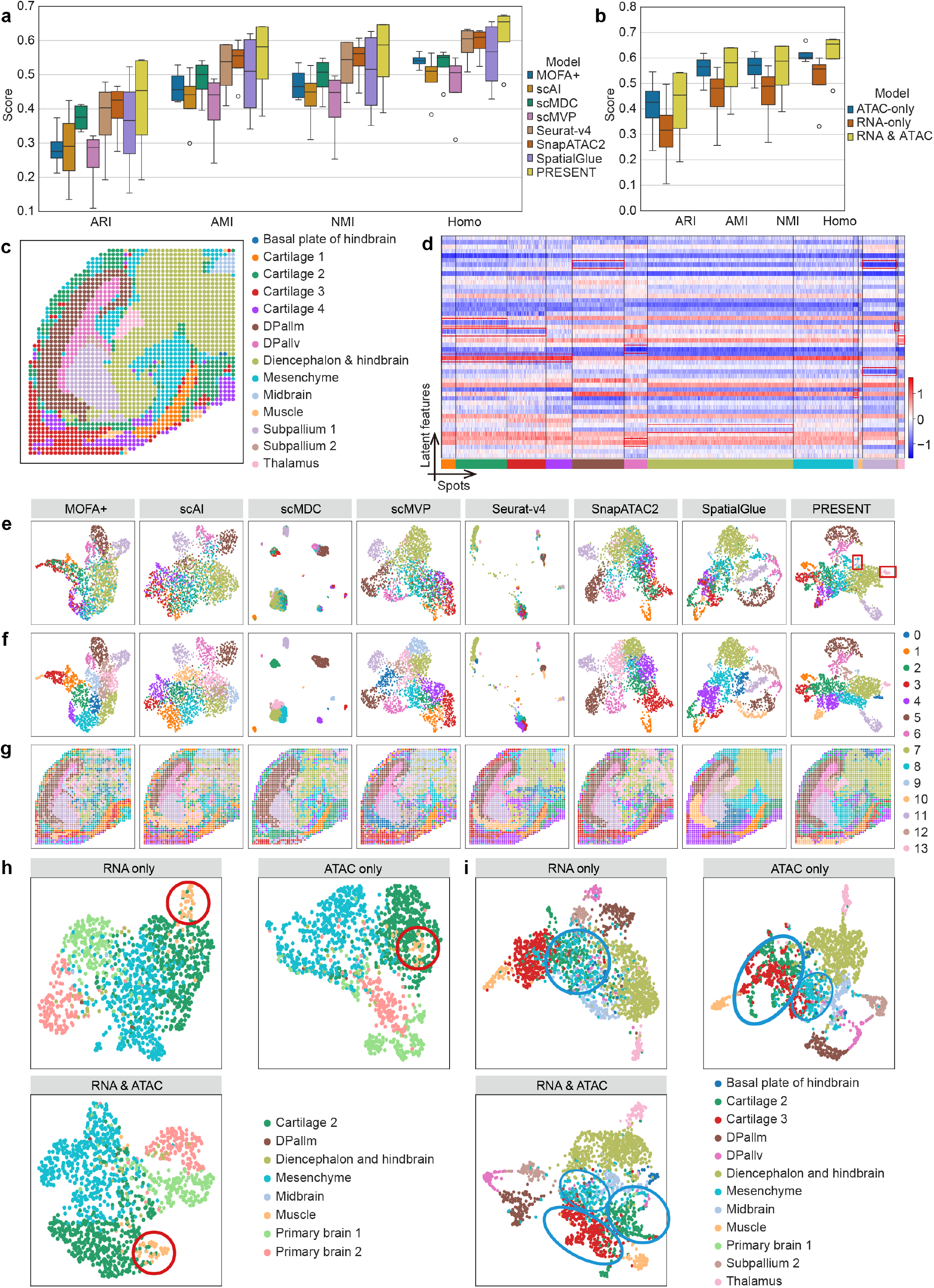
PRESENT integrates spatially aware multi-omics biological signals in spatial RNA-ATAC data. **a**, The quantitative comparison between PRESENT and other baseline methods and **b**, the comparison between PRESENT utilizing both RNA and ATAC data (RNA & ATAC) and PRESENT using only RNA (RNA-only) or ATAC (ATAC-only) data based on box plot on the MISAR-seq mouse brain dataset. The center line, box limits and whiskers in the box plots are the median, upper and lower quartiles, and 1.5× interquartile range, respectively. **c**, The spatial visualization of spots from the E18.5 mouse brain sample colored by ground truth domain labels. **d**, The heatmap visualization of latent representations obtained by PRESENT. **e**, The UMAP visualization of latent representations obtained by different methods colored by ground truth domain labels. **f**, The UMAP visualization of latent representations obtained by different methods and **g**, the spatial visualization colored by spatial clusters identified based on different methods on the E18.5 mouse brain sample. The UMAP visualization of latent embeddings obtained by PRESENT using only RNA data, only ATAC data and both RNA & ATAC data, colored by ground truth domain labels on the **h**, E11.0 and **i**, E15.5 mouse brain samples.

Taking the E18.5 mouse brain sample with the most complex structures as an example (Fig. 3c), we intuitively visualized the latent embeddings learned by PRESENT using heatmap (Fig. 3d). The spots from different spatial domains could be successfully distinguished via the combination of several latent features or even a single feature in the latent feature space, including large domains such as cartilage 1-4, mantle zone of dorsal pallium (DPallm), ventricular zone of dorsal pallium (DPallv), diencephalon & hindbrain and subpallium 1, and small domains like midbrain and thalamus. We further demonstrated the UMAP visualization of latent representations obtained by different methods on the E18.5 mouse brain sample (Fig. 3e, f). Most baseline methods, including MOFA+, scAI, scMDC, scMVP and Seurat-v4, could only roughly separate large domains such as DPallm, DPallv, diencephalon & hindbrain and subpallium 1. SnapATAC2 could additionally recognize the cartilage 1 but still failed to distinguish others. In comparison, spatially aware methods, including SpatialGlue and PRESENT, not only identified large spatial domains such as diencephalon & hindbrain, subpallium 1, DPallm, DPallv, cartilage 1-4, but also accurately distinguished the two smaller spatial domains, i.e., midbrain and thalamus. Furthermore, domains identified by these two methods demonstrated enhanced spatial smoothness, highlighting the importance of integrating spatial information (Fig. 3g). The UMAP and spatial visualization of other three samples of the MISAR-seq mouse brain data^12^ achieved similar results (Supplementary Figs. 1-4).

Additionally, we have conducted comprehensive experiments to investigate the impact of biological signals from different omics on model performance to provide deeper insights into the model′s ability to extract multi-omics biological signals. Specifically, PRESENT utilizing only the transcriptomic data of E11.0 brain sample, denoted as RNA-only, successfully captured the difference between the muscle and the cartilage 2 domain, while the model based solely on epigenomic data, depicted as ATAC-only, failed to do so (red circles in Fig. 3h). On the other hand, compared with the failure of PRESENT using only transcriptomic data (RNA-only) in the E15.5 brain sample, the model based on the epigenomic data (ATAC-only) contributed more to the differentiation between cartilage 2, cartilage 3 and mesenchyme domains (blue circles in Fig. 3i). Therefore, relying solely on a single omics layer might lead to a biased interpretation of biological signals in spatial domain detection. When taking both omics layers as input (RNA & ATAC), PRESENT successfully captured variations among those domains in both E11.0 and E15.5 mouse brain samples (Fig. 3h, i), highlighting the capabilities of PRESENT in leveraging spatially aware transcriptomic & epigenomic multi-omics information for cross-modality representation and the accurate identification of spatial domains from a comprehensive perspective.

### PRESENT eliminates batch effects while preserving biological signals in spatial multi-omics data

The MISAR-seq mouse brain dataset simultaneously measured the transcriptomic and epigenomic profiles of developmental brain samples from four stages, including E11.0, E13.5, E15.5 and E18.5, providing a multi-omics landscape of the developmental mouse brain in the context of cellular organizations^12^. However, varying experimental conditions and sequencing batches often introduce potential technical biases, resulting in significant discrepancies in omics data across samples. These discrepancies, commonly referred to as batch effects in single-cell studies, pose challenges to the integrative analysis of multiple samples from different conditions and developmental stages^24, 26^. For illustration, we applied joint principal component analysis (PCA) and the UMAP algorithm to raw RNA and ATAC data of the four MISAR-seq mouse brain samples, revealing pronounced batch effects (Fig. 4a). Moreover, during the development of mouse brain, several functional structures might be specific to certain developmental stages (e.g., primary brain), while others may persist throughout the entire developmental process (e.g., midbrain and muscle), as shown in Fig. 4b. Therefore, the key challenges in integrating the MISAR-seq mouse brain samples lied not only in the incorporation of spatial multi-omics information and the correction of batch effects across samples, but also in the conservation of shared and sample-specific biological variations.

**Fig. 4.**
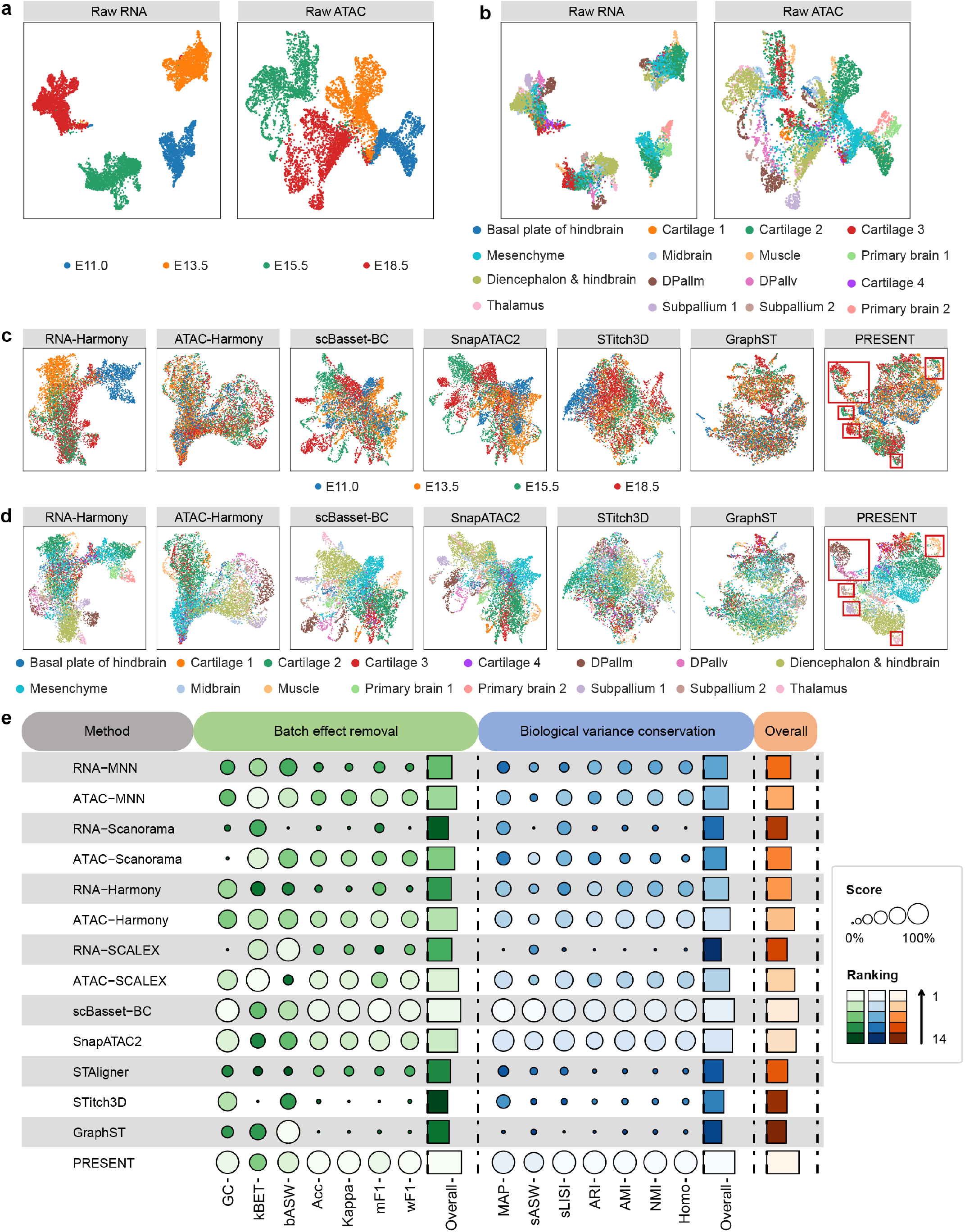
PRESENT eliminates batch effects while preserving biological signals in spatial multi-omics data. The UMAP visualization of spots across the four MISAR-seq mouse brain samples based on raw RNA or ATAC data colored by **a**, sample indices and **b**, ground truth domain labels. Note that prior to applying the UMAP algorithm to raw RNA or ATAC data, we first reduce the dimensionality to 50 using PCA transformation. **c**, The UMAP visualization of spots across the four MISAR-seq mouse brain samples based on the joint embeddings produced by different integration methods colored by **c**, sample indices and **d**, ground truth domain labels. **e**, The quantitative evaluation of different integration methods on the four MISAR-seq mouse brain samples using 14 metrics divided into two categories, namely batch effect removal and biological variance conservation. The category scores of these two aspects were calculated by averaging the metrics within each category. An overall score for each integration method was computed using a 40/60 weighted mean of the category scores for batch effect removal and biological variance conservation.

Due to the lack of multi-sample integration methods specially designed for spatial multi-omics data, we compared the integration performance of PRESENT with six methods developed for single-cell data (MNN^65^, Scanorama^25^, Harmony^24^, SCALEX^66^, scBasset-BC^67^ and SnapATAC2^64^) and three recently proposed methods designed for SRT data (STAligner^27^, Stitch3D^33^ and GraphST^22^). Specifically, MNN, Scanorama, Harmony and SCALEX could be applied to batch integration of either RNA-seq or ATAC-seq data, denoted as RNA-* and ATAC-*, respectively, where * represents these methods. scBasset-BC was featured as a sequence-based method specially designed for single-cell ATAC-seq data, while STAligner, Stitch3D and GraphST were intended for integrative analysis of SRT data. SnapATAC2 has provided guidance for incorporating multi-omics (i.e., transcriptomics & epigenomics) information for integrated analysis, serving as a multi-sample integration workflow for single-cell multi-omics.

We first applied the UMAP algorithm to the joint embeddings generated by PRESENT and other baseline methods for visualization, with the spots colored by batch indices (Fig. 4c and Supplementary Fig. 5a) and ground truth domain labels (Fig. 4d and Supplementary Fig. 5b). PRESENT successfully integrated spots from different developmental stages (red boxes in Fig. 4c) while maintaining the biologically distinct patterns of DPallm, DPallv, subpallium 1 & 2, thalamus and muscle (red boxes in Fig. 4d). Specifically, spots of muscle domain persisted throughout the entire life of mouse brain development, while those of subpallium 1&2, DPallm, DPallv and thalamus only existed in specific stages. The success of PRESENT in distinguishing these domains illustrated its capabilities in eliminating batch effects while capturing shared and stage-specific biological variance. Additionally, we conducted joint spatial clustering across the four stages based on the joint embeddings obtained by different methods, as shown in Supplementary Fig. 6. Except for the scBasset-BC and SnapATAC2, other baseline methods delivered limited clustering results. In contrast, PRESENT successfully detected not only large domains like DPallm (cluster 7), but also small domains including thalamus (cluster 16) and muscle (cluster 11). It is worth noting that when employing individual analysis of the sample from the E18.5 stage (Supplementary Fig. 4), PRESENT was unable to identify these small domains, underscoring the advantages of integrative analysis across multiple samples.

Furthermore, to intuitively evaluate the integration and biological conservation capabilities of PRESENT and other baseline methods, we systematically benchmarked them using 14 metrics, evenly divided into two categories: batch effect removal and biological variance conservation, following a recent benchmarking framework for single-cell omics data integration^68^ (Methods). As shown in Fig. 4e, scBasset-BC and SnapATAC2 provided satisfactory performance through incorporating sequence and multi-omics information, respectively. PRESENT, which simultaneously integrated spatial dependency and multi-omics information of all the samples, achieved the best performance in both batch effect removal and biological variance conservation among all the 14 methods, illustrating its superiority in multi-sample integration for spatial multi-omics data.

### PRESENT facilitates effective cross-modal representation in spatial single-omics data

PRESENT is highly extensive and can be seamlessly adapted from spatial multi-omics data to spatial single-omics data, such as spatial epigenomics and transcriptomics data, by simplifying the multi-view autoencoder into a single-view model (Methods). We first accessed the performance of PRESENT on spatial epigenomics data by collecting the spatial ATAC mouse embryo datasets which consist of three developmental stages^9^, including E15.5, E13.5 and E12.5. The datasets include six samples, with each developmental stage containing two replicated samples, denoted as *-S1 and *-S2. Initially, we evaluated the data quality of the six samples using logarithmic total fragment counts, denoted as log10(nFrags), and transcription starting site (TSS) enrichment scores. The former metric represents the cellular sequencing depth of samples^69^, while the latter reflects the signal-to-noise ratio of datasets^70^. As shown in Fig. 5a, b, the samples of E15.5 stage, i.e., E15.5-S1 and E15.5-S2, achieved the highest sequencing depth and signal-to-noise ratio, while the E13.5-S1 sample and E12.5-S2 sample suffered from the lowest sequencing depth and signal-to-noise ratio, respectively.

**Fig. 5.**
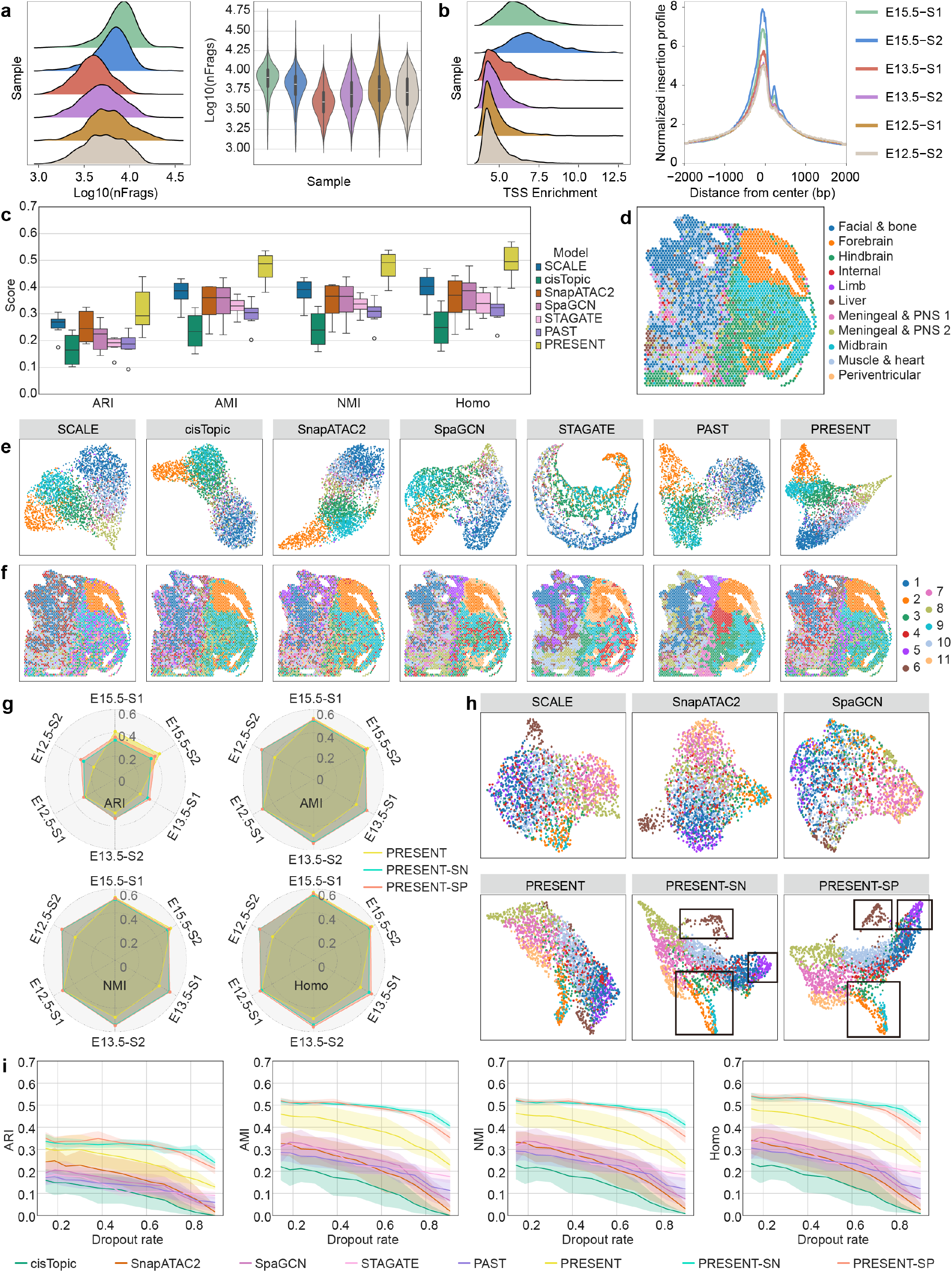
PRESENT facilitates effective cross-modal representation in spatial single-omics data. **a**, The ridge plot and violin plot of log10 total fragment counts, namely log10(nFrags), across the six spatial ATAC mouse embryo samples. **b**, The ridge plot of TSS enrichment scores for spots and the line plot of the enrichment profiles of fragments around the TSS across the six mouse embryo samples. **c**, The quantitative evaluation of spatial clustering performance of different methods using four metrics, including ARI, AMI, NMI and Homo. The center line, box limits and whiskers in the box and violin plots are the median, upper and lower quartiles, and 1.5× interquartile range, respectively. **d**, The spatial visualization colored by ground truth domain labels, **e**, the UMAP visualization of the embeddings obtained by different methods colored by ground truth domain labels and **f**, the spatial visualization of clusters identified using the embeddings obtained by different methods on the E15.5-S1 mouse embryo sample. **g**, The detailed comparison of performance of PRESENT and PRESENT-SN & SP on the six mouse embryo samples based on radar plots. **h**, The UMAP visualization of the latent representations obtained by different methods colored by ground truth domain labels on E12.5-S2 mouse embryo sample. **i**, The spatial clustering performance of baseline methods, PRESENT and PRESENT-SN & SP on the six mouse embryo samples under different levels of dropout rate. Here SCALE encountered error when the dropout rate was larger than 0.2.

For spatial domain identification, we benchmarked PRESENT and other six baseline methods, including three methods for single-cell ATAC-seq data (SCALE^71^, cisTopic^72^ and SnapATAC2^64^) and three methods for SRT data (SpaGCN^31^, STAGATE^32^ and PAST^19^). As shown in Fig. 5c, PRESENT achieved the best performance, with improvements of more than 8.60%, 26.13%, 25.91%, and 22.94% compared to the second-best baseline method across the clustering metrics, respectively. Using the E15.5-S1 sample as an example, we further demonstrated the UMAP and spatial visualization of spots colored by ground truth domains (Fig. 5d, e) and the spatial clusters identified based on different methods (Fig. 5f). Baseline methods like cisTopic and STAGATE could only identify the forebrain domain. Although SCALE, SnapATAC2, SpaGCN and PAST roughly recognized the subdomains of central nervous system, i.e., forebrain, hindbrain and midbrain, the low-dimensional representations acquired by them are decentralized and scattered, impeding their performance for spatial clustering (Fig. 5c, f). In contrast, PRESENT effectively distinguished domains including the forebrain, hindbrain, midbrain, facial & bone, muscle & heart, and meningeal & peripheral nervous system (PNS) 2. Visualization of the remaining five samples obtained similar results (Supplementary Figs. 7-13), demonstrating the capabilities of PRESENT in cross-modal representation of spatial epigenomics data.

Despite the superiority of PRESENT over existing baseline methods, low sequencing depth and signal-to-noise ratio stills constrained the effectiveness of the model, as illustrated in Fig. 5a, b and Supplementary Fig. 13. A previous study has shown that integrating external reference data through BNN module can effectively address issues related to high noise levels in SRT data^19^. Inspired by this, we collected a single-nucleus ATAC-seq mouse embryo dataset involving cells from three developmental stages, including E12, E13 and E15, which was sequenced in conjunction with the spatial ATAC mouse embryo data^9^. We utilized the single-nucleus ATAC-seq dataset (denoted as PRESENT-SN) and the other five spatial ATAC samples (denoted as PRESENT-SP) as the reference data for PRESENT when analyzing each spatial ATAC mouse embryo sample (Methods). As shown in Fig. 5g and Supplementary Fig. 14, the introduction of prior information from reference data facilitated enhanced performance of PRESENT, especially in samples with lower sequencing depth and signal-to-noise ratio, such as the E13.5-S1 and E12.5-S2 mouse embryo samples. This finding was further validated by the UMAP visualization of the two low-quality samples (Fig. 5h and Supplementary Figs. 9 and 12). Specifically, PRESENT-SN and PRESENT-SP successfully identified spatial domains including liver, limb, forebrain, hindbrain and midbrain, which other baseline methods and PRESENT failed to distinguish. For further validation, we increased the noise level of datasets by converting lower fragment counts to zero with growing dropout rates for each sample, according to a study that models the dropout events in single-cell data^73^. As shown in Fig. 5i, PRESENT-SN and PRESENT-SP gained greater advantages as the noise level increased, highlighting the improved benefits of integrating prior information from reference data with the growing dropout rates in spatial ATAC-seq datasets.

We also carried out extensive experiments on SRT axolotl brain datasets generated by Stereo-seq, which include six samples from different developmental stages, including stage 44, stage 54, stage 57, juvenile, adult and metamorphosis^6^. The performance of PRESENT was benchmarked with several recently developed SRT methods, involving SpaGCN^31^, STAGATE^32^, PAST^19^ and SpatialPCA^74^. As shown in Supplementary Figs. 15-21, experimental results demonstrated the consistent superiority of PRESENT in cross-modal representation, spatial visualization and domain identification of SRT data. In addition, utilizing an SRT regenerated axolotl brain data at 20 days post-injury as reference data^6^, PRESENT-SP again gained promoted performance through leveraging prior information contained in reference data (Supplementary Fig. 15). In summary, PRESENT not only promotes effective cross-modal representation in both spatial ATAC-seq and SRT data, but also empowers the integration of reference data to address the low sequencing depth and signal-to-noise ratio of spatial datasets.

### PRESENT enables vertical and horizontal integration of spatial single-omics samples

Integrated analysis of multiple samples across different developmental stages (vertically serial tissue samples) and dissected areas (horizontally adjacent tissue samples), commonly referred to as vertical and horizontal integration, provides a comprehensive approach for deciphering the hierarchical structures of cellular organizations^22^. We first evaluated the performance of PRESENT in the integrative analysis of vertically serial tissue samples, using the mouse embryo samples from three developmental stages (E15.5, E13.5 and E12.5) generated by spatial ATAC technique^9^. Due to the lack of methods specially designed for the multi-sample integration of spatial ATAC data, we compared PRESENT with six methods developed for single-cell data (MNN^65^, Scanorama^25^, Harmony^24^, SCALEX^66^, scBasset-BC^67^ and SnapATAC2^64^) and two recently proposed SRT methods (STAligner^27^ and GraphST^22^). We employed the benchmarking framework consisting of 14 metrics to simultaneously accessed the batch effect removal and biological variance conservation capabilities of these integration methods (Methods). As shown in Fig. 6a, PRESENT achieved the best overall score for multi-sample vertical integration, ranking first in biological variance conservation and second in batch effects removal among all the integration methods. The UMAP visualization of spot embeddings across different developmental stages also demonstrated that, compared with other methods, PRESENT gained better performance in aggregating spots from the same domain but in different batches (Supplementary Fig. 22). We further performed spatial clustering based on the joint embeddings produced by different methods. As shown in Fig. 6b, c, Scanorama and STAligner could hardly distinguish different spatial domains across different developmental stages. MNN, Harmony, SCALEX and SnapATAC2 provided improved results by roughly recognizing forebrain and liver in E12.5 and E13.5 stages, but they still failed in other domains. In comparison, PRESENT not only identified forebrain (cluster 1) and liver (cluster 5) in all the three stages, but also gained satisfactory results in domains including hindbrain (cluster 2) and midbrain (cluster 8).

**Fig. 6.**
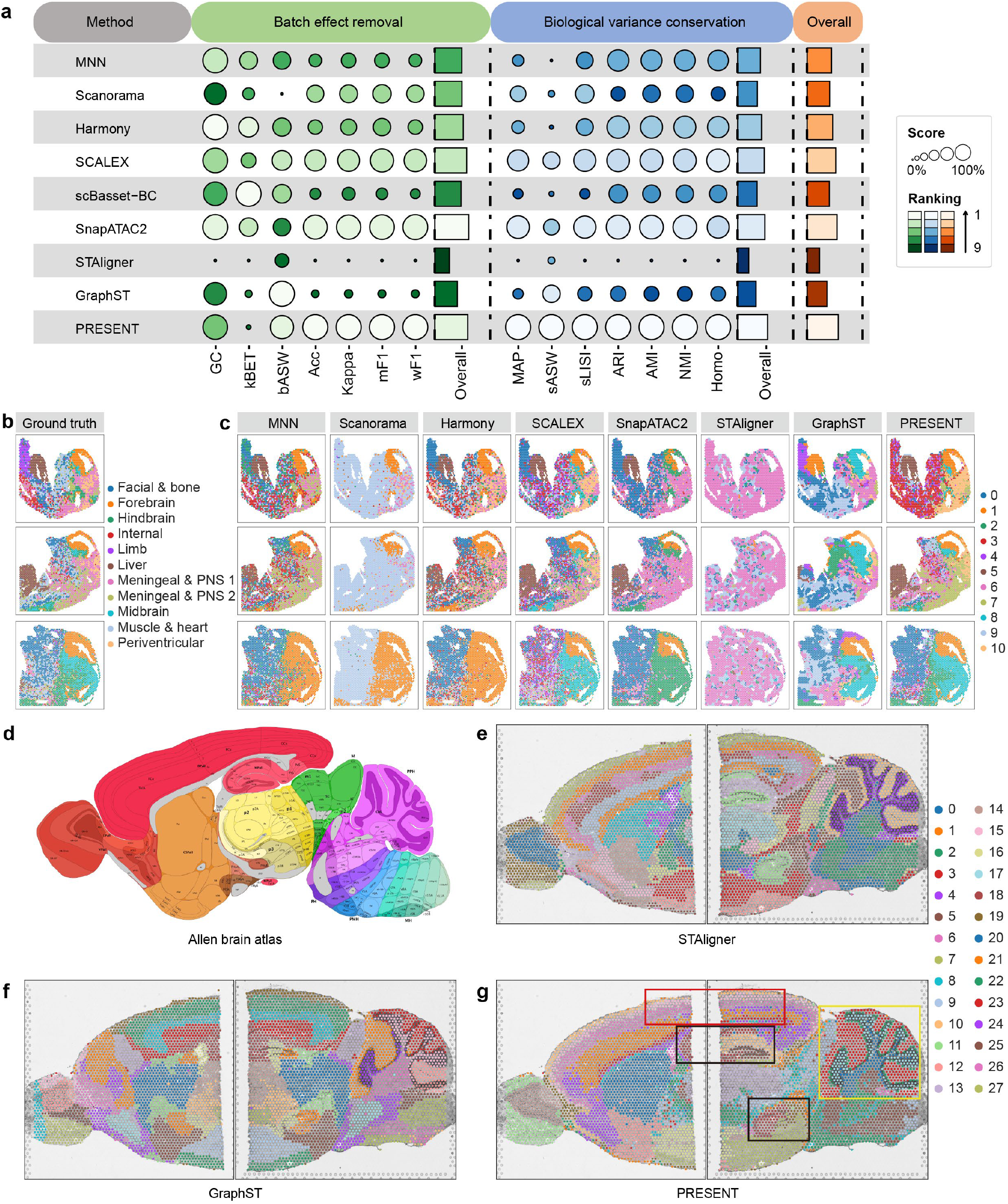
PRESENT enables vertical and horizontal integration of spatial single-omics samples. **a**, The quantitative evaluation of different integration methods on the three spatial ATAC mouse embryo samples using 14 metrics divided into two categories, namely batch effect removal and biological variance conservation. The category scores of these two aspects were calculated by averaging the metrics within each category. An overall score for each integration method was computed using a 40/60 weighted mean of the category scores for batch effect removal and biological variance conservation (Methods). **b**, The spatial visualization of spots across the three spatial ATAC mouse embryo samples colored by ground truth spatial domains and **c**, the spatial clusters identified based on different integration methods. The first, second and third row of **b** and **c** denotes the samples from E12.5, E13.5 and E15.5 stages, respectively. **d**, The anatomic annotation of the sagittal region in P56 mouse brain provided by Allen Reference Atlas. The joint spatial clustering results based on the latent embeddings obtained by **e**, STAligner, **f**, GraphST and **g**, PRESENT on the two horizontal mouse brain sagittal samples generated by the 10x Genomics Visium platform.

Furthermore, we collected two horizontal adjacent SRT mouse brain samples generated by the 10x Genomics Visium platform, which involves a sagittal anterior dissected section and a sagittal posterior dissected section. We applied two SRT integration methods, namely STAligner^27^ and GraphST^22^, and PRESENT to perform integrated analysis of these horizontally adjacent brain samples and utilized the anatomic annotation of the sagittal region in postnatal day 56 (P56) mouse brain provided by Allen Reference Atlas^75^ to intuitively compare the performance of these methods (Fig. 6d). As shown in Fig. 6e-g, compared with STAligner and GraphST, analytical results of PRESENT achieved improvements in shared and sample-specific domain identification. For example, PRESENT captured the classical hierarchical structures, i.e., from outer layer 1 to inner layer 6, of dorsal pallium in both sagittal anterior and posterior sections, as illustrated in the red rectangle of Fig. 6d. In addition, PRESENT successfully distinguished the typical cortex Ammonis (CA, cluster 18) and dentate gyrus (DG, cluster 25) structures simultaneously in the hippocampus (black frames in Fig. 6d), which have been proved to be associated with mood, anxiety and personality disorders caused by various factors^76^. We further conducted differentially expressed analysis for domain CA and domain DG, acquiring marker genes including *C1ql2* and *Hpca*, respectively (Supplementary Figs. 23 and 24). According to previous studies, *C1ql2* serves as a marker gene for dentate gyrus granule cells (DGCs) enriched in the DG structure^77^, while *Hpca* is the encoding gene for a high-affinity calcium-binding protein restricted to the central nervous system, abundant in pyramidal cells of the hippocampus CA region^78^.

Moreover, PRESENT also exhibited great potential in capturing sample-specific functional domains and uncovering biological implications in identified spatial domains. PRESENT successfully revealed the sophisticated substructures of the cerebellar vermis, which only exists in the sagittal posterior section of mouse brain and plays important roles in coordinating movements of the central body such as trunk, head, and proximal limbs^79^. Specifically, PRESENT successfully detected the molecular layer (cluster 3), Purkinje cell layer (cluster 22) and internal granular layer (cluster 17) of cerebellar vermis cortex and the white matter (cluster 20) of cerebellar vermis. The differentially expressed genes in the corresponding domains have also been validated in previous studies, such as *S100b*^80^, *Car8*^*81*^, *Atp1a1*^*82*^ and *Mbp*^83^ (Supplementary Figs. 25-28). For example, *Car8* and *Atp1a1*, the mutation of which cause the motor coordination defect and dominant Charcot-Marie-Tooth type 2, were proved to be highly expressed in the granular layers and Purkinje cells in recent studies^*81, 82*^, respectively. In summary, these experimental results demonstrate the superior performance of PRESENT in the integrative analysis of both vertically serial and horizontally adjacent spatial samples generated by various spatial omics technologies, thereby enabling the discovery of biological insights within identified spatial domains across multiple samples.

## Discussion

Recent breakthroughs in spatial omics sequencing technologies have enabled the simultaneous acquisition of spatial coordinate information and the regulatory status of the biomolecules within cells, such as chromatin regions, genes, and proteins, accumulating massive spatial omics data in repositories and databases^13, 14^. In this article, we proposed PRESENT, an effective and scalable contrastive learning framework, to address the cross-modality representation and multi-sample integration of spatial omics data, including spatial multi-omics, epigenomics and transcriptomics data. To capture the patterns of various omics layers, we designed omics-specific encoders and decoders for different omics data, where the transcriptomic and epigenomic decoders model the fragment counts via the zero-inflation negative binomial (ZINB) distribution^39^ and zero-inflation Poisson (ZIP) distribution^40^, respectively (Methods). In addition, a cross-omics alignment module built on a contrastive learning inter-omics alignment loss function^41^ and graph attention networks^37^ (GAT) were also introduced to capture the spatial dependency and complementary multi-omics information (Methods). We further extended the framework to the multi-sample integration of spatial omics data, addressing the challenges including the inconsistency of spatial coordinates between samples as well as the intricate coupling between batch effects and biological signals. Specifically, we adopted a contrastive learning-based inter-batch alignment strategy^42^ and a batch-adversarial learning strategy^43^ to eliminate batch effects as well as an intra-batch preserving loss to preserve biological signals simultaneously. To validate of the model design, we have conducted a comprehensive ablation study, as shown in Supplementary Figs. 29 and 30.

Through extensive experiments on different species involving human, mouse and axolotl, we highlighted the superior performance of PRESENT in cross-modality representation and multi-sample integration of spatial omics data generated by various spatial sequencing technologies, encompassing spatial multi-omics techniques (spatial RNA-Protein^11, 12^ and spatial RNA-ATAC^12^) and spatial single-omics technologies (spatial ATAC^9^ and spatial transcriptomics^6^). Specifically, PRESENT effectively integrated the spatial dependency and complex multi-omics information contained in spatial multi-omics or spatial single-omics techniques (Figs. 2, 3 and 5). Additionally, PRESENT also showed great potential for leveraging various reference data based on the BNN module to address issues related to low sequencing depth and signal-to-noise ratio in spatial omics data (Fig. 5). More importantly, PRESENT can be further extended to the integrative analysis of multiple spatially resolved samples across distinct dissection areas and developmental stages, thereby promoting the detection of hierarchical structures and biological functions within cellular organizations from spatiotemporal views (Figs. 4 and 6).

Nevertheless, several avenues remain for further improvement of PRESENT. First, Due to technical and cost limitations, the number of unpaired spatial samples exceeds that of paired spatial samples^84^, emphasizing the importance of unpaired integration. While PRESENT can anchor shared spots to analyze paired multi-omics sequencing data or leverage shared genomic features to integrate multiple spatial samples, it still struggles with the integration of unpaired samples with distinct feature space^84^, such as the integration of two samples generated by spatial ATAC-seq and SRT techniques, respectively. Second, explorative analyses based on single-cell data and spatial data are limited by lack of spatial information and low spatial resolution, respectively. As an integrative framework for spatially resolved omics data, PRESENT could be further extended to integrate single-cell and spatial data, computationally addressing these technological limitations. Third, beyond integrating samples into the latent space, the alignment of spatial coordinates in physical space also plays an important role in joint analysis of multiple spatial samples^23, 26^. Finally, with the development of spatially resolved sequencing technology, imaged-based spatial techniques, such as 10x Genomics Xenium, produce high-resolution histology image of dissected samples, providing additional morphological features for enhanced analysis of spatial samples.

## Methods

This study aims to investigate spatially functional domains as well as biological regulatory processes within microenvironments through incorporating cross-modality information, including spatial coordinate data and omics data, which involves epigenomic data generated by the assay for transposase-accessible chromatin with sequencing (ATAC), transcriptomic data measured by RNA sequencing (RNA), and proteomic data sequenced by antibody-derived tags (ADT). To achieve this goal, we have established a self-supervised contrastive learning framework, namely PRESENT, for the cross-modality representation and multi-sample integration of spatial multi-omics, epigenomics and transcriptomics data. Furthermore, we have also systematically evaluated the performance of PRESENT on massive real-world spatial datasets across various sequencing technologies, species and tissues.

### Cross-modality representation of spatial multi-omics data

First, we detail the modeling approach of PRESENT for cross-modality representation of spatial multi-omics data, using a complex scenario as an example, which involves the comprehensive integration of spatial coordinate data, epigenomic data (ATAC data), transcriptomic data (RNA data), and proteomic data (ADT data). As shown in Fig. 1a, PRESENT is based on a multi-view autoencoder framework and comprises omics-specific encoders, a cross-omics alignment module, and omics-specific decoders, aiming to incorporate the cross-modality information into a low-dimensional feature matrix.

### Multi-modality input of PRESENT

The input of PRESENT is a spatial coordinate matrix ***M*** ∈ *R*^*n*×2^ and three fragment count matrices, namely the ATAC data ***X***^***ATAC***^ ∈ *R*^*n*×*r*^, RNA data ***X***^***RNA***^ ∈ *R*^*n*×*g*^ and ADT data ***X***^***ADT***^ ∈ *R*^*n*×*p*^. Here *n* denotes the number of spots, *r* the number of genomic regions, *g* the number of highly variable genes and *p* the number of input proteins. Before feeding the spatial and omics data into the omics-specific encoders, we transform the spatial coordinate matrix ***M*** into an undirected neighborhood graph ***G*** through adding undirected edges between all spots and their *k* nearest neighbors, where *k* is set to 6 by default. Afterwards, we apply term frequency-inverse document frequency (TF-IDF) normalization to ATAC data^85-88^, library size normalization followed by log-transformation to RNA data^19, 31, 32^ and the centered log ratio (CLR) transformation to ADT data ^17, 89, 90^, obtaining normalized omics data 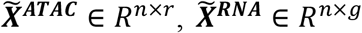 and 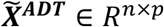, respectively.

### Omics-specific encoder

Graph neural networks, such as graph attention networks^37^ (GAT), have showcased successful application in integrating spatial and transcriptomic information for spatial transcriptomics^19, 22, 31, 32^. On the other hand, research has proved that introducing weight uncertainty into models through Bayesian neural networks^38^ (BNN) improves generalization in nonlinear regression problems^38^ and offers an extension for leveraging prior information from existing sequencing data^19^. Inspired by these studies, we design an omics-specific encoder, consisting of a GAT module and a BNN module, to capture spatially organized and biologically regulatory signals embedded in each omics layer. In the following two subsections, taking the ATAC omics-specific encoder as an example, we delineate the detailed design of GAT and BNN module.

#### Graph attention network

Given the input undirected neighborhood graph ***G*** and the normalized ATAC matrix 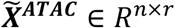, we denote the normalized accessibility vector of spot *i* in 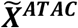 as 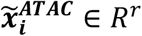 and the spatial neighborhood of spot *i* as 𝒩_i_. Initially, we acquire the embedding from the first hidden layer of spot *i*, i.e., 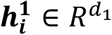 using a fully connected network (FCN):

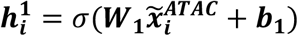

where *d*_1_ represents dimension of the first hidden layer, *σ* indicates the combination of layer normalization and exponential linear unit (ELU) activation function, and 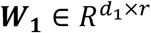 and 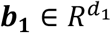 indicate the weight and bias parameters of the FCN, respectively. Afterwards, The second hidden-layer embedding of spot *i*, namely 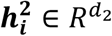, is obtained based on a graph attention layer^37^, depicted as:

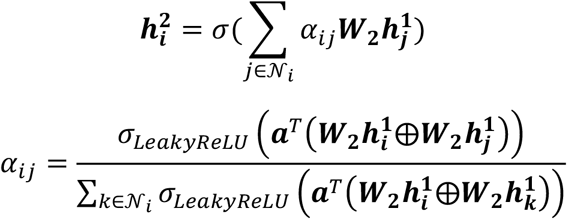

where *α*_*ij*_ is the attention score between spot *i* and *j, d*_2_ denotes the dimension of the second hidden layer, 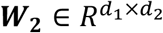 and 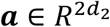 are learnable parameters of the graph attention layer, *σ*_*LeakyReLU*_ denotes the LeakyReLU activation function with a negative slope of 0.2, and ⨁ refers to the concatenation function. Finally, 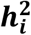 undergoes transformation by another graph attention layer to obtain 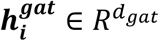 as the output, where *d*_*gat*_ denotes the output dimension of GAT module.

#### Bayesian neural network

According to a previous work, BNN module provides an opportunity to leverage biological prior information from existing sequencing data to improve the analytical performance in spatial transcriptomic data^19^. Inspired by this, PRESENT includes the BNN module, which assumes that the parameters of weight matrix, denoted as 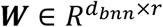, follow Gaussian distributions. Specifically, each element of ***W*** is governed by a Gaussian distribution parameterized by the corresponding elements in the mean matrix ***µ***^***b***^ and variance matrix 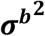, depicted as:

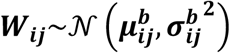

In addition, ***W***_***ij***_ is also governed by a Gaussian prior parameterized by mean 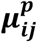 and variance 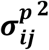. Inheriting the original settings^19^, we calculate the PCA projection weight matrix of normalized ATAC matrix 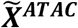 as the mean matrix ***µ***^***p***^ and utilize the standard deviation of all the elements in mean matrix ***µ***^***p***^ to construct the standard deviation matrix ***σ***^***p***^. The distance between the prior and the distributions of the parameters in BNN module is restricted with Kullback-Leibler divergence (KLD):

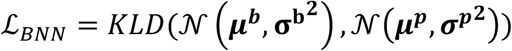

It is notable that the parameters of prior Gaussian distributions, i.e., the mean matrix ***µ***^***p***^ and the variance matrix 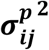, can also be constructed with reference ATAC data, enabling PRESENT to incorporate prior information from existing ATAC-seq data generated by various technologies, including single-cell ATAC-seq or spatial ATAC-seq. The unification of peak sets between reference ATAC-seq data and target ATAC-seq data is conducted according to the merging objects vignette provided by the Signac package^91^.

The output embeddings of spot *i* from BNN module, i.e., 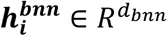, is denoted as:

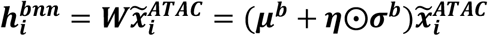

where *d*_*bnn*_ denotes the output dimension of BNN module, ***η*** is the standard Gaussian noise matrix and ⨀ denotes element-wise product.

Finally, the output embedding of spot *i* from the ATAC omics-specific encoder, denoted as 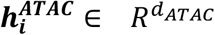, is the concatenation of output embeddings from the GAT and BNN module:

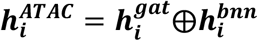

where *d*_*ATAC*_ = *d*_*gat*_ + *d*_*bnn*_ is the output dimension of omics-specific encoder for ATAC data.

### Cross-omics alignment module

Based on the omics-specific encoders, the embeddings of different omics layers are separately extracted. However, accurately deciphering sophisticated gene regulation process requires integrating the status of biological macromolecules in cells from different omics perspectives^15-18^. Inspired by a recent study that achieved multi-modal entity alignment through contrastive learning^41^, we designed a cross-omics alignment module to model the complex regulatory interactions between different omics layers.

Given the omics-specific embeddings of spot *i* obtained by omics-specific encoders, including 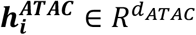 for ATAC, 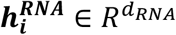 for RNA and 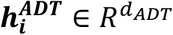 for ADT, we utilized a multi-layer perception (MLP) module to fuse the embeddings from different omics layers to acquire a cross-omics representation of spot *i*, denoted as:

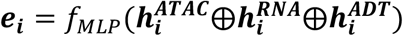

where ***e***_***i***_ ∈ *R*^*d*^ represents the final cross-omics representation of spot *i, f*_*MLP*_ denotes the transformation function of the MLP module, *d* is the latent dimension of PRESENT, and *d*_*ATAC*_, *d*_*RNA*_ and *d*_*ADT*_ refers to the output dimensions of omics-specific encoders for ATAC, RNA and ADT data, respectively. The MLP module is composed of three FCNs, the first two of which are followed by a layer normalization and an ELU activation function.

However, only the MLP module cannot perfectly incorporate omics-specific knowledge from different omics layers into the final cross-omics embeddings^41^, which is also illustrated in our ablation study shown in Supplementary Fig. 29. Therefore, we design an inter-omics alignment (IOA) loss to minimize the multi-omics directional KLD over the output distributions between the joint embeddings and other omics-specific embeddings:

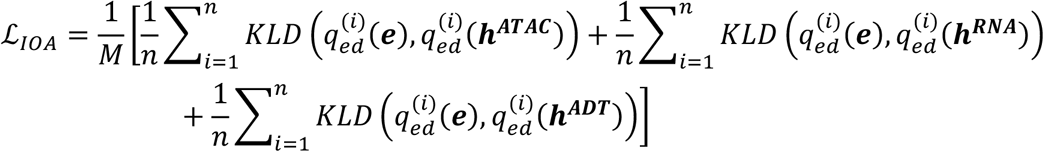

where *M* represents the number of omics layers, ***e, h***^***ATAC***^, ***h***^***RNA***^ and ***h***^***ADT***^ denotes the joint, ATAC, RNA and ADT embeddings respectively, and 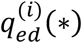 represents the embedding distribution of spot *i* over other spots, defined as (taking ***e*** as an example):

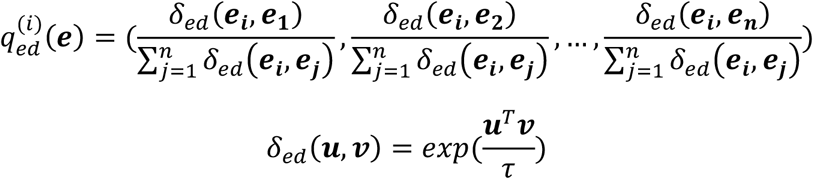

where *τ* is a temperature parameter and set to 0.1 according to the original study^41^.

### Omics-specific decoder

The patterns and distributions of sequencing data vary significantly across different omics layers, revealing complementary biologically regulatory information^15-18^. For example, ADT sequencing data, which focuses on dozens of surface proteins, provides intuitive and reliable quantification of cellular activities with low dimensionality, few dropout events and low sparsity^17^. In contrast, RNA and ATAC data, which measure the expression levels of genes and accessibility scores of genomic regions respectively, suffer from high dropout rates, dimensionality, and sparsity^92^. Therefore, we designed omics-specific decoder for each modality to address the discrepancies between omics layers.

Inspired by a recently published study which modeled the fragment counts of single-cell ATAC-seq data with Poisson distribution^40^, we fit the fragment counts of spatial ATAC-seq data ***X***^***ATAC***^ ∈ *R*^*n*×*r*^ with the zero-inflation Poisson (ZIP) distribution to accommodate the lower capture rate of spatial technologies^9, 12^. Taking the fragment count vector of spot *i*, i.e., 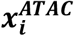, as an example, the omics-specific decoder of ATAC data as well as the probability model is described as:

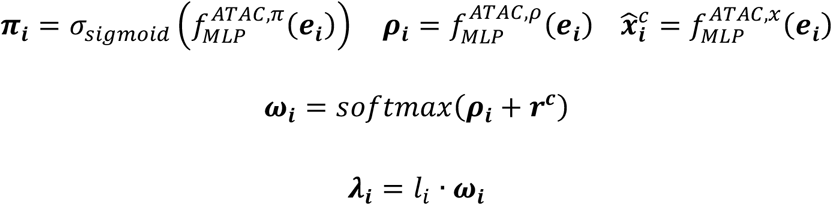

where *σ*_*sigmoid*_ represents the sigmoid activation function, ***r***^***c***^ is designed to capture region specific bias such as peak length, 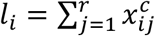 refers to the total fragment counts of spot *i*, 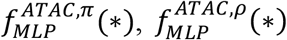 and 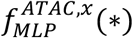 are three output layers of 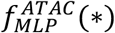, the MLP module of ATAC omics-specific decoder. Specifically, 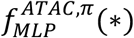 and 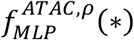 output parameters of the ZIP probability model, while 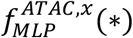 outputs reconstructed normalized profiles of ATAC data. Then for each region *j* = 1,2, …, *r*, the fragment count of *i*-th spot 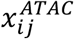 satisfies:

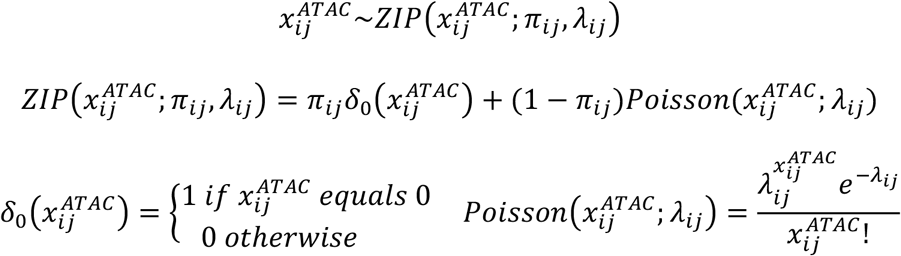

Therefore, the negative log likelihood (NLL) loss function of the ZIP probability model is:

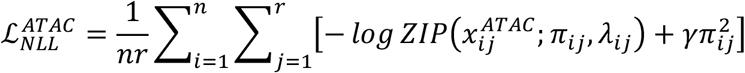

where the 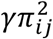 refers to a regularization term and *γ* is set to 0.5 by default. The reconstruction loss of ATAC omics layer is measured with mean squared error (MSE):

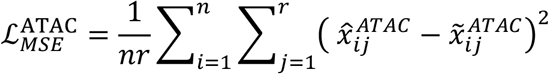

Similarly, we also model the counts of RNA data ***X***^***RNA***^ ∈ *R*^*n*×*g*^ using the zero-inflation negative binomial (ZINB) distribution according to a single-cell RNA-seq analytical research^39^, denoted as:

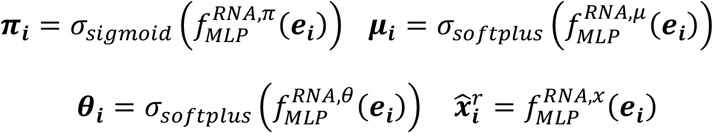

where *σ*_*softplus*_ refers to the softplus activation function, 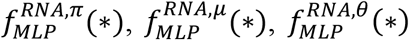 and 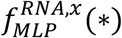 are four output layers of 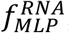, the MLP module of the RNA omics-specific decoder. Then the count of spot *i* on gene *j* satisfies the following model:

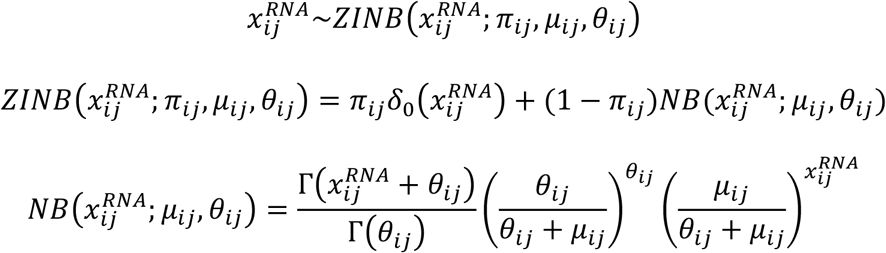

The NLL loss function of the ZINB probability model for RNA data can be denoted as:

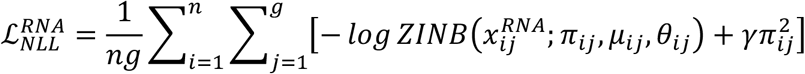

where the regularization parameter *γ* is also set to 0.5 by default. In addition, the reconstruction loss of RNA omics layer is depicted as:

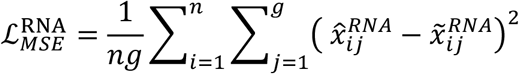

Compared with ATAC and RNA data, lower dropout rates, dimensionality and sparsity make the analysis of ADT data less challenging. Therefore, the ADT omics-specific decoder is built on the MLP module with one output layer to reconstruct the normalized expression level of proteins, denoted as:

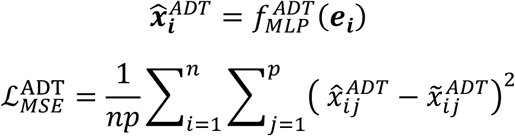

Therefore, the overall NLL loss and reconstruction loss of PRESENT are summarized as:

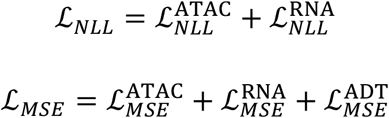

### Loss function for cross-modality representation of spatial multi-omics data

Based on the above descriptions, the overall loss function for cross-modality representation of spatial multi-omics data consists of four parts, including the loss of BNN module, the inter-omics alignment loss, the negative log likelihood loss of ZIP and ZINB probability model, and the mean squared error loss for reconstruction of normalized omics profiles, denoted as:

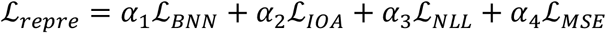

where *α*_1_, *α*_2_, *α*_3_ and *α*_4_ are coefficients of the corresponding loss term. The default values of *α*_1_, *α*_2_, *α*_3_ and *α*_4_ are all set to 1.

### Multi-sample integration of spatial multi-omics data

Multi-sample integration offers a comprehensive perspective for deciphering the structures of cellular organizations^22, 23, 26, 27^. For example, integrated analysis of samples under different stages can reveal developmental dynamics^27, 28^, while integrating disease and normal samples facilitates the identification of disease-specific domains and promotes the investigation of disease mechanisms^27, 29^. Therefore, we have extended the PRESENT framework to multi-sample integration of spatial multi-omics data and further incorporated mechanisms to address the discrepancies of spatial coordinates as well as batch effects between samples. Overall, the multi-sample integration workflow based on the PRESENT framework involves two stages. The first stage serves as a warmup to pretrain the PRESENT model, while the second stage leverages a cyclic iterative optimization progress to eliminate batch effects while preserving biological variations across samples. Analogous to cross-modality representation of spatial multi-omics data, we use a scenario of integrating two spatial multi-omics sequencing samples as an example to illustrate the workflow of multi-sample integration with PRESENT, where ATAC, RNA, ADT, and spatial coordinate data are simultaneously provided for each sample (Fig. 1b).

### Feature sets unification

The first step in multi-sample integration is to align the feature sets of different sequencing samples, ensuring that all samples share a common feature space. For ATAC data, following the merging objects vignette provided by the Signac package^91^, a unified set of peak regions is first obtained by merging all intersecting peaks from all the samples. The fragment counts of each spot within the unified peak set are then quantified. For RNA and ADT data, whose feature sets are predefined genes or proteins, we directly leverage the intersection of gene or protein sets of all the samples as the unified gene or protein set. After the gene intersection step, highly variable genes will be selected based on the unified gene set. For convenience, we denoted the ATAC, RNA, ADT and spatial data of the two samples as 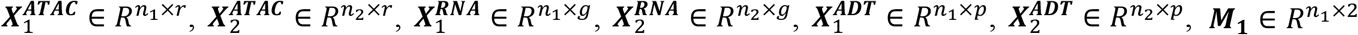, and 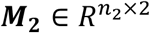, respectively. Here, *r* represents the number of unified peak regions, *g* the number of highly variable genes, *p* the number of unified proteins, and *n*_1_ and *n*_2_ the number of spots of the two samples. The unified ATAC, RNA, ADT data are then concatenated and preprocessed using the methods described in the cross-modality representation section, obtaining normalized data, i.e., 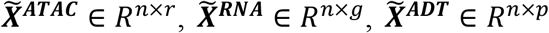 and ***M*** ∈ *R*^*n*×2^, where *n* = *n*_1_ + *n*_2_ represents the number of spots from the two samples.

### Cross-sample graph construction

Due to inconsistent dissection and placement of sequencing samples, the spatial coordinates between these samples cannot be directly compared with each other, resulting in the failure of cross-modality representation methods in capturing relationships across samples^23, 26^. To address this gap, we proposed a strategy to construct a cross-sample graph. Firstly, we find intra-sample spatial neighbors for each spot based on the spatial coordinate matrix, building undirected edges between all spots and their *k* nearest neighbors in Euclidean space from the same sample. Secondly, we find inter-sample neighbors based on omics features, including ATAC, RNA and ADT profiles, to establish inter-sample undirected edges between all spots and their *k* most similar neighbors in cosine space from other samples. Note that the parameter *k* is set to 6 by default. Finally, we acquire a cross-sample graph which consists of both intra-sample edges and inter-sample edges.

### First-stage pretraining

In the first stage, we pretrain the PRESENT framework based on the cross-sample graph and the concatenated omics data of different samples, including 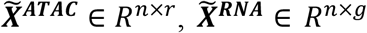 and 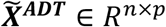. To integrate batch information into the PRESENT model, we concatenate the one-hot encoded batch labels with the output of the cross-omics alignment module, using the combined matrix as the input of the omics-specific decoders. The objective function and training process of PRESENT model during the first pretraining stage are identical to those used for cross-modality representation of individual samples, as detailed in the previous section.

### Second-stage integration

The technical noise between different samples, i.e., batch effects, have not been fully addressed in the first stage. Therefore, we incorporate a contrastive learning strategy and a batch-adversarial learning strategy in the second stage to eliminate the batch effects across samples while preserving the biological signals simultaneously, as detailed in the following subsections.

#### Inter-batch alignment

We designed an inter-batch alignment (IBA) loss based on contrastive learning^42^, aiming to align spots with similar omics features across different samples. Specifically, we first find mutual nearest neighbors from other samples as the positive set of for spot *i*, denoted as 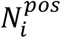, referring to the inter-sample neighbors that are connected to spot *i* in the cross-sample graph. Second, we randomly select 1/4 proportion of spots from the same sample as the negative set of spot *i*, depicted as 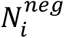. Finally, the IBA loss of spot *i* is:

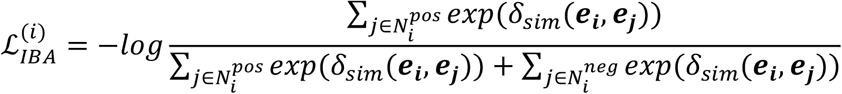

where ***e***_***i***_ ∈ ***R***^*d*^ denotes the latent representation of spot *i, d* is the output dimension of PRESENT and *δ*_*sim*_ denotes the cosine similarity. The inter-batch alignment loss of all the spots is:

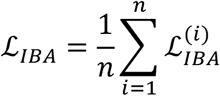

#### Batch-adversarial learning

Inspired by a domain-adversarial learning research^93^ and its’s success in multi-batch integration of single-cell ATAC-seq data^43^, we implement the batch-adversarial learning strategy to remove batch effect across spatial samples. Specifically, we introduce a batch-discriminator, aiming to predict the batch index of each spot based on its latent embedding, while the omics-specific encoders and the cross-omics alignment module are treated as the generator, aiming to produce latent embeddings without batch effect, thereby confusing the batch-discriminator in an adversarial manner. The loss function of the batch-adversarial learning is:

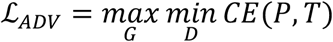

where *G* refers to the generator, *K* denotes the batch-discriminator, *CE* represents the cross-entropy loss, *Z* denotes the predicted batch indices and *T* refers to the ground truth batch labels.

#### Intra-batch preserving

The contrastive learning-based inter-batch alignment strategy and the batch-adversarial learning strategy aim for batch correction across samples. However, a key challenge of multi-sample integration lies in eliminating batch effects while simultaneously preserving the biological signals within each individual sample. We therefore proposed an intra-batch preserving (IBP) loss to maintain the biological pattern of latent representations within each sample during training process, denoted as:

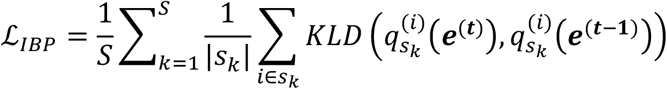

where *S* represents the number of samples, *s*_*k*_ denotes the spots in sample *k*, | *s*_*k*_ | denotes the number of spots in sample *k*, ***e***^(***t***)^ and ***e***^(***t*** − **1**)^ refer to the joint embeddings of all spots in current epoch and the last epoch, respectively. 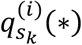 represents the embedding distribution of spot *i* over other spots within sample *k*, defined as (taking the embeddings of current iteration ***e*** ^(***t***)^ as an example):

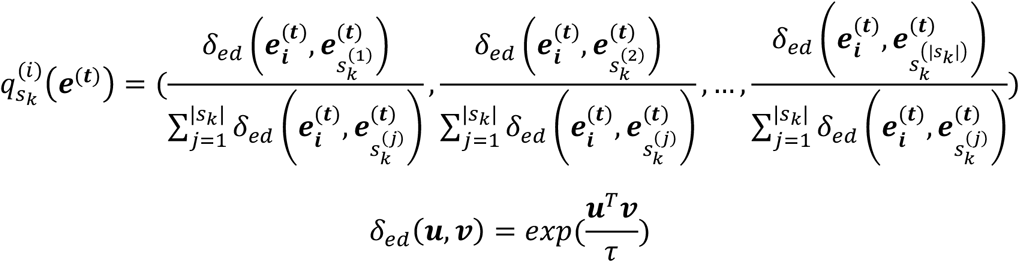

where 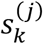 denotes the *j*-th spot in sample *k, τ* is a temperature parameter and set to 0.1 by default^41^. As shown in the above formula, the IBP loss is designed to preserve the overall manifold structure of spots within each sample during training process.

#### Training process of the second stage

The training process of the second stage is carried out in a cyclic iterative manner. First, we obtain the joint embeddings of multiple samples using the trained PRESENT model. Next, we construct a cross-sample graph according to the strategy described above, where intra-sample connections are established based on the spatial coordinates of spots and the inter-sample edges are based on the cosine similarity between the newly acquired joint embeddings of spots from different samples. Finally, we train the PRESENT model for one epoch using the newly constructed cross-sample graph. This cyclic iterative optimization process, aiming to iteratively refine the cross-sample graph, is repeated until the model converges. When updating the PRESENT model, we first update the parameters of the batch-discriminator using the cross-entropy loss 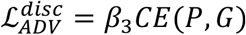, with parameters of the generator (i.e., the omics-specific encoders and the cross-omics alignment module) and omics-specific decoders fixed. Next, we fix the parameters of batch-discriminator and update the parameters of generator and omics-specific decoders based on an overall loss function, including the cross-modality representation loss, the inter-batch alignment loss, the intra-batch preserving loss and the negative cross-entropy loss for the batch-adversarial learning strategy:

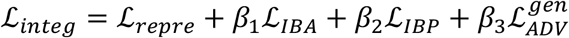

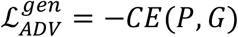

where *β*_1_, *β*_2_ and *β*_3_ are coefficients of the corresponding loss function and set to 1 by default.

### Applications of PRESENT to spatial single-omics data

PRESENT is highly versatile and can be seamlessly adapted from spatial multi-omics data to spatial single-omics data, such as SRT and spatial epigenomics data, by simplifying the multi-view autoencoder framework into a single-view autoencoder. Specifically, for cross-modality representation or multi-sample integration of spatial single-omics data, only the respective omics-specific encoder and decoder are required. For example, only the ATAC omics-specific encoder and decoder are required when modeling spatial single-omics ATAC-seq data, while only the RNA omics-specific encoder and decoder are needed when modeling spatial transcriptomic data. Additionally, both the cross-omics alignment module and the associated inter-omics alignment loss are excluded from the PRESENT model in these cases, as spatial single-omics data consist of only a single omics layer.

### Implementation details

With the development of spatial omics sequencing technologies, the number of spots in sequenced tissue has been quickly increasing from thousands to tens and even hundreds of thousands^9, 10, 13, 14, 94^. PRESENT is built upon graph neural networks, which face challenges such as memory explosion and neighborhood expansion when the number of nodes in graph grows large^95, 96^. To address these problems, we adopted necessary sparse adjacency matrix implementation based on the PyG framework when deploying our model. Additionally, we implemented the ClusterGCN algorithm to divide the entire graph into multiple small subgraphs^95^, which enables our model to be trained on GPUs with limited memory and extend to larger spatial omics data.

The input dimensions of ATAC and RNA data are generally much higher than the ADT data, resulting in different hidden dimension settings for them. Specifically, the hidden and output dimensions of omics-specific encoders for ATAC and RNA data were (1024, 512) and 50, respectively, while those of omics-specific encoder for ADT data were 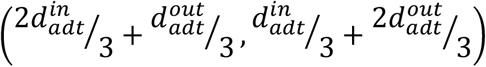 and 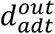. Here 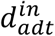 denotes the input dimension of ADT data and 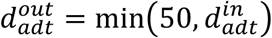 refers to the output dimension of omics-specific encoder for ADT data. The final output dimension of PRESENT was set to 50 by default. Additionally, the hidden dimensions of omics-specific decoders were symmetric to the corresponding omics-specific encoders, i.e., 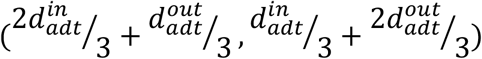 for ADT data and (512, 1024) for ATAC and RNA data. We trained the model for at most 100 epochs using the Adam optimizer, updating the parameters with an initial learning rate of 1e-3 and a weight decay of 1e-4.

### Data collection and description

#### 10x Visium RNA-Protein human lymph node data

The human lymph node dataset, generated by 10x Genomics Visium transcriptomics and proteomics co-profiling technology, was collected from a spatial multi-omics study^35^. This dataset was manually annotated by the clinician group according to the histological image^35^, containing the spatial coordinates and the expression level of 18,085 genes and 31 proteins on 3,484 spots.

#### SPOTS mouse spleen data

The mouse spleen data^11^ was generated by a high-throughput spatial transcriptomics and proteomics co-profiling technology, namely Spatial PrOtein and Transcriptome Sequencing (SPOTS). This dataset provides the spatial coordinates and the expression level of 32,285 genes and 21 proteins on 2,568 spots.

#### MISAR-seq mouse brain data

The MISAR-seq mouse brain data^12^ provides the paired sequencing data of spatial gene expression and chromatin accessibility during mouse brain development. We collected four manually annotated samples from original study^12^, including E11.0 (1,258 spots), E13.5 (1,777 spots), E15.5 (1,949 spots) and E18.5 (2,129 spots). The manual annotations were produced with reference to Kaufman’s Atlas of Mouse Development and Allen Brain Atlas^12^. We called peak for each sample based on the MACS2 v2.2.9.1 package, obtaining 242,024, 271,126, 265,014 and 294,734 peak regions for downstream analyses, respectively.

#### Single-nucleus ATAC-seq and spatial ATAC mouse embryo data

The single-nucleus ATAC-seq mouse embryo data^9^ measured the chromatin accessibility of 1,879 cells from three developmental stages, including E12, E13 and E15. The spatial ATAC mouse embryo data^9^ is composed of six samples, including E12.5-S1 (3,274 spots), E12.5-S2 (3,903 spots), E13.5-S1 (3,160 spots), E13.5-S2 (2,970 spots), E15.5-S1 (2,246 spots) and E15.5-S2 (2,234 spots). Associated domain annotations were also provided within these samples, which have been validated through extensive experiments in the original study^9^. The single-cell ATAC-seq datasets and spatial ATAC datasets were merged and 248,047 peak regions were called based on MACS2 v2.2.6 package.

#### Stereo-seq axolotl brain data

We collected the Stereo-seq axolotl brain developmental datasets from a recent study which explored the regeneration and development of axolotl telencephalon using spatial transcriptomic atlas^6^. The datasets consist of six sections from developmental stage 44 (1,477 spots), stage 54 (2,929 spots), stage 57 (4,410 spots), juvenile (11,698 spots), adult (8,243 spots), and metamorphosis (7,441 spots), measuring the expression level of 12,704 genes. These samples were manually annotated with reference to eHistology Kaufman Annotations and Allen Brain Atlas^12^.

#### 10x Visium mouse brain data

The spatial transcriptomic postnatal day 56 (P56) mouse brain data, including a sagittal anterior section and a sagittal posterior section, were collected from the 10x Genomics Visium platform. The mouse sagittal anterior section contains the expression level of 32,285 genes on 2,823 spots, while the posterior section provides the expression level of 32,285 genes on 3,289 spots.

### Experiment setup and baseline methods

#### Cross-modality representation of 10x Visium RNA-Protein human lymph node data

We excluded genes or proteins with zero expression across all spots. Baseline methods includes totalVI^46^, CiteFuse^47^, Seurat-v4^48^, SpatialGlue^35^ and spaMultiVAE^36^, all implemented following their respective tutorials.

#### Cross-modality representation of MISAR-seq mouse brain data

We filtered out peaks accessible in less than 3% spots and genes with zero expression in all spots for quality control. The filtered count matrices of ATAC data and RNA data along with the spatial coordinate matrix were then fed into the workflow of PRESENT and other baseline methods, involving SpatialGlue^35^, MOFA+^61^, scAI^62^, scMDC^63^, scMVP^21^, Seurat-v4^48^ and SnapATAC2^64^. Note that totalVI^46^, CiteFuse^47^ and spaMultiVAE^36^ were not used as they are specially designed for single-cell or spatial RNA-ADT data. All the baseline methods were implemented following the respective tutorials and default settings. Additionally, following the workflow of processing single-cell^97^ and spatial^19, 32, 74^ RNA-seq data, PRESENT selected 3,000 highly variable genes as the transcriptomic input for further analysis.

#### Multi-sample integration of MISAR-seq mouse brain data

We merged the peak sets of different mouse brain samples according to the merging objects vignette provided by the Signac package^91^, which first obtained a unified peak set by merging all intersecting peaks from all the samples and then quantified the fragment count of each spot on the newly obtained peak set. Consistent with the experimental settings in cross-modality representation, peaks accessible in less than 3% spots in the concatenated matrix were filtered out and 3,000 highly variable genes of transcriptomic data were selected for PRESENT. Due to the lack of methods specially designed for the integration of spatial multi-omics data, we compared the batch effect removal and biological variance conservation performance of PRESENT with six methods developed for single-cell data (MNN^65^, Scanorama^25^, Harmony^24^, SCALEX^66^, scBasset-BC^67^ and SnapATAC2^64^) and three recently proposed SRT multi-sample integration methods (STAligner^27^, Stitch3D^33^ and GraphST^22^). Specifically, MNN, Scanorama, Harmony, SCALEX could be applied to both transcriptomic and epigenomic data, denoted as RNA-* or ATAC-*, respectively, where * denotes respective methods. MNN, Scanorama and Harmony performed batch correction based on embeddings extracted from RNA or ATAC data, which were accomplished based on the workflow of SCANPY^97^ and EpiScanpy^98^, respectively. All the methods were implemented using default settings.

#### Cross-modality representation of spatial ATAC mouse embryo data

The spatial ATAC mouse embryo data firstly went through a quality control process, where peaks accessible in less than 0.5% spots were filtered out. Using the filtered ATAC count matrices as input, we then benchmarked the performance of PRESENT with four methods designed for single-cell ATAC-seq data, including SCALE^71^, cisTopic^72^ and SnapATAC2^64^, and three methods designed for SRT data, i.e., SpaGCN^31^, STAGATE^32^ and PAST^19^. All the baseline methods were implemented using their default settings, except for STAGATE and PAST. In original tutorials, the two SRT methods adopted a feature selection process which is specially designed for transcriptomic data and only selected 3,000 features, resulting significantly lower performance (Supplementary Fig. 31). Since other methods utilized all the input peaks for dimension reduction, we ignored the feature selection step of these two methods for fairness and better performance.

#### Cross-modality representation of Stereo-seq axolotl brain data

The genes with zero expression in all spots were filtered out. The filtered count matrices and the spatial locations were fed into PRESENT and other baseline methods, including SpaGCN^31^, STAGATE^32^, PAST^19^ and SpatialPCA^74^. All baseline methods were implemented with default settings in their tutorials. PRESENT again employed a gene selection procedure, obtaining 3,000 highly variable genes as input.

#### Multi-sample integration of spatial ATAC mouse embryo data

The three spatial ATAC mouse embryo samples from three developmental stages have the unified peak sets and were directly concatenated. Following the experimental settings in cross-modality representation of a single spatial sample, peaks accessible in less than 0.5% spots were filtered out. We benchmarked PRESENT with six integration methods developed for single-cell data (MNN^65^, Scanorama^25^, Harmony^24^, SCALEX^66^, scBasset-BC^67^ and SnapATAC2^64^) and two recently proposed SRT methods (STAligner^27^ and GraphST^22^). Here Stitch3D^33^ was not used as it encountered error. MNN, Scanorama and Harmony performed batch correction based on the latent embeddings obtained by EpiScanpy^98^. All the methods were implemented according to their default settings except for STAligner. Since other methods utilized all the input peaks, we ignored the feature selection step of STAligner to ensure better performance and fairness.

#### Multi-sample integration of 10x Visium mouse brain data

The intersected gene set of the two mouse brain samples, including a sagittal anterior section and a sagittal posterior section, were utilized as the unified gene set. Next, gene count matrices of the two samples were concatenated and genes with zero expression in all spots were filtered out. Similarly, 3,000 highly variable genes were selected as the input for PRESENT. PRESENT was compared with two recently developed SRT methods, i.e., STAligner^27^ and GraphST^22^, which were conducted according to their default settings. Specifically, we utilized PASTE^23^ to accomplish the initial coordinate alignment step for GraphST according to the original study^22^.

### Evaluation metrics

#### Spatial clustering metrics

We implemented four commonly used metrics^50, 51^ to quantitatively evaluate the performance of spatial clustering, including adjusted rand index (ARI), adjusted mutual information (AMI), normalized mutual information (NMI) and homogeneity (Homo).

ARI, ranging from 0 to 1, denotes the consistent adjustment of the expectations of rand index:

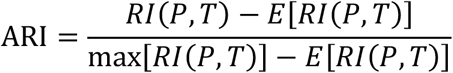

where *RI*(·) denotes the function of rand index, *T* refers to the ground truth labels, and *P* represents the clustering labels. Rand index is the probability of the agreement between the clustering result and the ground truth labels on a randomly selected spot set.

Mutual Information (MI) accesses the similarity between the clustering labels and ground truth labels, from which both AMI and NMI originate. AMI ranges from -1 to 1, while NMI ranges from 0 to 1. AMI is calculated as:

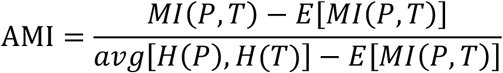

where *MI*(·) refers to the mutual information, *E*(·) represents the expectation function and *H* (·) denotes the function of information entropy. NMI can be calculated as:

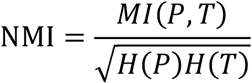

Homogeneity is another metric to measure the clustering performance. Specifically, if all the clusters involve only data points that are members of a single class in the ground truth labels, the clustering results could be interpreted as satisfying homogeneity, calculated as:

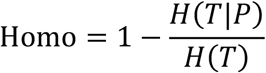

#### Integration metrics

According to a recent benchmarking framework for single-cell omics data integration, two important perspectives need to be considered when evaluating the performance of multi-sample integration, namely the batch effect removal and the biological variance conservation^68^.

To quantitatively evaluate the performance of batch effect removal, we introduced several metrics used in the benchmarking framework^68^, including graph connectivity(GC), k-nearest-neighbor batch-effect test (kBET) and batch average silhouette width (bASW). In addition, we also implemented a cross-validation strategy to evaluate the cross-batch annotation performance based on latent representations obtained by different integration methods, which is also an important assessment of batch correction^99^. Specifically, using the spots from a specific sample as the test set and those from other samples along with the domain labels as the training set, we transferred the domain labels of the training set into the test set based on a *k*-NN classifier and the joint embeddings of all the samples. *k*-NN classifier was used as its performance mainly depended on the quality of joint embeddings obtained by different methods. The performance of label transfer was evaluated with four commonly used metrics implemented in the scikit-learn package, i.e., accuracy (Acc), kappa, macro F1 score (mF1) and the weighted F1 score (wF1). After all the samples were used as the test set in a cross-validation manner, we calculated the average score for each metric to evaluate the performance of cross-batch annotation.

The quantitative evaluation for biological variance conservation involves the assessment of representative quality and the clustering performance. The representative quality was accessed via the mean average precision (MAP)^100^ and two metrics included in the benchmarking framework^68^, namely the spot average silhouette width (sASW) and spot local inverse Simpson’s index (sLISI), while the evaluation of clustering performance was based on the four metrics, i.e., ATI, AMI, NMI and Homo.

We calculated category scores of the batch effect removal and the biological variance conservation through averaging the metrics mentioned in the two aspects, respectively. We also calculated an overall score using a 40/60 weighted mean of the two category scores following previous studies^68, 100^.

### Downstream analyses

#### Visualization

Based on the latent representations obtained by various methods, we utilized the uniform manifold approximation and projection (UMAP)^52^ algorithm to further reduce the dimension of latent representation matrices into two-dimensional matrices for visualization. In addition, the two-dimensional UMAP and spatial coordinate matrices were visualized using the functions *scanpy.pl.umap()* and *scanpy.pl.spatial()* wrapped in the SCANPY package^97^.

#### Spatial clustering

We implemented a binary search strategy to tune the resolution of Leiden algorithm^49^, a graph-based clustering method, to match the number of spatial clusters as closely as possible with the number of ground truth domains. Specifically, we set a minimum resolution *res*_*max*_ and a maximum resolution *res*_*max*_, and obtain clustering results using a resolution equal to 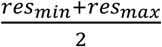. If the number of clusters exceeds the number of ground truth domains, *res*_*max*_ is updated to 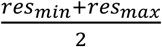. Otherwise, *res*_*max*_ is updated to 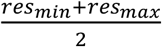. The iterative process continues until the number of clusters matches the number of ground truth domains or the current iteration reaches the maximum number of iterations.

#### Differentially expressed gene and protein analysis

We implemented differential analysis based on the Wilcoxon rank-sum test^54^ wrapped in *scanpy.tl.rank_genes_groups()* function of the SCANPY package^97^.

#### Gene ontology analysis

Given a specific gene set, we conducted gene ontology (GO) analysis using the clusterProfiler package^101^, which automated the process of biological-term classification and the enrichment analysis of gene clusters.

## Data availability

The single-nucleus ATAC-seq and spatial ATAC mouse embryo datasets^9^ are available at Gene Expression Omnibus (GEO) using the accession code GSE214991. The Stereo-seq axolotl brain datasets^6^ as well as the reference dataset, i.e., axolotl brain data at 20 days post-injury, can be obtained through their database https://db.cngb.org/stomics/artista/. The 10x Visium mouse brain datasets, including a sagittal anterior section and a sagittal posterior section, are accessible at the 10x Genomics websites https://www.10xgenomics.com/datasets/mouse-brain-serial-section-2-sagittal-anterior-1-standard and https://www.10xgenomics.com/datasets/mouse-brain-serial-section-2-sagittal-posterior-1-standard, respectively. The SPOTS mouse spleen data^11^ is accessible at GEO with accession code GSE198353. The 10x Visium RNA & protein human lymph node dataset^35^ can be downloaded from the Zenodo database https://zenodo.org/records/10362607. The MISAR-seq mouse brain data^12^ is available from the National Genomics Data Center via accession number OEP003285.

## Code availability

The code as well as the detailed tutorials for implementing PRESENT can be found at the Github website https://github.com/lizhen18THU/PRESENT.

## Acknowledgements

This work was partly supported by the National Key Research and Development Program of China (grant nos. 2022YFF1202400 and 2023YFF1204802), the National Natural Science Foundation of China (grant nos. 62273194), and Beijing Natural Science Foundation (L242026).

## Author contributions

R.J. and L.Z. conceived the study and supervised the project. Z.L. designed, implemented and validated the PRESENT framework. X.J.C., X.Y.C., Z.G., Y.L., Y.P., S.C. and H.L. helped with analyzing the results. Z.L. and R.J. wrote the manuscript, with input from all the authors.

