## Supplementary materials for "Cross-modality representation and multi-sample integration of spatially resolved omics data"

### Supplementary Figures

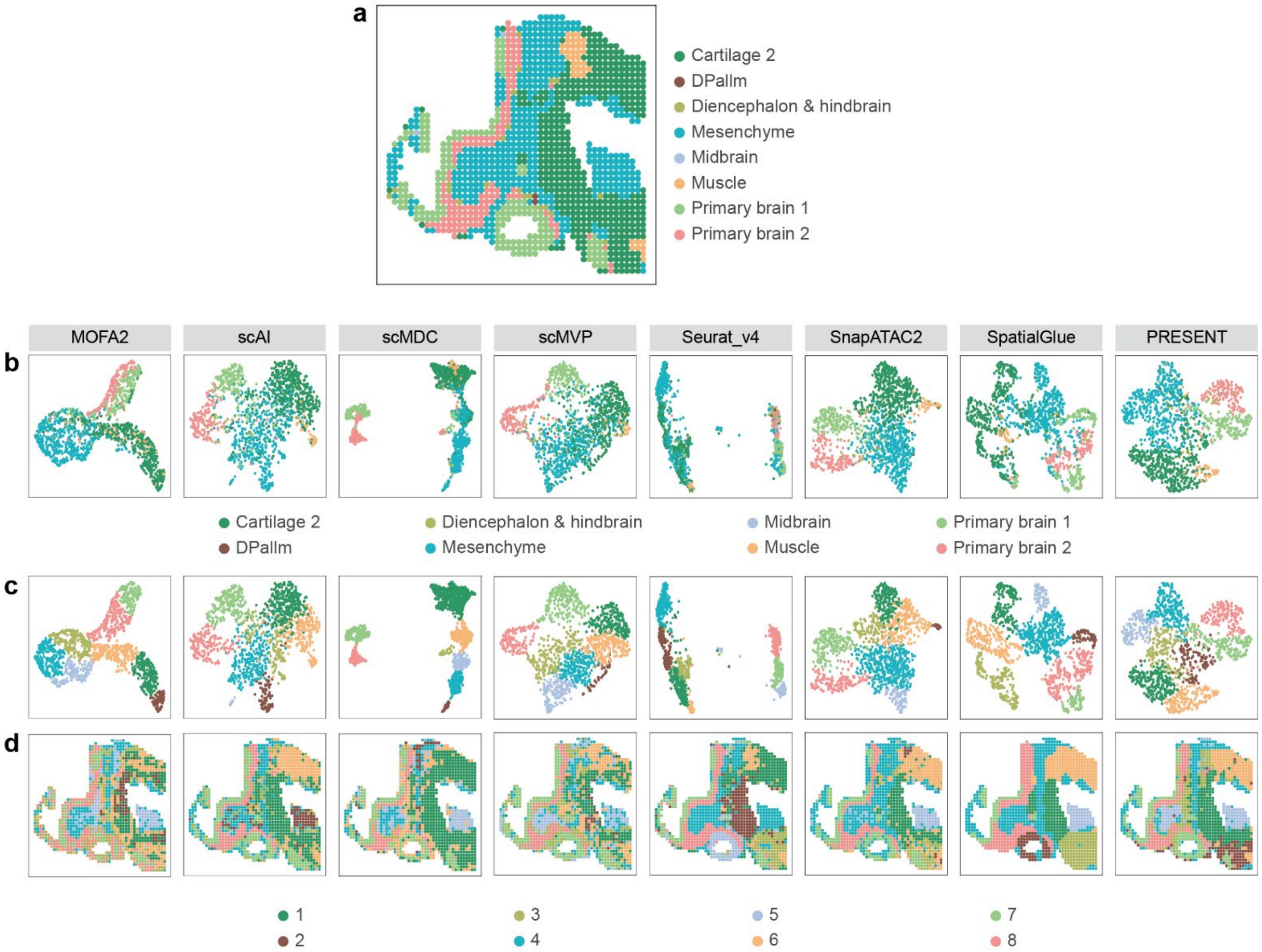

**Supplementary Fig. 1 | The UMAP and spatial visualization of spots in MISAR-seq E11.0 mouse brain sample. a,** The spatial visualization colored by ground truth domain labels, **b,** the UMAP comparison of latent representations obtained by different methods colored by ground truth domain labels, **c,** the UMAP comparison of latent representations obtained by different methods colored by spatial clusters identified based on different methods and **d,** the spatial visualization colored by spatial clusters identified based on different methods on the E11.0 mouse brain sample.

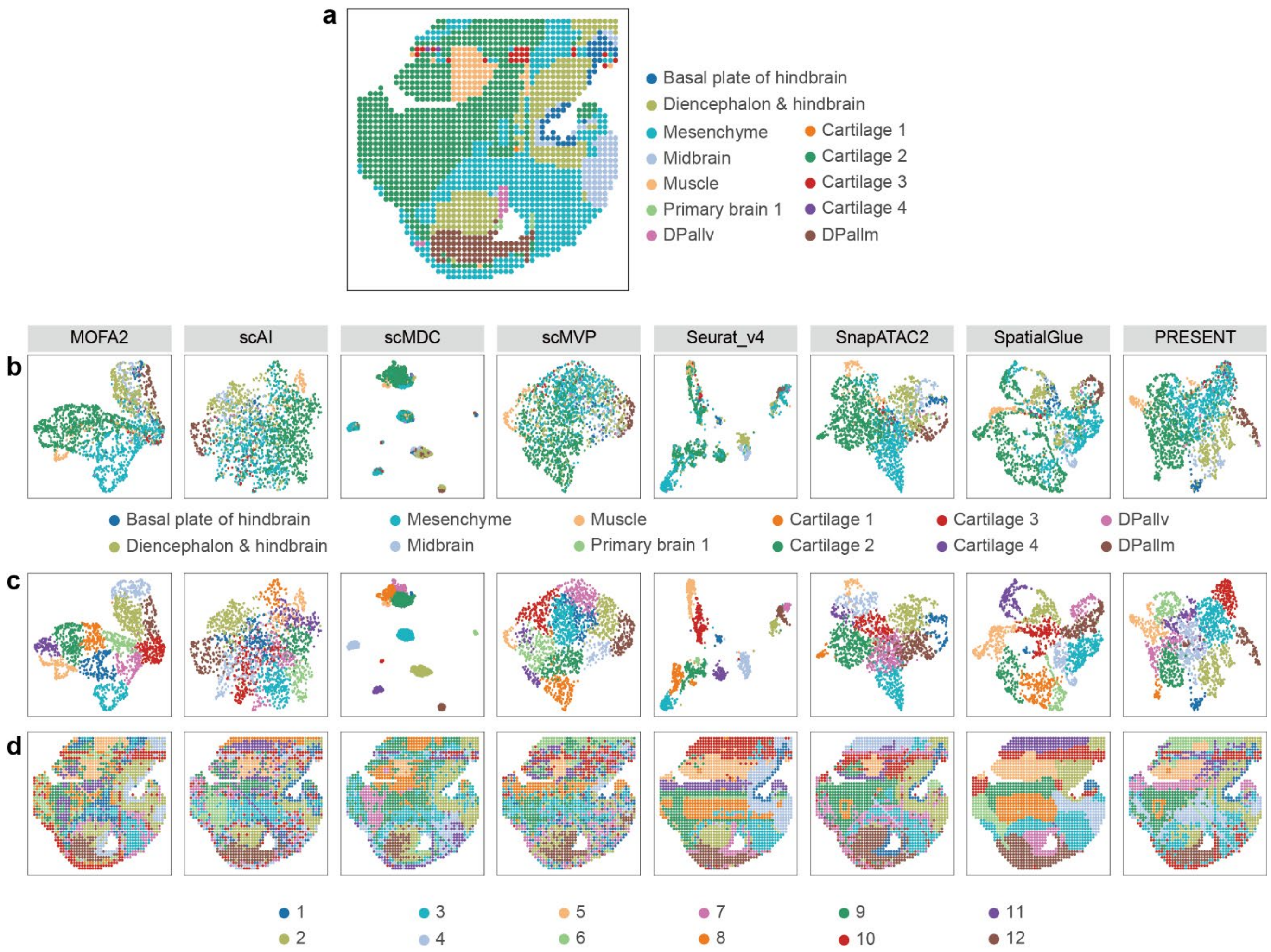

**Supplementary Fig. 2 | The UMAP and spatial visualization of spots in MISAR-seq E13.5 mouse brain sample. a,** The spatial visualization colored by ground truth domain labels, **b,** the UMAP comparison of latent representations obtained by different methods colored by ground truth domain labels, **c,** the UMAP comparison of latent representations obtained by different methods colored by spatial clusters identified based on different methods and **d,** the spatial visualization colored by spatial clusters identified based on different methods on the E13.5 mouse brain sample.

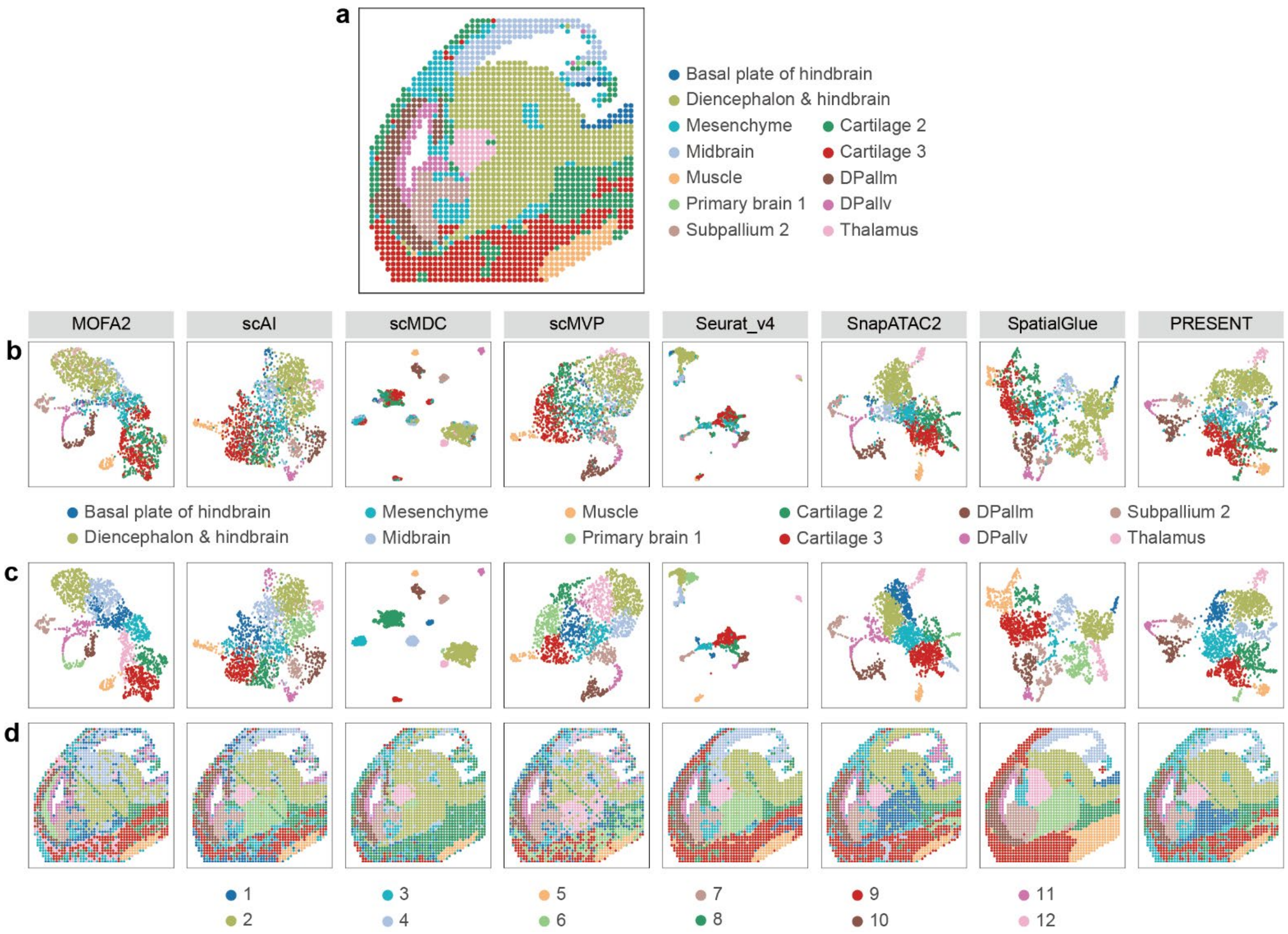

**Supplementary Fig. 3 | The UMAP and spatial visualization of spots in MISAR-seq E15.5 mouse brain sample. a,** The spatial visualization colored by ground truth domain labels, **b,** the UMAP comparison of latent representations obtained by different methods colored by ground truth domain labels, **c,** the UMAP comparison of latent representations obtained by different methods colored by spatial clusters identified based on different methods and **d,** the spatial visualization colored by spatial clusters identified based on different methods on the E15.5 mouse brain sample.

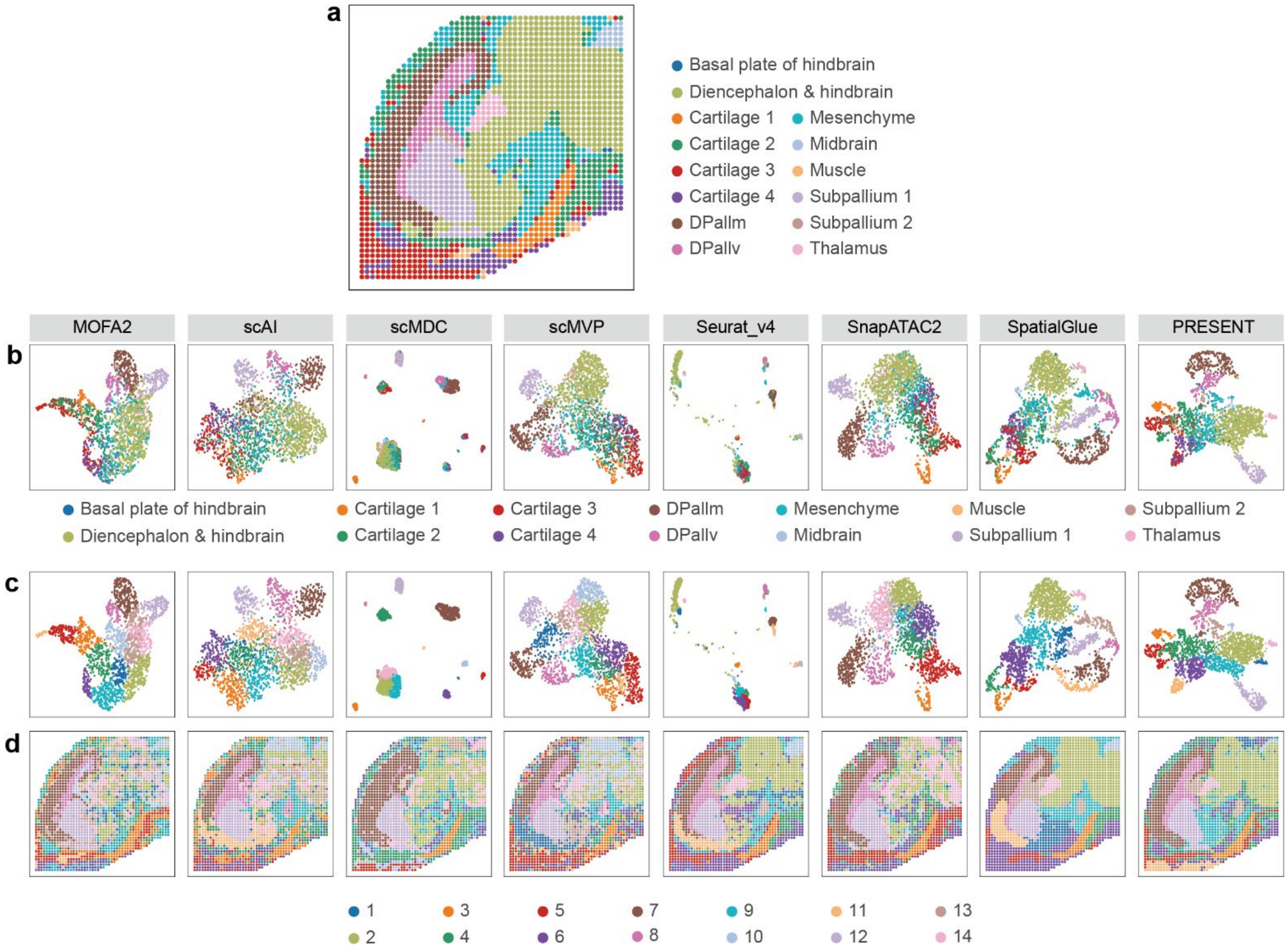

**Supplementary Fig. 4 | The UMAP and spatial visualization of spots in MISAR-seq E18.5 mouse brain sample. a,** The spatial visualization colored by ground truth domain labels, **b,** the UMAP comparison of latent representations obtained by different methods colored by ground truth domain labels, **c,** the UMAP comparison of latent representations obtained by different methods colored by spatial clusters identified based on different methods and **d,** the spatial visualization colored by spatial clusters identified based on different methods on the E18.5 mouse brain sample.

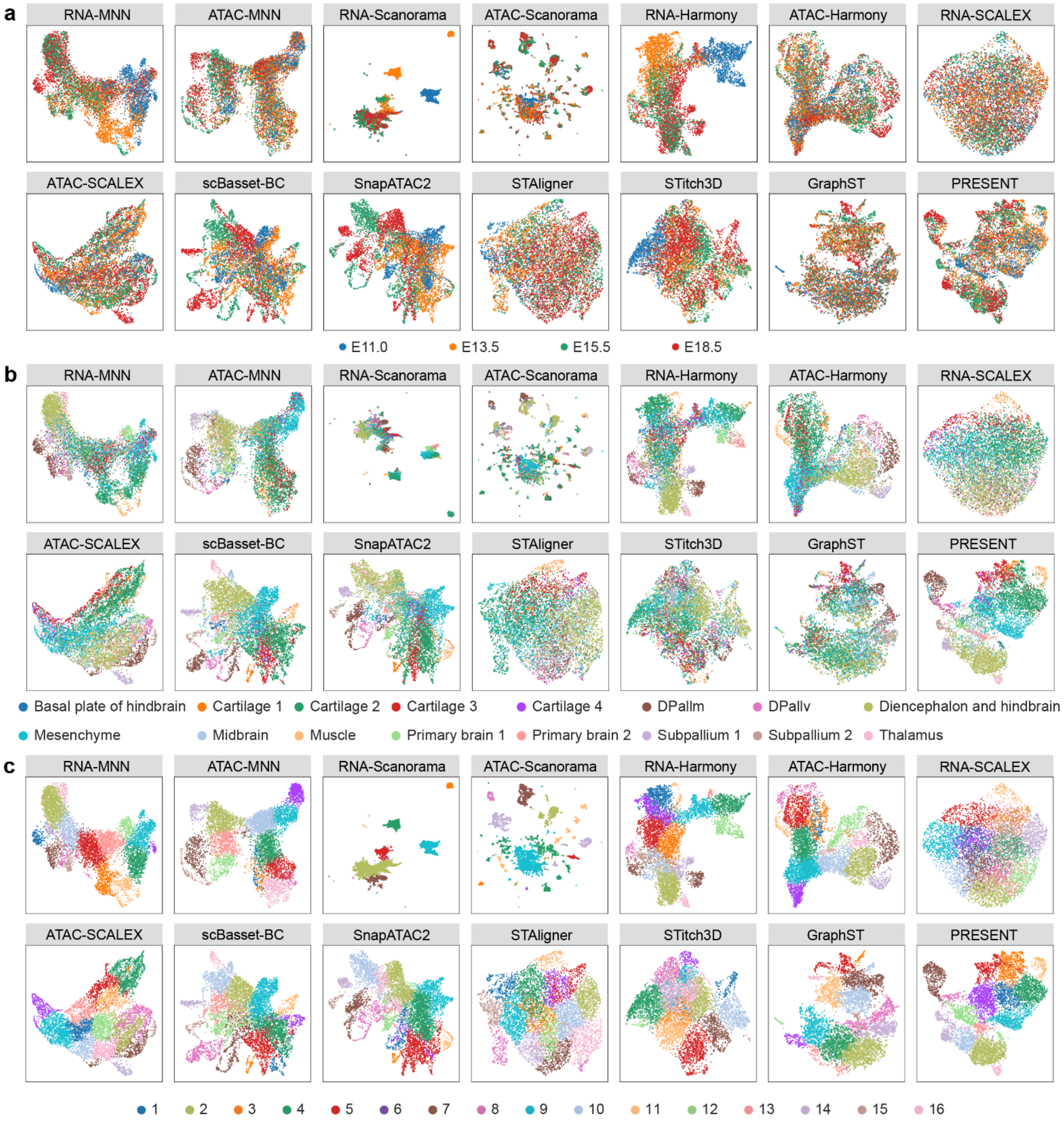

**Supplementary Fig. 5 | The joint UMAP visualization of spots in MISAR-seq mouse brain samples.** The UMAP visualization of latent embeddings obtained by different methods colored by **a,** sample indices, **b,** ground truth domain labels and **c,** joint spatial cluster labels obtained based on different methods on the four MISAR-seq mouse brain samples from E11.0, E13.5, E15.5 and E18.5 developmental stages.

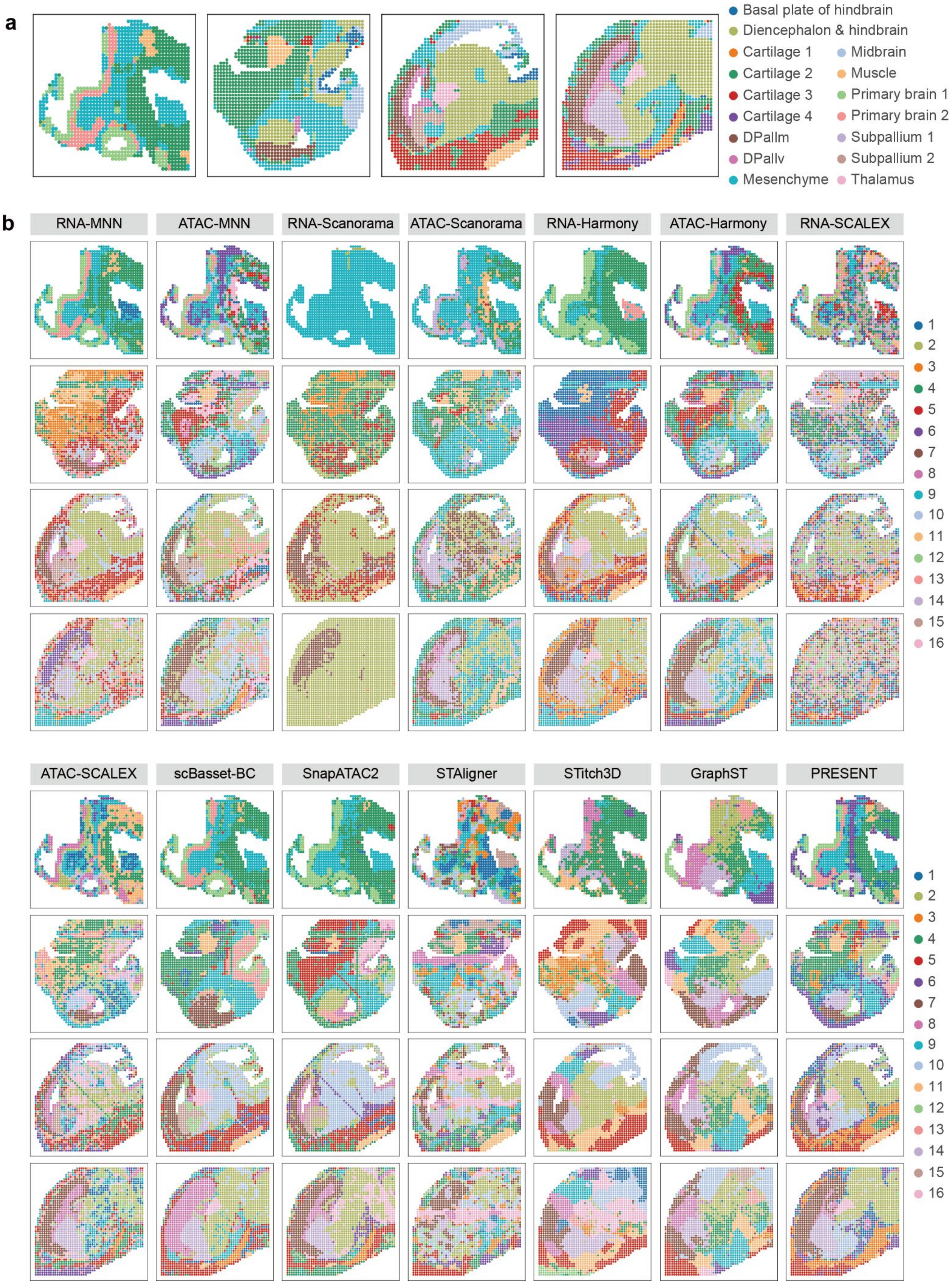

**Supplementary Fig. 6 | The spatial visualization of spots in MISAR-seq mouse brain samples.** The spatial visualization colored by **a,** ground truth domains and **b,** joint clustering labels obtained based on different methods.

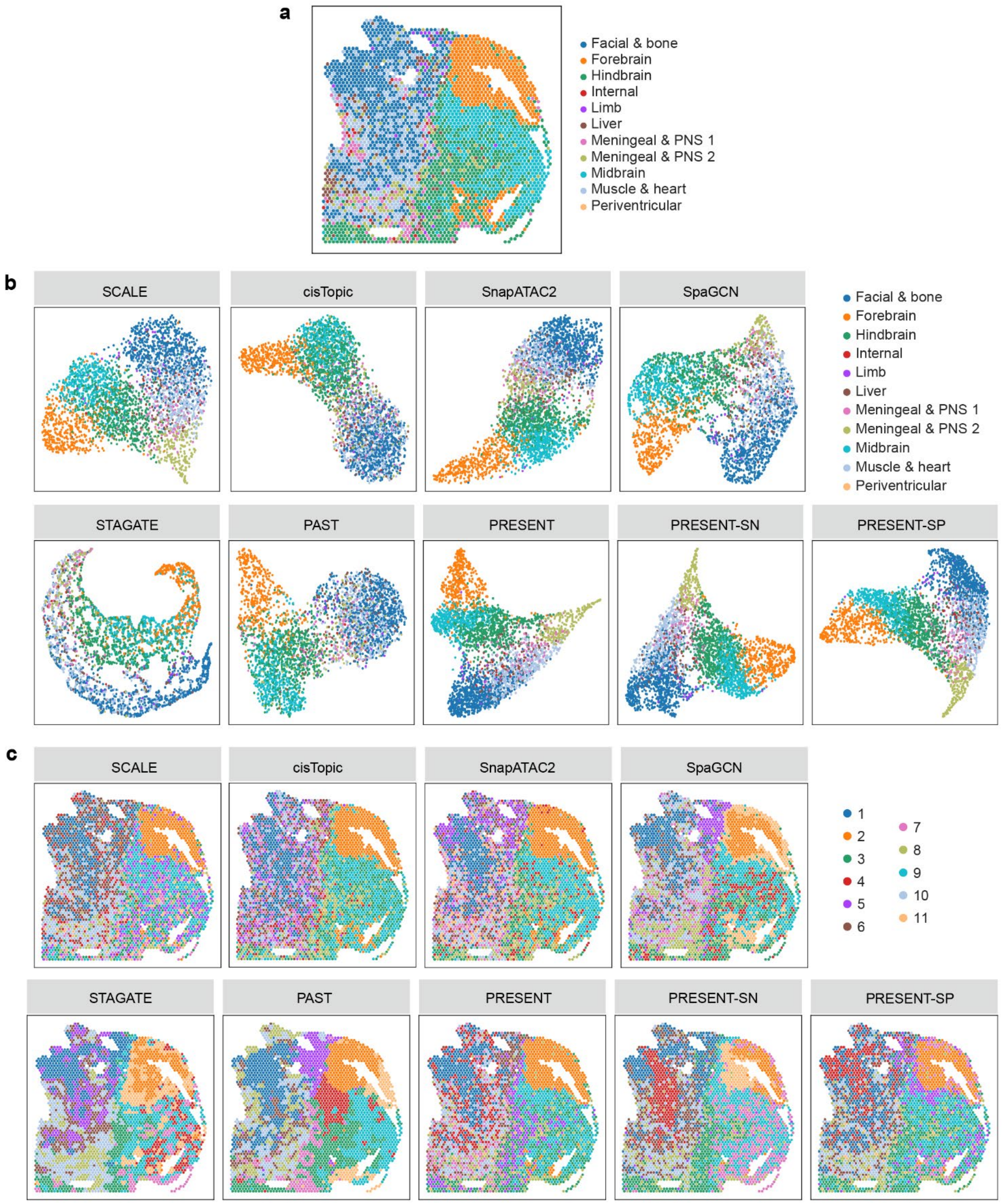

**Supplementary Fig. 7 | The UMAP and spatial visualization of spots in spatial ATAC E15.5-S1 mouse embryo sample. a,** The spatial visualization colored by ground truth domain labels, **b,** the UMAP comparison of latent representations obtained by different methods colored by ground truth domain labels and **c,** the spatial visualization colored by spatial clusters identified based on different methods on the E15.5-S1 mouse embryo sample.

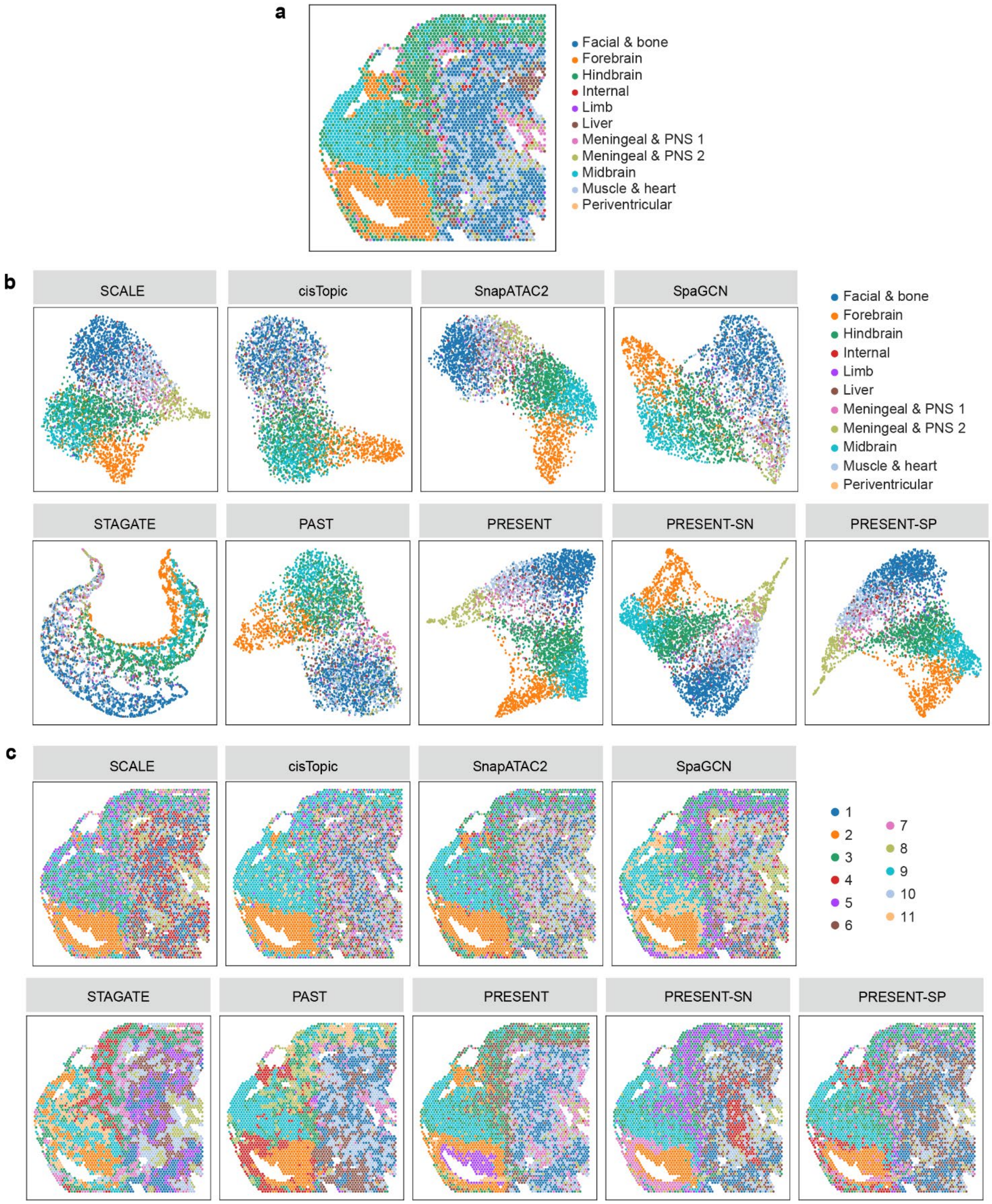

**Supplementary Fig. 8 | The UMAP and spatial visualization of spots in spatial ATAC E15.5-S2 mouse embryo sample. a,** The spatial visualization colored by ground truth domain labels, **b,** the UMAP comparison of latent representations obtained by different methods colored by ground truth domain labels and **c,** the spatial visualization colored by spatial clusters identified based on different methods on the E15.5-S2 mouse embryo sample.

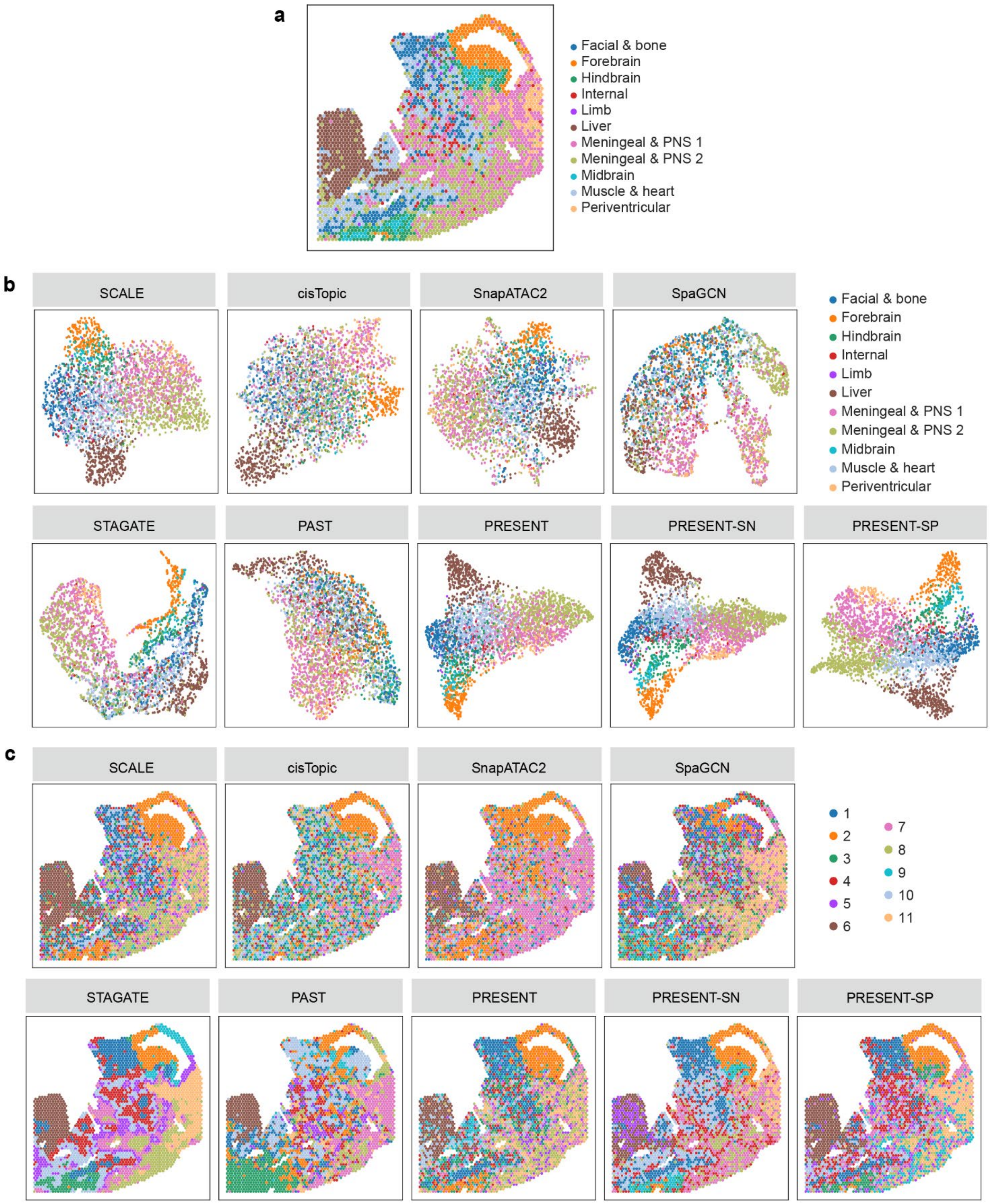

**Supplementary Fig. 9 | The UMAP and spatial visualization of spots in spatial ATAC E13.5-S1 mouse embryo sample. a,** The spatial visualization colored by ground truth domain labels, **b,** the UMAP comparison of latent representations obtained by different methods colored by ground truth domain labels and **c,** the spatial visualization colored by spatial clusters identified based on different methods on the E13.5-S1 mouse embryo sample.

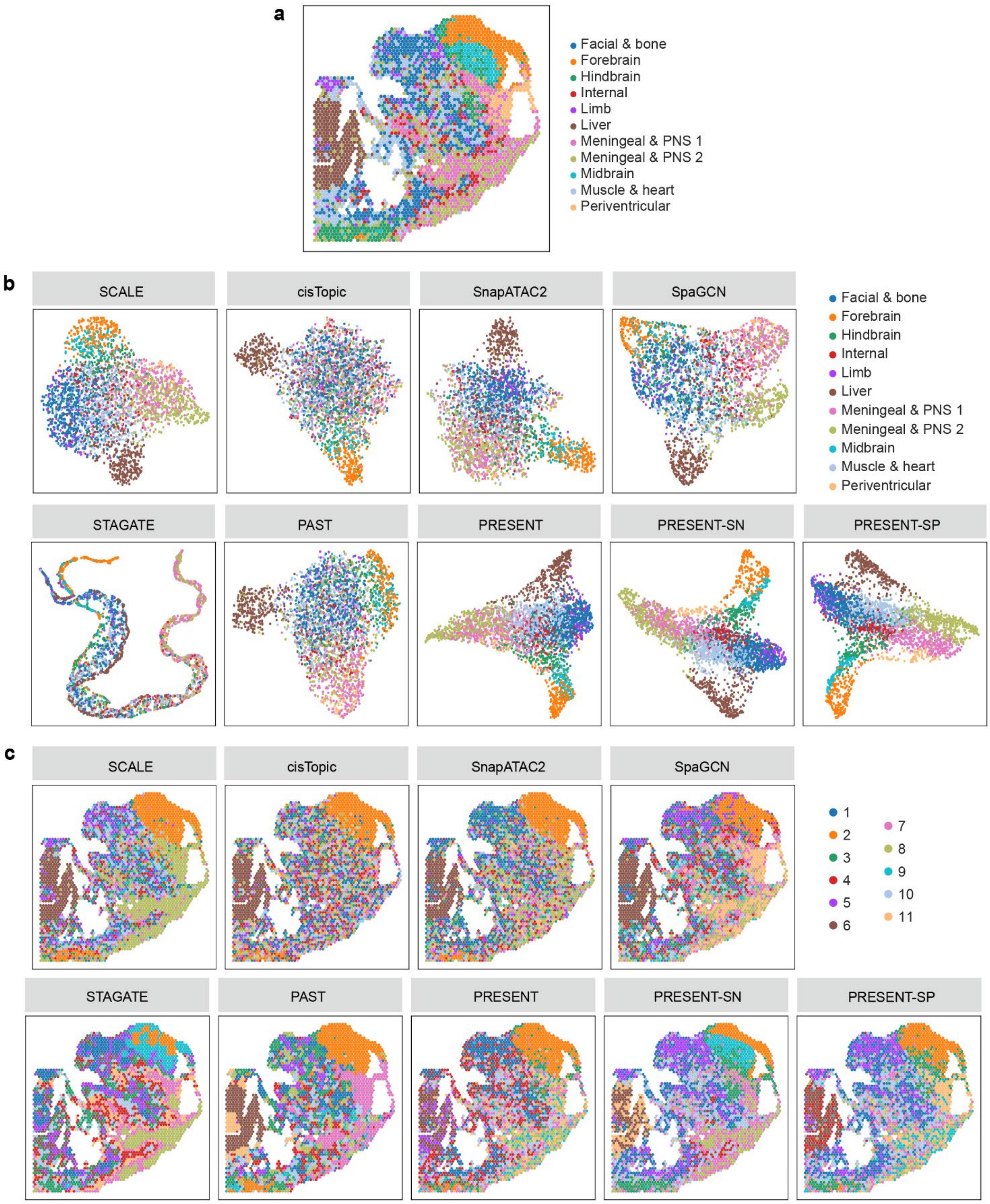

**Supplementary Fig. 10 | The UMAP and spatial visualization of spots in spatial ATAC E13.5-S2 mouse embryo sample. a,** The spatial visualization colored by ground truth domain labels, **b,** the UMAP comparison of latent representations obtained by different methods colored by ground truth domain labels and **c,** the spatial visualization colored by spatial clusters identified based on different methods on the E13.5-S2 mouse embryo sample.

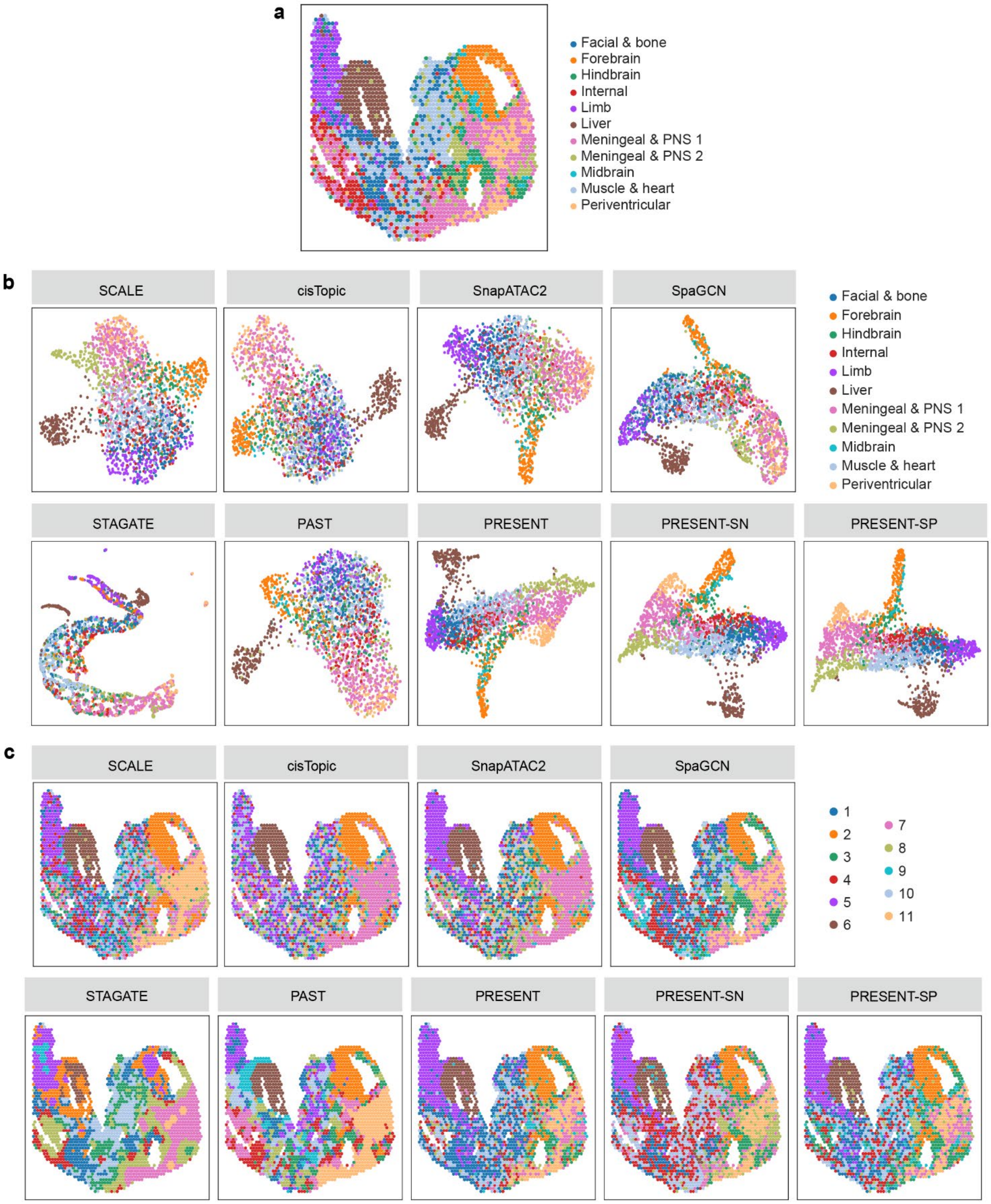

**Supplementary Fig. 11 | The UMAP and spatial visualization of spots in spatial ATAC E12.5-S1 mouse embryo sample. a,** The spatial visualization colored by ground truth domain labels, **b,** the UMAP comparison of latent representations obtained by different methods colored by ground truth domain labels and **c,** the spatial visualization colored by spatial clusters identified based on different methods on the E12.5-S1 mouse embryo sample.

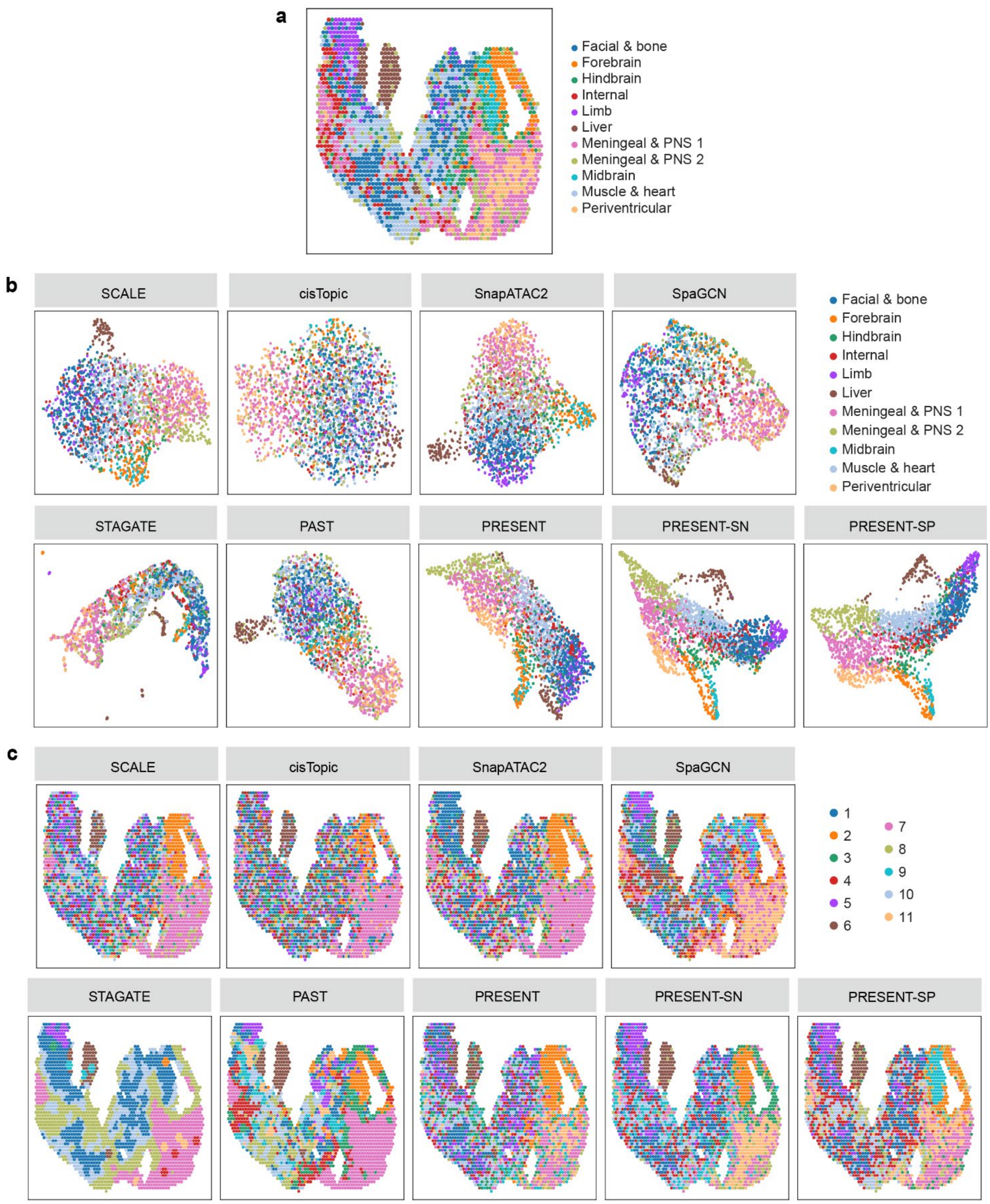

**Supplementary Fig. 12 | The UMAP and spatial visualization of spots in spatial ATAC E12.5-S2 mouse embryo sample. a,** The spatial visualization colored by ground truth domain labels, **b,** the UMAP comparison of latent representations obtained by different methods colored by ground truth domain labels and **c,** the spatial visualization colored by spatial clusters identified based on different methods on the E12.5-S2 mouse embryo sample.

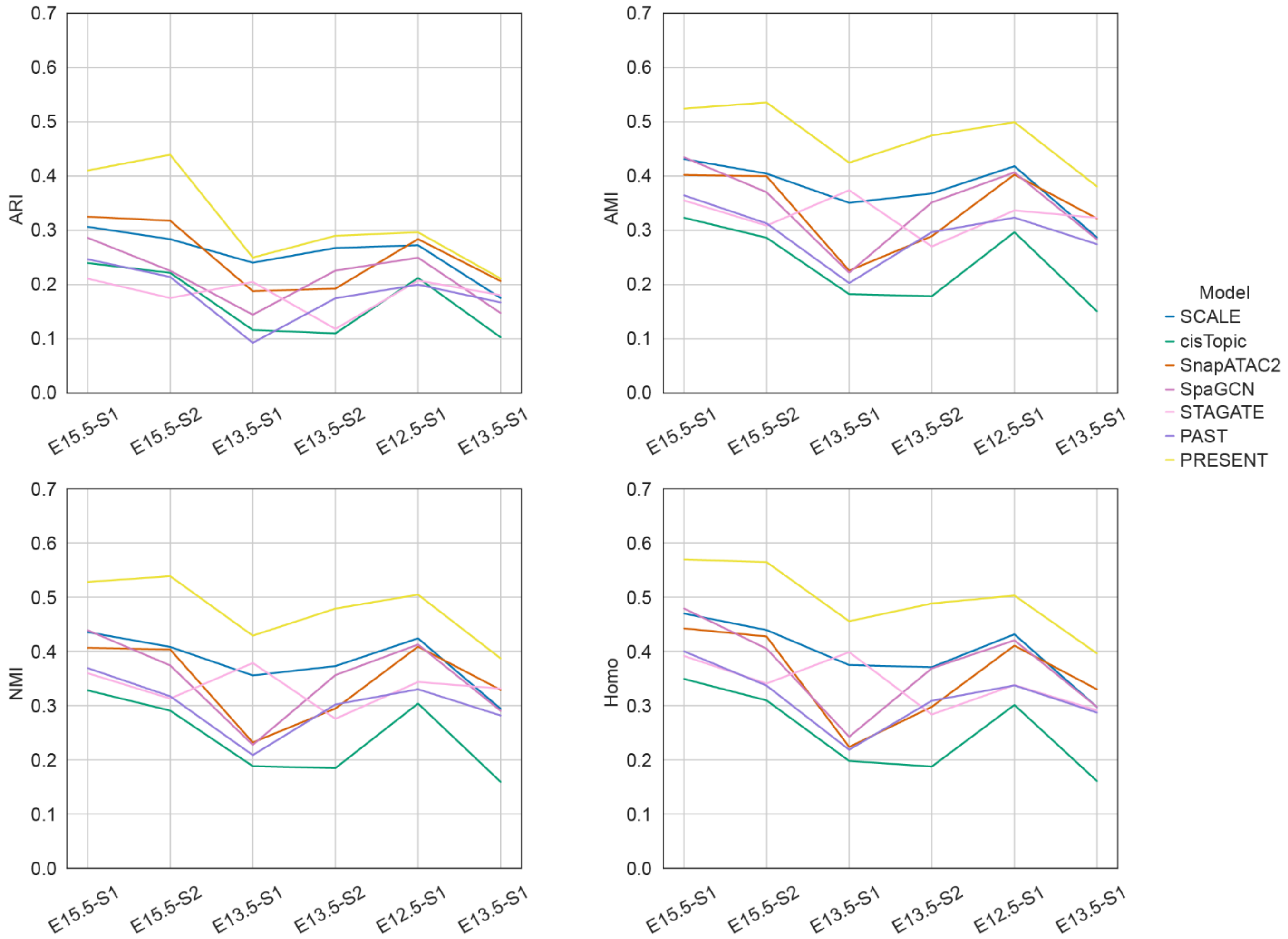

**Supplementary Fig. 13 | The detailed evaluation of different methods on the six spatial ATAC mouse embryo samples.** The detailed benchmarking results on PRESENT and other baseline methods using four metrics, namely ARI, AMI, NMI and Homo, on the six mouse embryo samples, including E15.5-S1, E15.5-S2, E13.5-S1, E13.5-S2, E12.5-S1 and E12.5-S2.

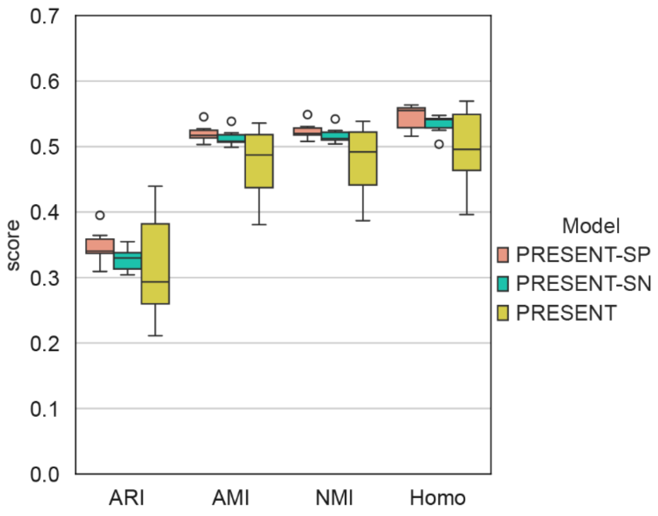

**Supplementary Fig. 14 | The comparison of PRESENT with or without reference data on the six spatial ATAC mouse embryo samples.** PRESENT using the single-nucleus ATAC-seq dataset as reference data is denoted as PRESENT-SN, while PRESENT using the other five spatial ATAC samples as reference is denoted as PRESENT-SP.

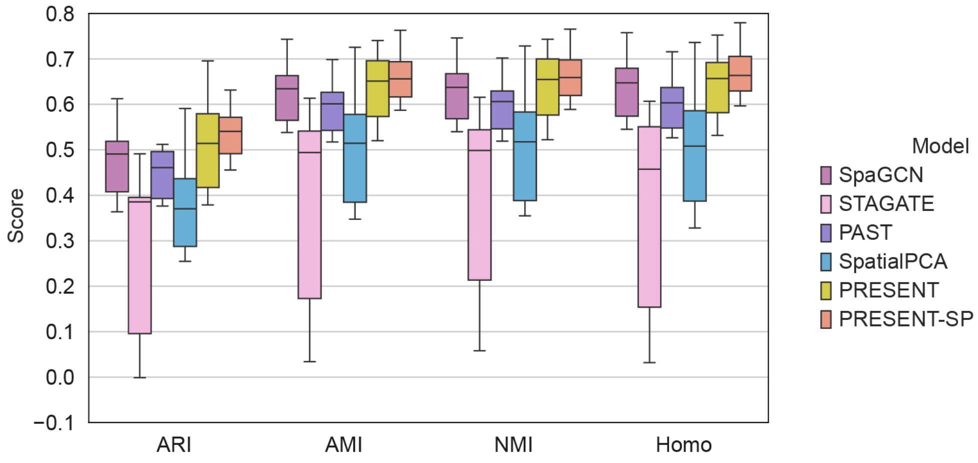

**Supplementary Fig. 15 | The evaluation of different methods on the six Stereo-seq axolotl brain samples.** PRESENT using the regenerated axolotl brain data at 20 days post-injury as reference data is denoted as PRESENT-SP.

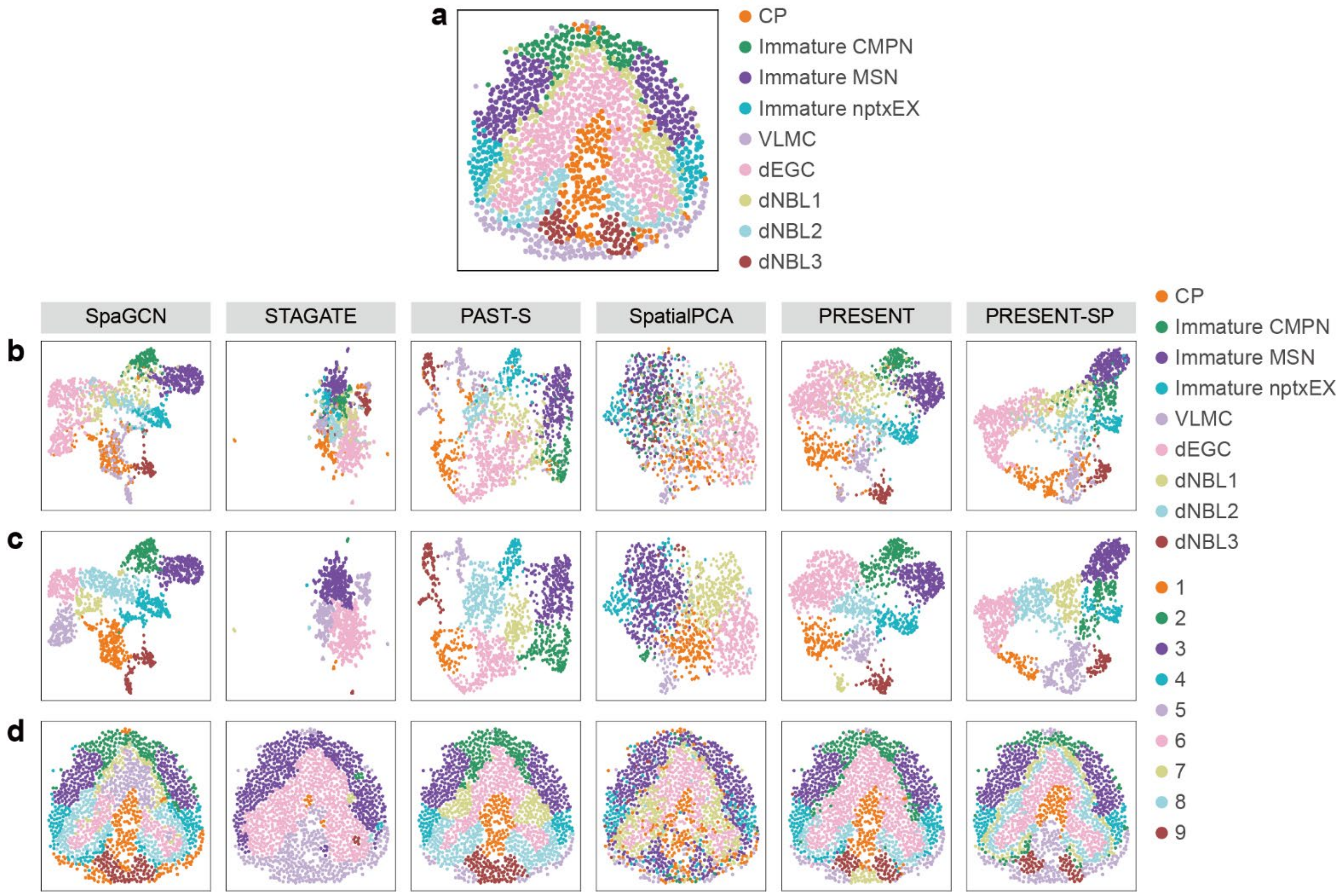

**Supplementary Fig. 16 | The UMAP and spatial visualization of spots in Stereo-seq stage 44 axolotl brain sample.** **a,** The spatial visualization colored by ground truth domain labels, **b,** the UMAP comparison of latent representations obtained by different methods colored by ground truth domain labels, **c,** the UMAP comparison of latent representations obtained by different methods colored by spatial clusters identified based on different methods and **d,** the spatial visualization colored by spatial clusters identified based on different methods on the stage 44 axolotl brain sample. CP, choroid plexus; CMPN, cholinergic, monoaminergic, and peptidergic neuron; MSN, medium spiny neuron; nptxEX, $Nptx^{+}$ lateral pallium excitatory neuron; VLMC, vascular leptomeningeal cell; dEGC, developmental ependymoglial cell; dNBL, developmental neuroblast.

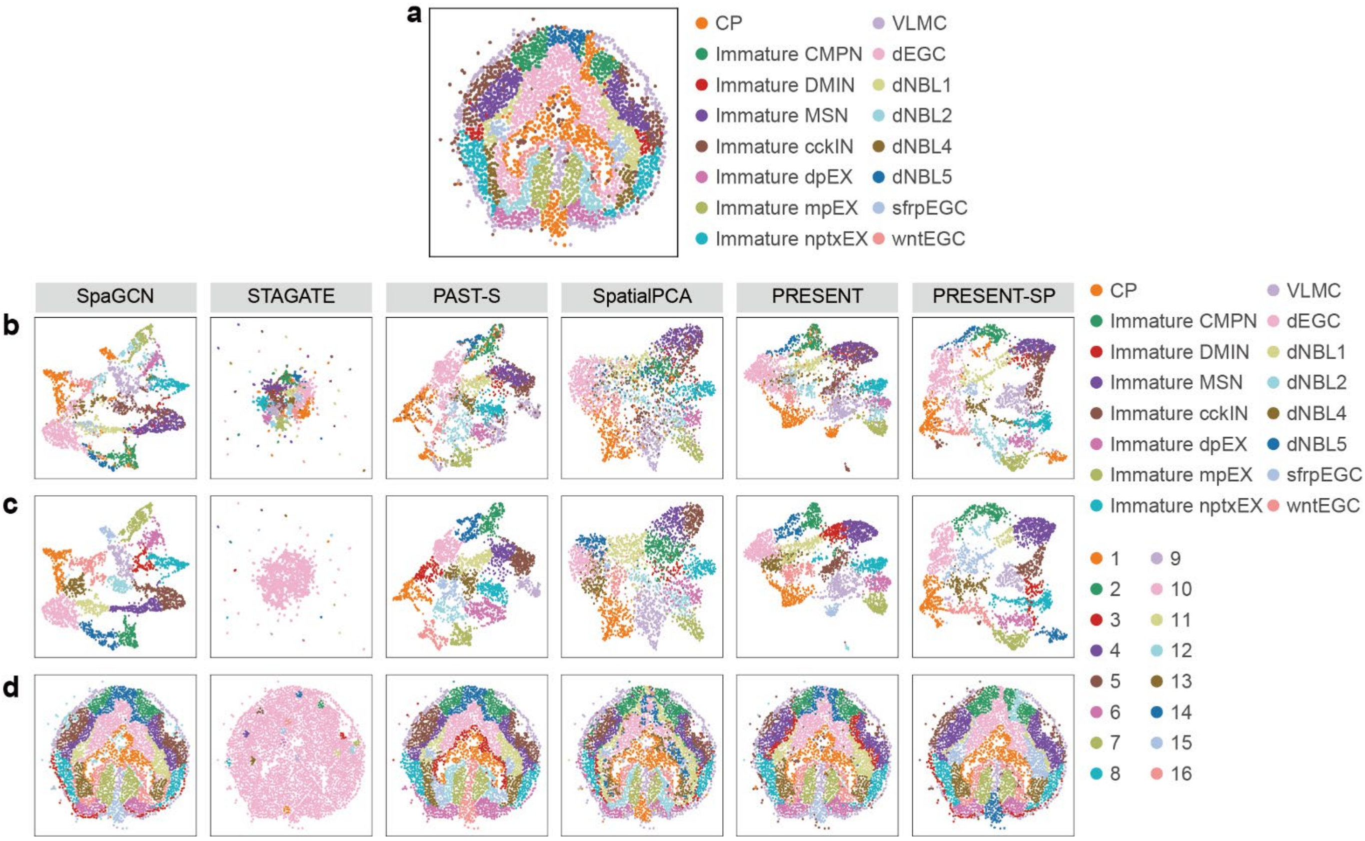

**Supplementary Fig. 17 | The UMAP and spatial visualization of spots in Stereo-seq stage 54 axolotl brain sample.** **a,** The spatial visualization colored by ground truth domain labels, **b,** the UMAP comparison of latent representations obtained by different methods colored by ground truth domain labels, **c,** the UMAP comparison of latent representations obtained by different methods colored by spatial clusters identified based on different methods and **d,** the spatial visualization colored by spatial clusters identified based on different methods on the stage 54 axolotl brain sample. CP, choroid plexus; CMPN, cholinergic, monoaminergic, and peptidergic neuron; DMIN, dopaminergic periglomerular inhibitory neuron; MSN, medium spiny neuron; cckIN, Cck$^{+}$ inhibitory neuron; dpEX, dorsal palliumexcitatory neuron; mpEX, medial pallium excitatory neuron; nptxEX, Nptx$^{+}$ lateral pallium excitatory neuron; VLMC, vascular leptomeningeal cell; dEGC, developmental ependymoglial cell; dNBL, developmental neuroblast; sfrpEGC, Sfrp$^{+}$ ependymoglial cell; wntEGC, Wnt$^{+}$ ependymoglial cell.

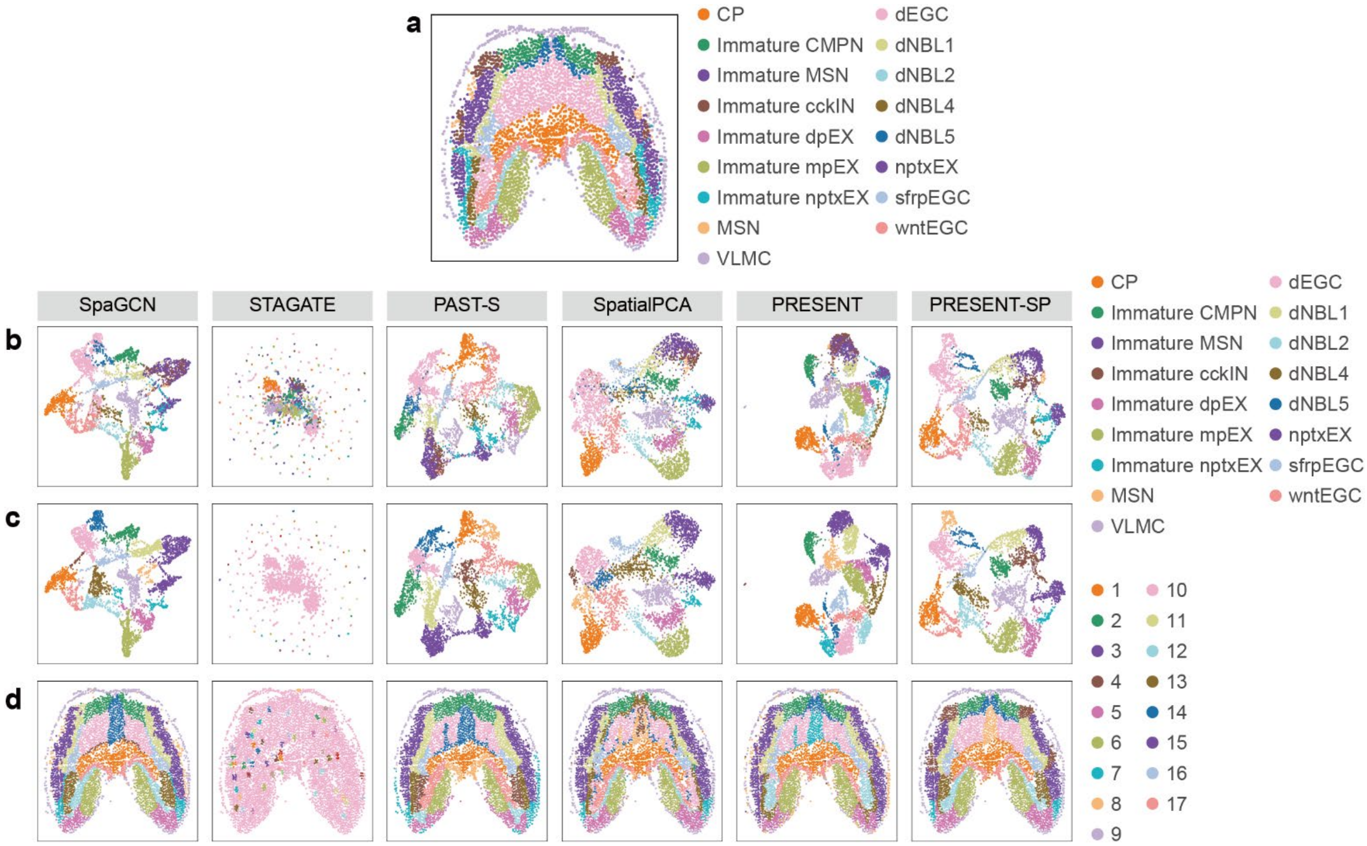

**Supplementary Fig. 18 | The UMAP and spatial visualization of spots in Stereo-seq stage 57 axolotl brain sample.** **a,** The spatial visualization colored by ground truth domain labels, **b,** the UMAP comparison of latent representations obtained by different methods colored by ground truth domain labels, **c,** the UMAP comparison of latent representations obtained by different methods colored by spatial clusters identified based on different methods and **d,** the spatial visualization colored by spatial clusters identified based on different methods on the stage 57 axolotl brain sample. CP, choroid plexus; CMPN, cholinergic, monoaminergic, and peptidergic neuron; MSN, medium spiny neuron; cckIN, Cck$^{+}$ inhibitory neuron; dpEX, dorsal palliumexcitatory neuron; mpEX, medial pallium excitatory neuron; nptxEX, Nptx$^{+}$ lateral pallium excitatory neuron; VLMC, vascular leptomeningeal cell; dEGC, developmental ependymoglial cell; dNBL, developmental neuroblast; sfrpEGC, Sfrp$^{+}$ ependymoglial cell; wntEGC, Wnt$^{+}$ ependymoglial cell.

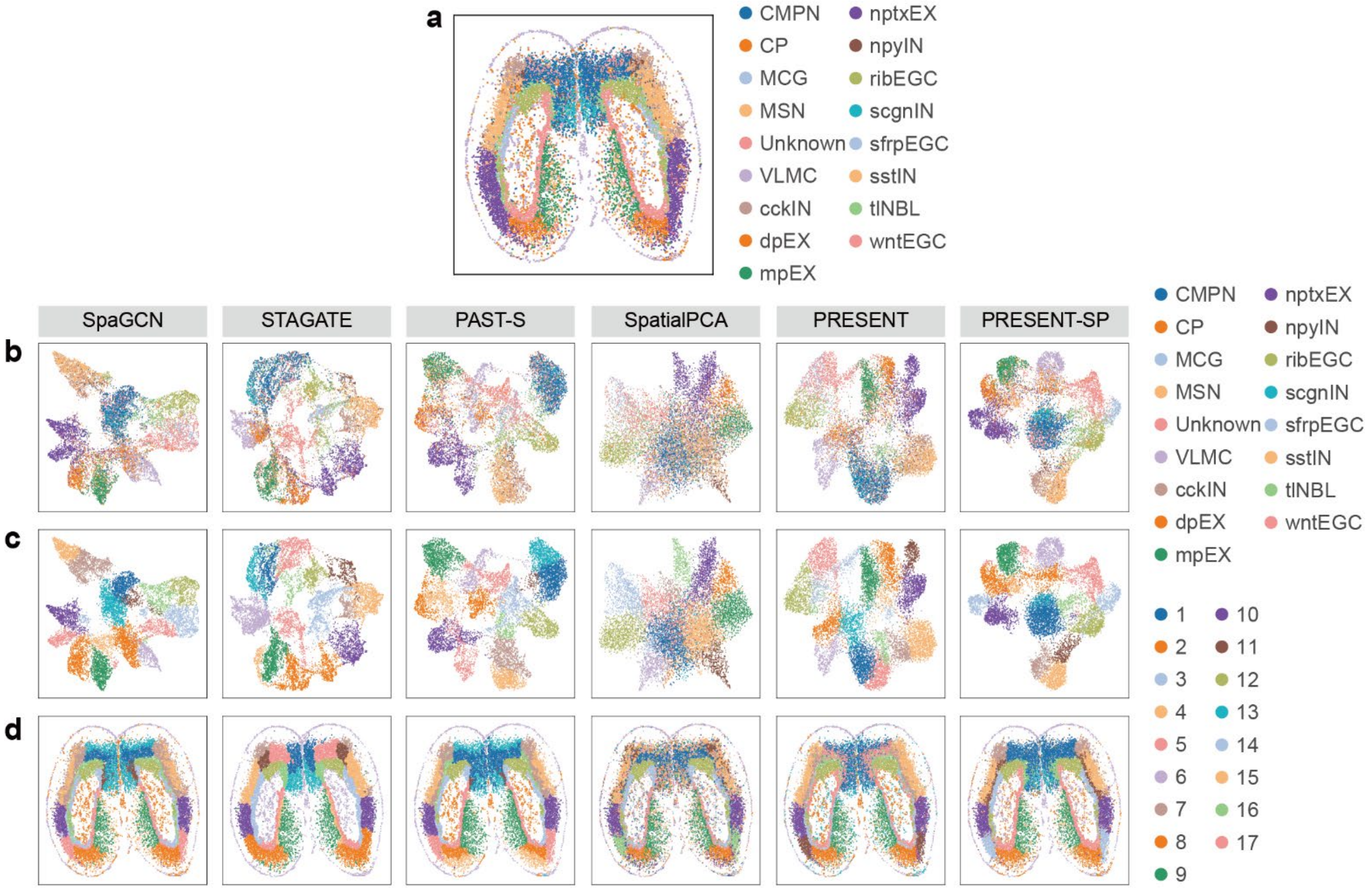

**Supplementary Fig. 19 | The UMAP and spatial visualization of spots in Stereo-seq juvenile axolotl brain sample.** **a,** The spatial visualization colored by ground truth domain labels, **b,** the UMAP comparison of latent representations obtained by different methods colored by ground truth domain labels, **c,** the UMAP comparison of latent representations obtained by different methods colored by spatial clusters identified based on different methods and **d,** the spatial visualization colored by spatial clusters identified based on different methods on the juvenile axolotl brain sample. CMPN, cholinergic, monoaminergic, and peptidergic neuron; CP, choroid plexus; MCG, microglial cell; MSN, medium spiny neuron; VLMC, vascular leptomeningeal cell; cckIN, Cck$^{+}$ inhibitory neuron; dpEX, dorsal palliumexcitatory neuron; mpEX, medial pallium excitatory neuron; nptxEX, Nptx$^{+}$ lateral pallium excitatory neuron; npyIN, Npy$^{+}$ inhibitory neuron; ribEGC, ribosomal ependymoglial cell; scgnIN, Scgn$^{+}$ inhibitory neuron; sfrpEGC, Sfrp$^{+}$ ependymoglial cell; sstIN, Sst$^{+}$ inhibitory neuron; tlNBL, telencephalon neuroblast; wntEGC, Wnt$^{+}$ ependymoglial cell.

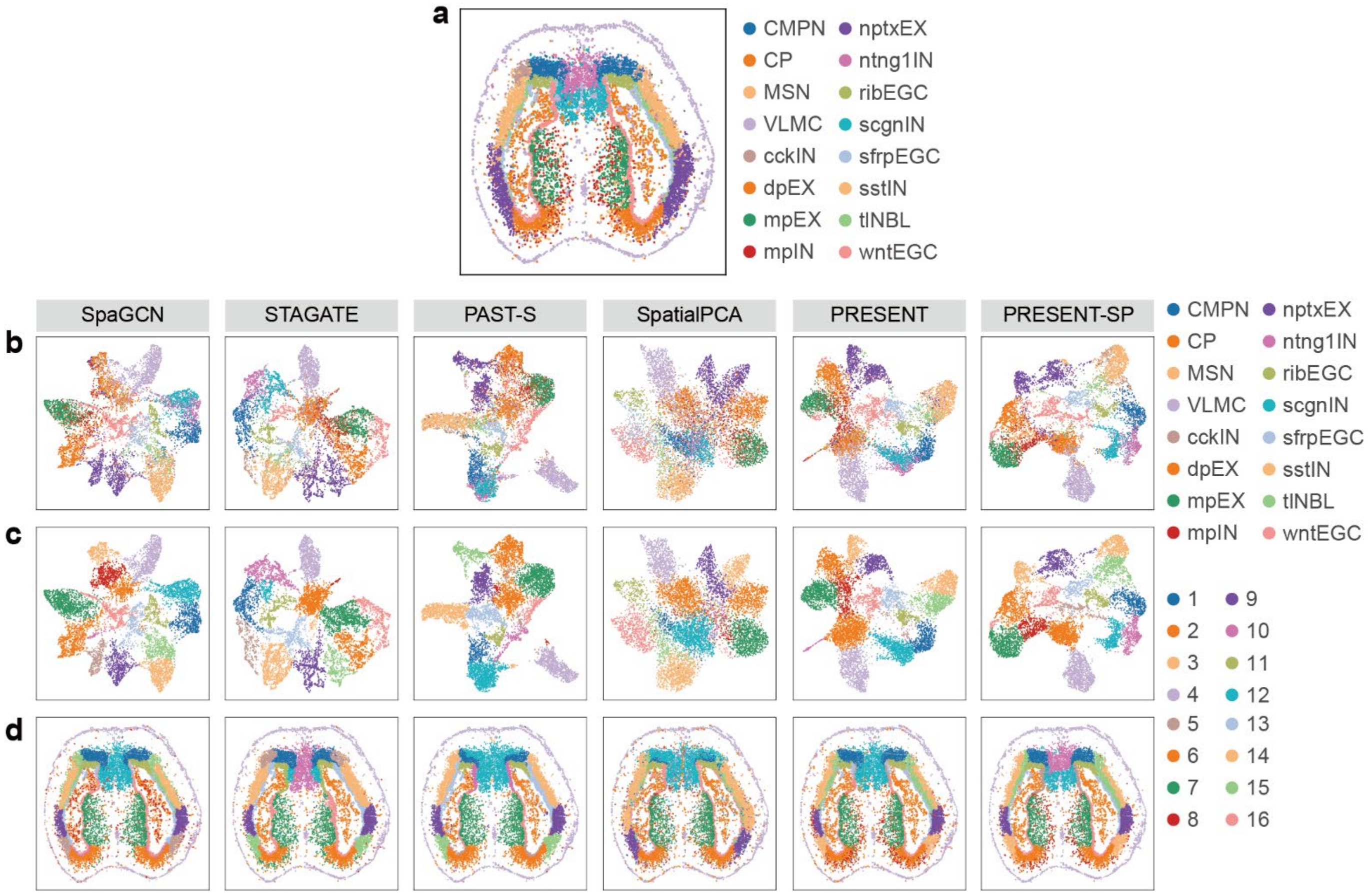

**Supplementary Fig. 20 | The UMAP and spatial visualization of spots in Stereo-seq adult axolotl brain sample. a,** The spatial visualization colored by ground truth domain labels, **b,** the UMAP comparison of latent representations obtained by different methods colored by ground truth domain labels, **c,** the UMAP comparison of latent representations obtained by different methods colored by spatial clusters identified based on different methods and **d,** the spatial visualization colored by spatial clusters identified based on different methods on the adult axolotl brain sample. CMPN, cholinergic, monoaminergic, and peptidergic neuron; CP, choroid plexus; MSN, medium spiny neuron; VLMC, vascular leptomeningeal cell; cckIN, Cck$^+$ inhibitory neuron; dpEX, dorsal palliumexcitatory neuron; mpEX, medial pallium excitatory neuron; mpIN, medial pallium inhibitory neuron; nptxEX, Nptx$^+$ lateral pallium excitatory neuron; ntng1IN, Ntng1$^+$ inhibitory neuron; ribEGC, ribosomal ependymoglial cell; scgnIN, Scgn$^+$ inhibitory neuron; sfrpEGC, Sfrp$^+$ ependymoglial cell; sstIN, Sst$^+$ inhibitory neuron; tlNBL, telencephalon neuroblast; wntEGC, Wnt$^+$ ependymoglial cell.

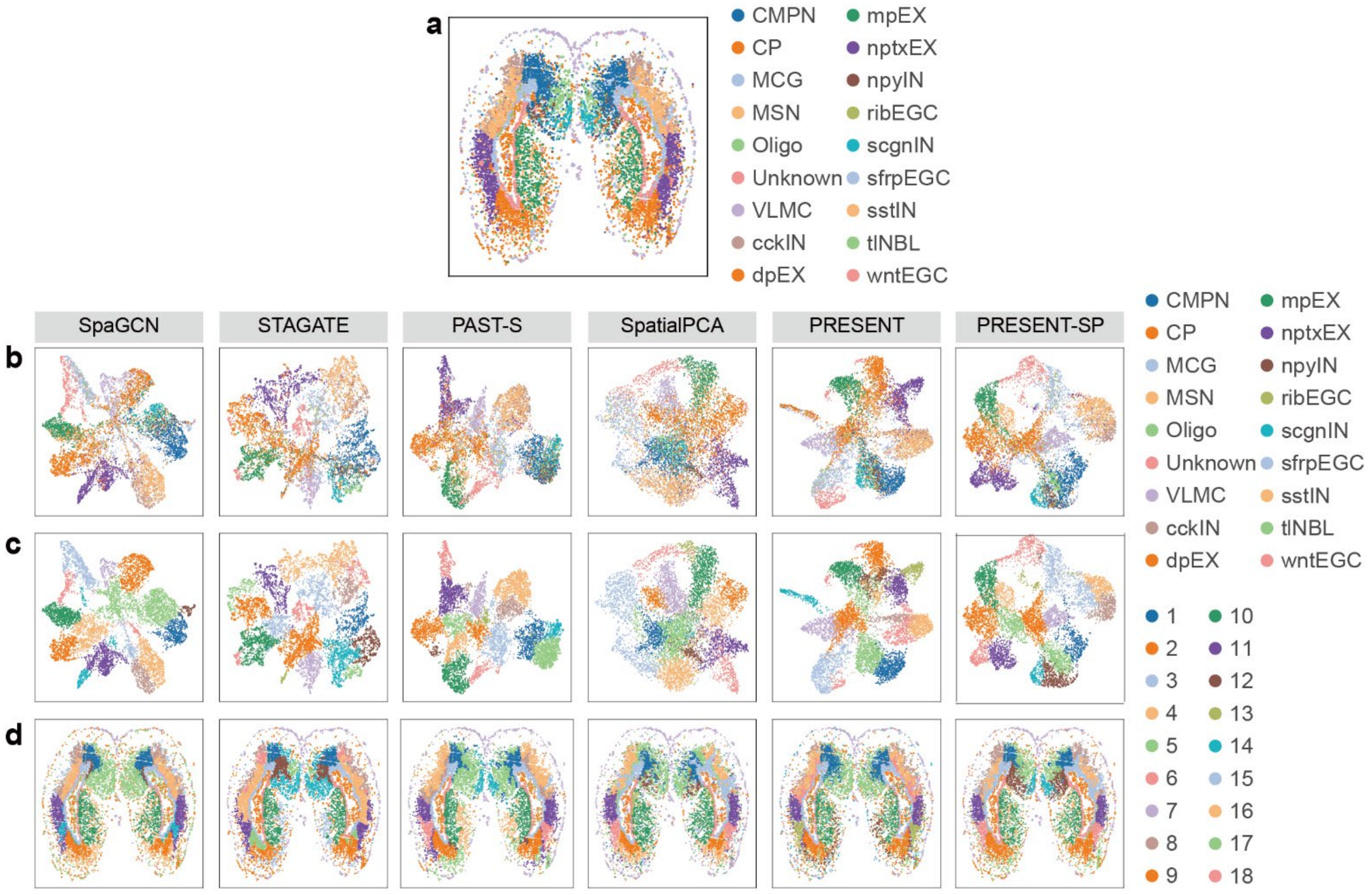

**Supplementary Fig. 21 | The UMAP and spatial visualization of spots in Stereo-seq metamorphosis axolotl brain sample. a,** The spatial visualization colored by ground truth domain labels, **b,** the UMAP comparison of latent representations obtained by different methods colored by ground truth domain labels, **c,** the UMAP comparison of latent representations obtained by different methods colored by spatial clusters identified based on different methods and **d,** the spatial visualization colored by spatial clusters identified based on different methods on the metamorphosis axolotl brain sample. CMPN, cholinergic, monoaminergic, and peptidergic neuron; CP, choroid plexus; MCG, microglial cell; MSN, medium spiny neuron; Oligo, oligodendrocyte; VLMC, vascular leptomeningeal cell; cckIN, $Cck^+$ inhibitory neuron; dpEX, dorsal palliumexcitatory neuron; mpEX, medial pallium excitatory neuron; nptxEX, $Nptx^+$ lateral pallium excitatory neuron; npyIN, $Npy^+$ inhibitory neuron; ribEGC, ribosomal ependymoglial cell; scgnIN, $Scgn^+$ inhibitory neuron; sfrpEGC, $Sfrp^+$ ependymoglial cell; sstIN, $Sst^+$ inhibitory neuron; tlNBL, telencephalon neuroblast; wntEGC, $Wnt^+$ ependymoglial cell.

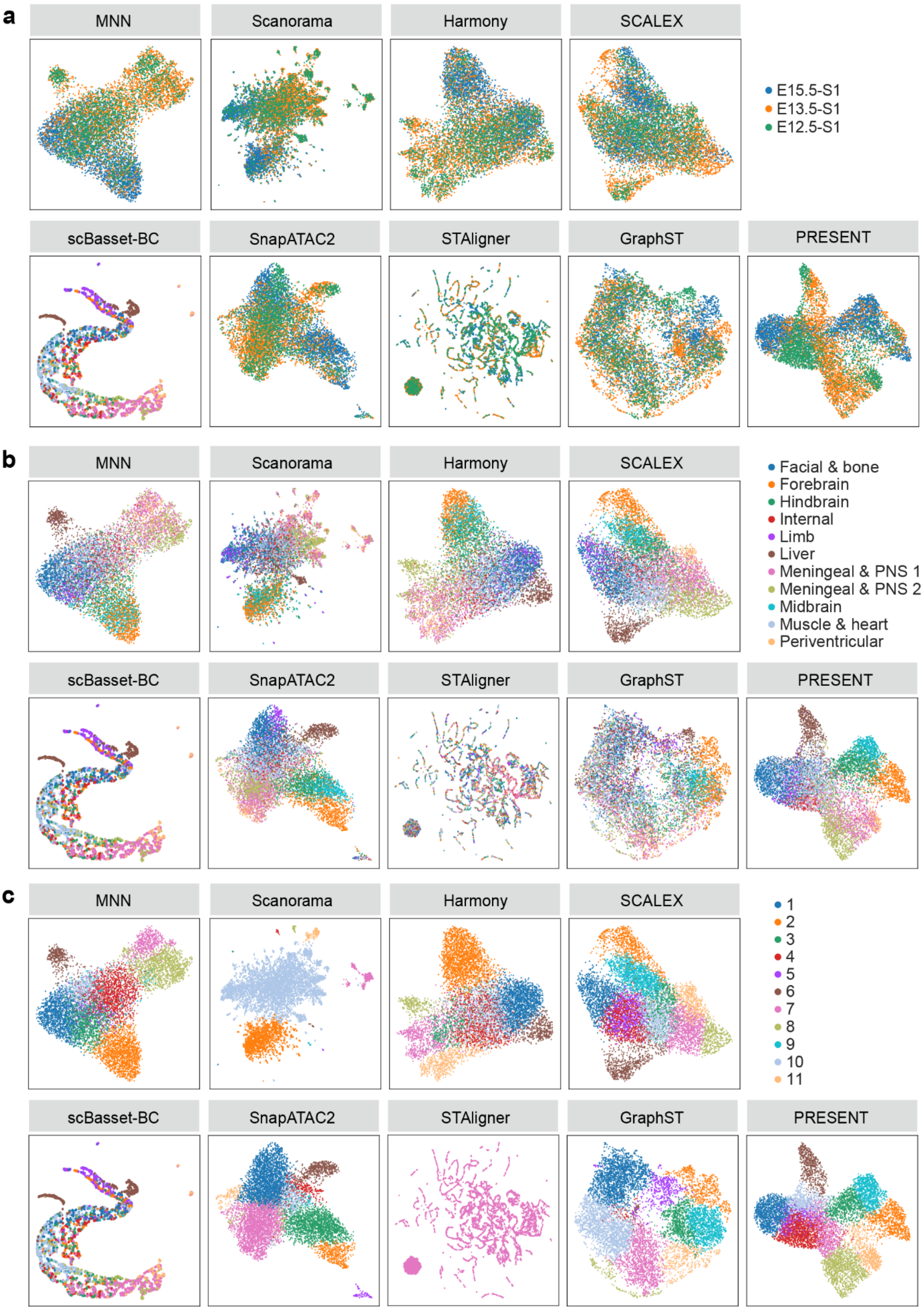

**Supplementary Fig. 22 | The joint UMAP visualization of spots in spatial ATAC mouse embryo samples.** The UMAP visualization of latent embeddings obtained by different methods colored by **a,** sample indices, **b,** ground truth domain labels, **c,** joint spatial cluster labels obtained based on different methods on the three spatial ATAC mouse embryo samples from E12.5, E13.5 and E15.5 developmental stages.

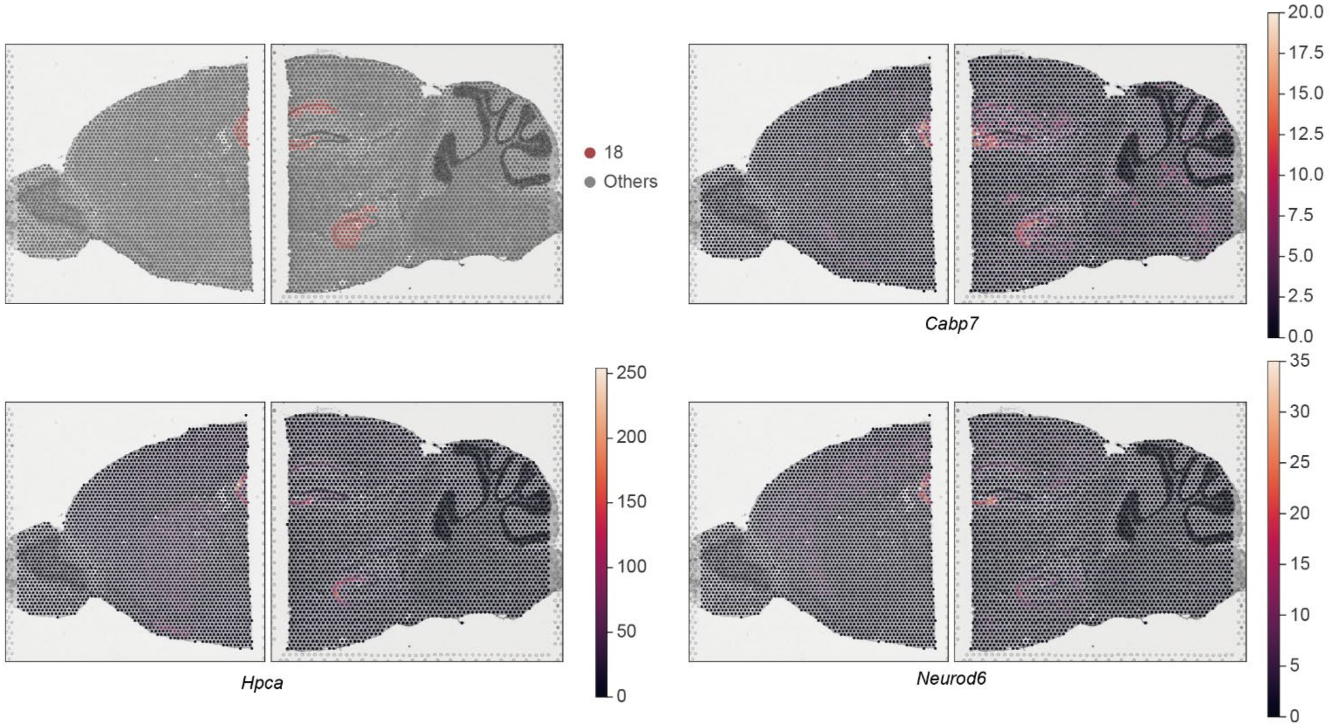

**Supplementary Fig. 23 | The spatial heatmap of differential expressed genes in cortex Ammonis (domain 18) on the two 10x Visium mouse brain sagittal samples.**

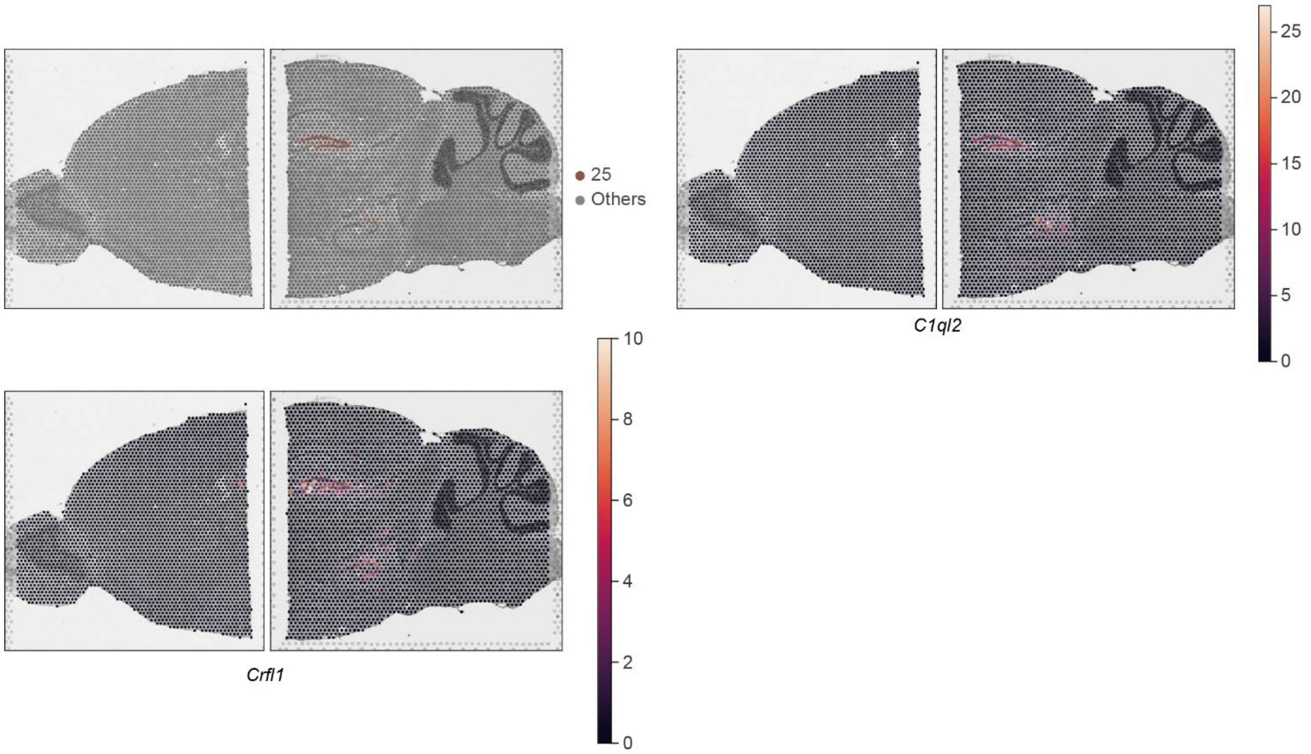

**Supplementary Fig. 24 | The spatial heatmap of differential expressed genes in dentate gyrus (domain 25) on the two 10x Visium mouse brain sagittal samples.**

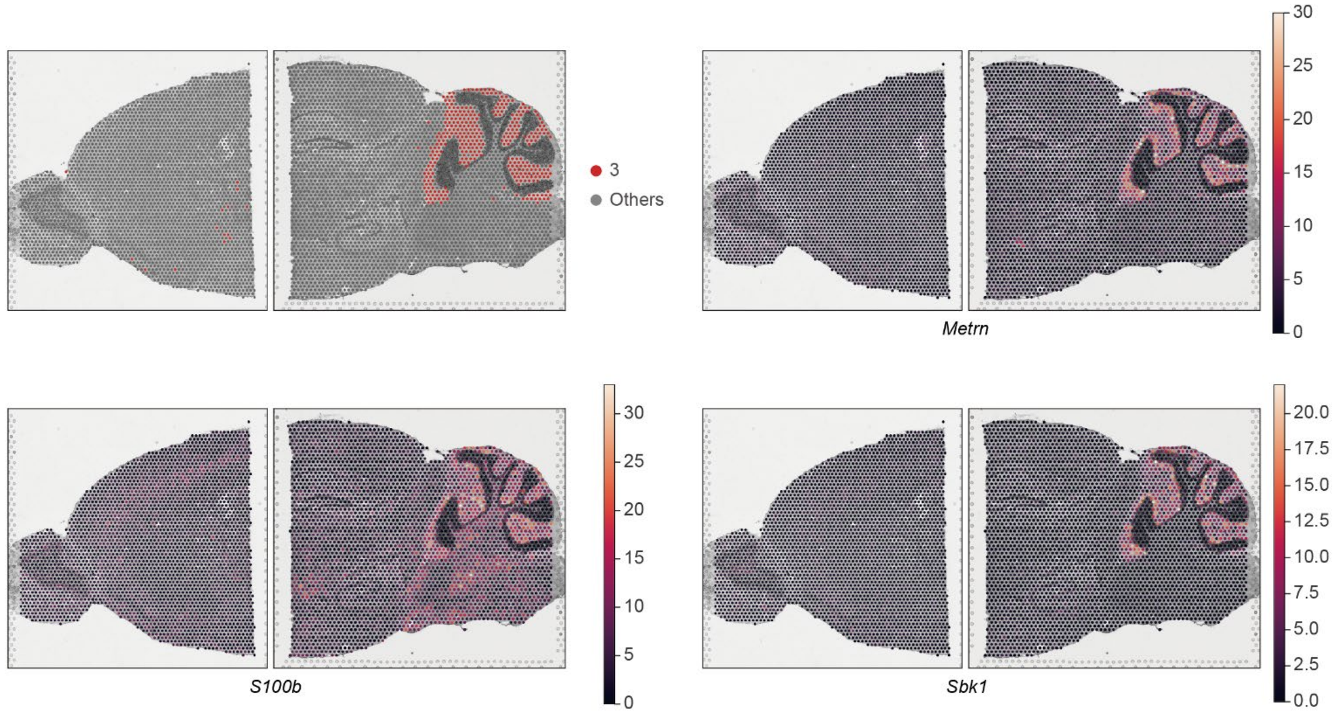

**Supplementary Fig. 25 | The spatial heatmap of differential expressed genes in molecular layer of cerebellar vermis cortex (domain 3) on the two 10x Visium mouse brain sagittal samples.**

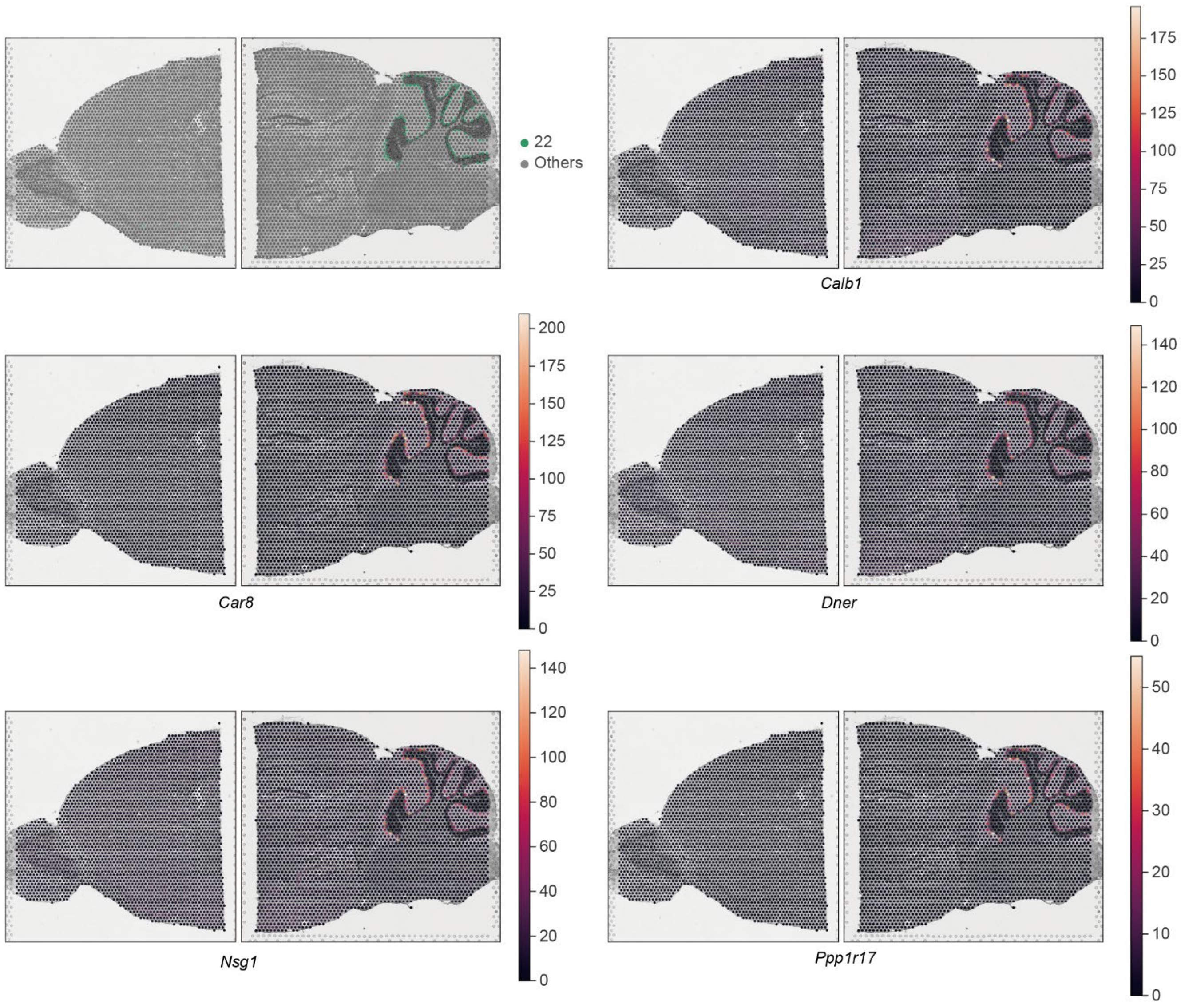

**Supplementary Fig. 26 | The spatial heatmap of differential expressed genes in Purkinje cell layer of cerebellar vermis cortex (domain 22) on the two 10x Visium mouse brain sagittal samples.**

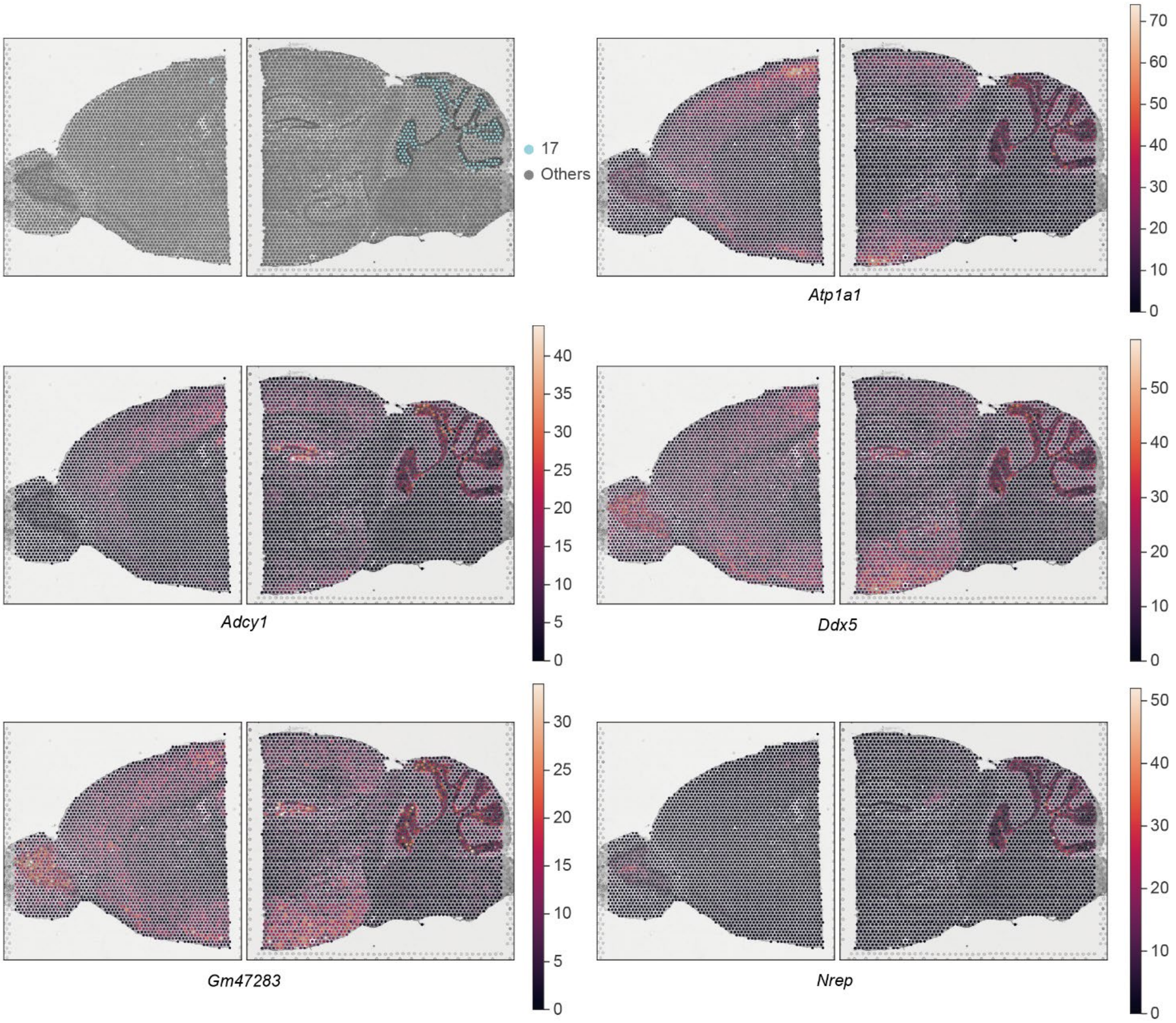

**Supplementary Fig. 27 | The spatial heatmap of differential expressed genes in internal granular layer of cerebellar vermis cortex (domain 17) on the two 10x Visium mouse brain sagittal samples.**

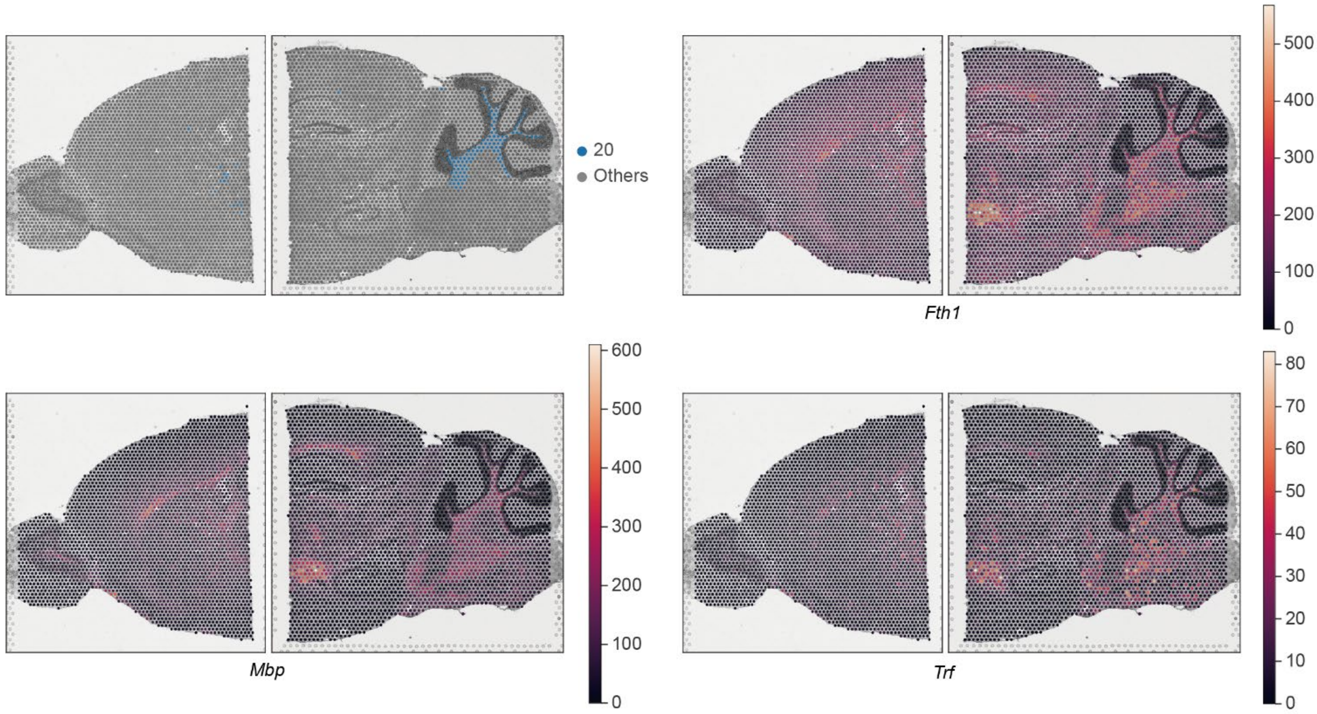

**Supplementary Fig. 28 | The spatial heatmap of differential expressed genes in white matter of cerebellar vermis (domain 20) on the two 10x Visium mouse brain sagittal samples.**

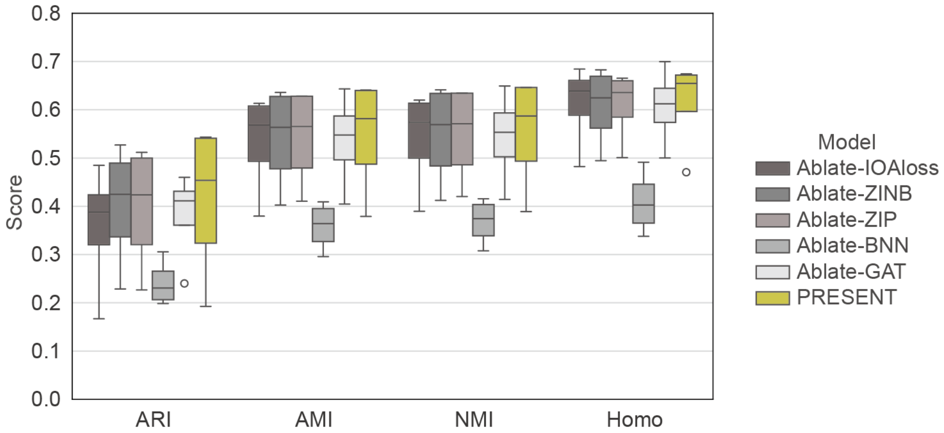

**Supplementary Fig. 29 | Ablation study of PRESENT for the cross-modality representation of MISAR-seq mouse brain samples.** Ablate-IOAloss, Ablate-ZINB, Ablate-ZIP, Ablate-BNN, Ablate-GAT denotes the ablation of inter-omics alignment (IOA) loss in the cross-omics alignment, zero-inflation negative binomial (ZINB) model in the RNA omics-specific decoder, zero-inflation Poisson model in the ATAC omics-specific decoder, Bayesian neural networks (BNN) and graph attention networks (GAT) in the omics-specific encoders, respectively.

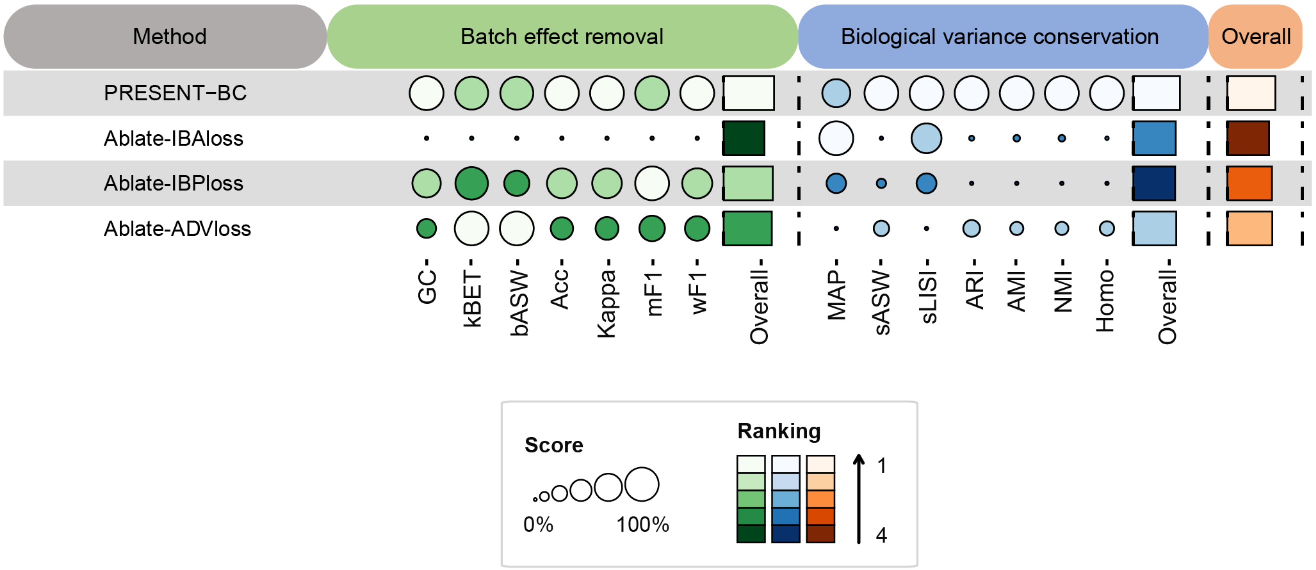

**Supplementary Fig. 30 | Ablation study of PRESENT for the multi-sample integration of MISAR-seq mouse brain samples.** The quantitative evaluation of different ablated models and complete PRESENT on the four MISAR-seq mouse brain samples using 14 metrics divided into two categories, namely batch effect removal and biological variance conservation. The category scores of these two aspects were calculated by averaging the metrics within each category. An overall score for each integration method was computed using a 40/60 weighted mean of the category scores for batch effect removal and biological variance conservation. Ablate-IBAloss, Ablate-IBPloss, Ablate-ADV denotes the ablation of inter-batch alignment (IBA) loss, intra-batch preserving (IBP) loss and batch-adversarial learning strategy in the second integration stage of PRESENT, respectively.

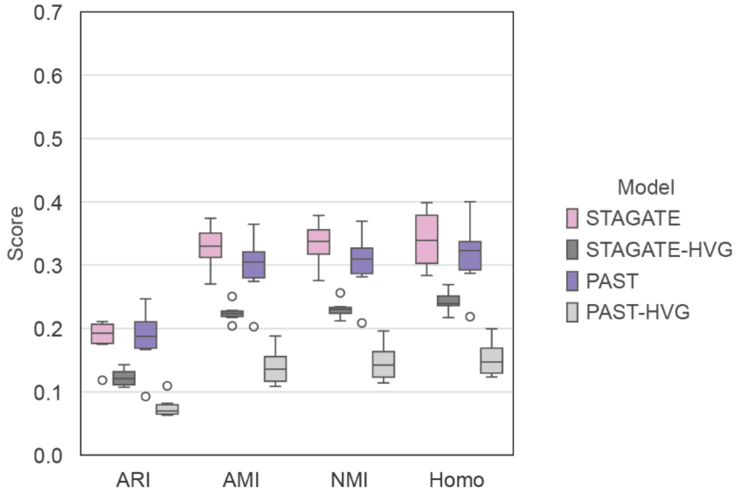

**Supplementary Fig. 31 | Evaluation of the performance of gene selection process on the spatial ATAC mouse embryo samples.** STAGATE and PAST which implemented the highly variable gene (HVG) selection process to obtain 3000 genes are denoted as STAGATE-HVG and PAST-HVG, respectively.

### Supplementary Tables

**Supplementary Table 1 | The gene ontology enrichment results based on the differential expressed genes in Mac1-enriched domain.** BP, biological process; CC, cellular component; MF, molecular function.

| Category | ID | Term | Genes | FDR |
|---|---|---|---|---|
| BP | GO:0048821 | erythrocyte development | Hba-a1, Slc4a1, Alas2, Dmtn, Rhd, Klf1 | 1.290E-08 |
| BP | GO:0061515 | myeloid cell development | Hba-a1, Slc4a1, Alas2, Dmtn, Rhd, Klf1 | 3.586E-07 |
| BP | GO:0030218 | erythrocyte differentiation | Hba-a1, Slc4a1, Alas2, Dmtn, Rhd, Klf1 | 5.480E-06 |
| BP | GO:0034101 | erythrocyte homeostasis | Hba-a1, Slc4a1, Alas2, Dmtn, Rhd, Klf1 | 6.695E-06 |
| BP | GO:0002262 | myeloid cell homeostasis | Hba-a1, Slc4a1, Alas2, Dmtn, Rhd, Klf1 | 2.063E-05 |
| BP | GO:0048872 | homeostasis of number of cells | Hba-a1, Slc4a1, Alas2, Dmtn, Rhd, Kif3a, Klf1 | 3.182E-05 |
| BP | GO:0030099 | myeloid cell differentiation | Hba-a1, Car2, Slc4a1, Alas2, Dmtn, Rhd, Klf1 | 8.713E-05 |
| CC | GO:0005833 | hemoglobin complex | Hbb-bt, Hba-a1, Hbb-bs | 3.631E-05 |
| CC | GO:0031838 | haptoglobin-hemoglobin complex | Hbb-bt, Hba-a1, Hbb-bs | 3.631E-05 |
| MF | GO:0030492 | hemoglobin binding | Hbb-bt, Hbb-bs, Slc4a1 | 3.537E-05 |
| MF | GO:0031720 | haptoglobin binding | Hbb-bt, Hba-a1, Hbb-bs | 3.537E-05 |
| MF | GO:0019825 | oxygen binding | Hbb-bt, Hba-a1, Hbb-bs | 2.381E-04 |
| MF | GO:0004601 | peroxidase activity | Hbb-bt, Hba-a1, Hbb-bs | 1.464E-03 |
| MF | GO:0016684 | oxidoreductase activity, acting on peroxide as acceptor | Hbb-bt, Hba-a1, Hbb-bs | 1.464E-03 |
| MF | GO:0005344 | oxygen carrier activity | Hbb-bt, Hba-a1 | 2.931E-03 |
| MF | GO:0016209 | antioxidant activity | Hbb-bt, Hba-a1, Hbb-bs | 3.043E-03 |
| MF | GO:0030506 | ankyrin binding | Slc4a1, Sptb | 6.065E-03 |
| MF | GO:0030507 | spectrin binding | Dmtn, Kif3a | 9.841E-03 |

**Supplementary Table 2 | The gene ontology enrichment results based on the differential expressed genes in Mac2-enriched domain.** BP, biological process; CC, cellular component; MF, molecular function.

| Category | ID | Term | Genes | FDR |
|---|---|---|---|---|
| BP | GO:0006911 | phagocytosis, engulfment | Igha, Marco, Ighg3, Iglc2 | 7.114E-05 |
| BP | GO:0099024 | plasma membrane invagination | Igha, Marco, Ighg3, Iglc2 | 7.114E-05 |
| BP | GO:0010324 | membrane invagination | Igha, Marco, Ighg3, Iglc2 | 7.114E-05 |
| BP | GO:0006909 | phagocytosis | Igha, Marco, Ighg3, Iglc2 | 4.341E-04 |
| BP | GO:0006959 | humoral immune response | Jchain, Igha, Ighg3, Iglc2 | 5.577E-04 |
| BP | GO:0002449 | lymphocyte mediated immunity | Igha, Tnfrsf1b, Ighg3, Iglc2 | 5.941E-04 |
| BP | GO:0002460 | adaptive immune response based on somatic recombination of immune receptors built from immunoglobulin superfamily domains | Igha, Tnfrsf1b, Ighg3, Iglc2 | 5.941E-04 |
| BP | GO:0042742 | defense response to bacterium | Jchain, Igha, Ighg3, Iglc2 | 5.941E-04 |
| BP | GO:0006910 | phagocytosis, recognition | Igha, Ighg3, Iglc2 | 7.319E-04 |
| BP | GO:0006958 | complement activation, classical pathway | Igha, Ighg3, Iglc2 | 7.492E-04 |
| BP | GO:0002455 | humoral immune response mediated by circulating immunoglobulin | Igha, Ighg3, Iglc2 | 8.082E-04 |
| BP | GO:0050853 | B cell receptor signaling pathway | Igha, Ighg3, Iglc2 | 8.082E-04 |
| BP | GO:0006956 | complement activation | Igha, Ighg3, Iglc2 | 8.222E-04 |
| BP | GO:0050871 | positive regulation of B cell activation | Igha, Ighg3, Iglc2 | 1.255E-03 |
| BP | GO:0016064 | immunoglobulin mediated immune response | Igha, Ighg3, Iglc2 | 1.913E-03 |
| BP | GO:0019724 | B cell mediated immunity | Igha, Ighg3, Iglc2 | 1.913E-03 |
| BP | GO:0050864 | regulation of B cell activation | Igha, Ighg3, Iglc2 | 1.913E-03 |
| BP | GO:0008037 | cell recognition | Igha, Ighg3, Iglc2 | 2.117E-03 |
| BP | GO:0050851 | antigen receptor-mediated signaling pathway | Igha, Ighg3, Iglc2 | 2.309E-03 |
| BP | GO:0002429 | immune response-activating cell surface receptor signaling pathway | Igha, Ighg3, Iglc2 | 2.864E-03 |
| BP | GO:0002757 | immune response-activating signal transduction | Igha, Ighg3, Iglc2 | 2.864E-03 |
| BP | GO:0002768 | immune response-regulating cell surface receptor signaling pathway | Igha, Ighg3, Iglc2 | 2.975E-03 |
| BP | GO:0002764 | immune response-regulating signaling pathway | Igha, Ighg3, Iglc2 | 2.975E-03 |
| BP | GO:0051251 | positive regulation of lymphocyte activation | Igha, Ighg3, Iglc2 | 4.587E-03 |
| BP | GO:0002253 | activation of immune response | Igha, Ighg3, Iglc2 | 4.587E-03 |
| BP | GO:0042113 | B cell activation | Igha, Ighg3, Iglc2 | 4.587E-03 |
| BP | GO:0002696 | positive regulation of leukocyte activation | Igha, Ighg3, Iglc2 | 6.349E-03 |
| BP | GO:0050867 | positive regulation of cell activation | Igha, Ighg3, Iglc2 | 6.697E-03 |
| CC | GO:0042571 | immunoglobulin complex, circulating | Jchain, Igha, Ighg3, Iglc2 | 3.583E-06 |

| | | | | |
|---|---|---|---|---|
| CC | GO:0019814 | immunoglobulin complex | Jchain, Igha, Ighg3, Iglc2 | 3.583E-06 |
| MF | GO:0034987 | immunoglobulin receptor binding | Jchain, Igha, Ighg3, Iglc2 | 1.111E-05 |
| MF | GO:0003823 | antigen binding | Jchain, Igha, Ighg3, Iglc2 | 2.556E-05 |
| MF | GO:0019865 | immunoglobulin binding | Jchain, Fcgrt | 4.961E-04 |
